# Tauroursodeoxycholic acid inhibits *Clostridioides difficile* toxin induced apoptosis

**DOI:** 10.1101/2022.04.13.488266

**Authors:** Colleen M. Pike, John Tam, Roman A. Melnyk, Casey M. Theriot

## Abstract

*C. difficile* infection (CDI) is a highly inflammatory disease mediated by the production of two large toxins that weaken the intestinal epithelium and cause extensive colonic tissue damage. Antibiotic alternative therapies for CDI are urgently needed as current antibiotic regimens prolong the perturbation of the microbiota and lead to high disease recurrence rates. Inflammation is more closely correlated with CDI severity than bacterial burden, thus therapies that target the host response represent a promising yet unexplored strategy for treating CDI. Intestinal bile acids are key regulators of gut physiology that exert cytoprotective roles in cellular stress, inflammation and barrier integrity, yet the dynamics between bile acids and host cellular processes during CDI have not been investigated. Here we show that several bile acids are protective against apoptosis caused by *C. difficile* toxins in Caco-2 cells and that protection is dependent on conjugation. Out of 20 tested bile acids, taurine conjugated ursodeoxycholic acid (TUDCA) was the most potent inhibitor yet unconjugated UDCA did not alter toxin-induced apoptosis. TUDCA treatment decreased expression of genes in lysosome associated and cytokine signaling pathways. TUDCA did not affect *C. difficile* growth or toxin activity *in vitro* whereas UDCA significantly reduced toxin activity in a Vero cell assay and decreased *tcdA* gene expression. These results demonstrate that bile acid conjugation can have profound effects on *C. difficile* as well as the host and that conjugated and unconjugated bile acids may exert different therapeutic mechanisms against CDI.

## Introduction

*Clostridioides difficile* infection (CDI) is a significant public health problem associated with increasing morbidity, mortality, and health-care related costs around the globe (1). CDI is a highly inflammatory disease mediated by the production of two large toxins, TcdA and TcdB, that glycosylate and inactivate Rho and Rac GTPases in colonic epithelial cells. Inactivation of these small GTPases disrupts the actin cytoskeleton, ultimately causing apoptosis and a severe inflammatory response (2, 3). Although antibiotics are the first line of treatment, they are the main risk factor in the development of CDI and 10-25% of successfully treated cases experience recurrent infection (4–8). Recent data has indicated an overly robust immune response is more closely correlated with CDI severity than bacterial burden (9–11), thus rather than modulating antibiotic regimens, therapies that target the host response may be a more effective approach.

Bile acids are key regulators of gastrointestinal physiology and drastically influence the *C. difficile* lifecycle. Primary bile acids cholic acid (CA) and chenodeoxycholic acid (CDCA) are synthesized from cholesterol in the liver and are conjugated with either taurine or glycine. This conjugation step decreases the hydrophobicity of these molecules, facilitating emulsification and absorption of lipids and nutrients from the small intestine into the portal vein (12, 13). Bile acids that are not reabsorbed by the small intestine are further metabolized by members of the gut microbiota. Several microbes deconjugate, dehydroxylate and epimerize primary bile acids into secondary bile acids (deoxycholic acid (DCA), lithocholic acid (LCA) and ursodeoxycholic acid (UDCA)) that are either excreted or reabsorbed and circulated back to the liver where they are reconjugated with taurine or glycine (14, 15). Members of the microbiota can reconjugate bile acids with several other amino acids such as leucine, phenylalanine and tyrosine which further increases the diversity and complexity of the intestinal bile acid pool (16).

Disruption of bile acid metabolism via diet or antibiotics can have detrimental consequences on gut physiology, including increased susceptibility to gastrointestinal and infectious diseases such as CDI (17–19). Prior to antibiotic treatment, microbial-derived secondary bile acids inhibit *C. difficile* growth (18, 20). Protection conferred by secondary bile acids is lost after antibiotics whereas primary bile acids levels increase, supporting *C. difficile* spore germination and outgrowth (21). Aside from directly inhibiting bacterial growth, bile acids can elicit protection against CDI through additional mechanisms. Several bile acids neutralize TcdB’s cytotoxicity by directly binding to the CROP domain, blocking TcdB-receptor binding and preventing entry into host cells (22). Oral administration of bile acids can alter host inflammation. Mice given UDCA prior to *C. difficile* challenge had less colonic edema associated with altered expression of genes in the NF-kB pathway (23). This was the first indication that bile acids may alter the host response during CDI, however it unclear whether this is due to activity against *C. difficile*, the host or both.

Bile acids are potent modulators of intestinal membrane integrity and cell signaling pathways (24), yet little attention has been given to how these metabolites influence the host during CDI. Several bile acids with anti-inflammatory properties and are in clinical trials for inflammatory diseases such as inflammatory bowel disease and CDI (25, 26). Mechanistically, bile acids can maintain intestinal barrier integrity, attenuate apoptosis, cytokine production and improve cell survival (25, 27). We therefore hypothesized that the cytoprotective properties of bile acids can be leveraged against damage caused by *C. difficile* toxins. Here, we screened twenty host- and microbiota-derived bile acids for their ability to inhibit toxin-induced apoptosis in Caco-2 cells. We found that several conjugated bile acids significantly inhibited toxin-induced caspase-3/7 activation with tauroursodeoxycholic acid (TUDCA) being the most potent inhibitor. TUDCA altered the Caco-2 transcriptional profile, decreasing the expression of genes involved in lysosome function and cytokine signaling. The unconjugated form of TUDCA, UDCA, did not attenuate caspase-3/7 activation but significantly decreased *C. difficile* toxin activity and gene expression of *tcdA in vitro*. Conversely, TUDCA had no effect on *C. difficile* pathogenesis *in vitro*. Collectively, our results demonstrate that cytoprotective bile acids elicit can be leveraged against *C. difficile* toxins and that bile acid conjugation can have profound effects on both *C. difficile* and the host.

## Results

### Conjugated bile acids inhibit toxin-induced caspase activation

Apoptosis induced by *C. difficile* toxins causes extensive colonic tissue damage, initiating inflammatory cascades that manifest as the detrimental symptoms of CDI (2, 28). Given that several bile acids have potent anti-apoptotic activity (25), we probed whether bile acids can inhibit toxin-induced apoptosis in human intestinal Caco-2 cells (29). Monolayers of Caco-2 cells were treated with TcdA and TcdB (30 pM) for 8 hr before the addition of 20 individual conjugated and unconjugated bile acids. Bile acids and toxins were added asynchronously to avoid effects that could be a result of direct binding between them (22). The colon contains a higher concentration of unconjugated bile acids (∼200 μM to 1 mM) due to conjugated bile acids being either reabsorbed in the small intestine or deconjugated by the microbiota. We treated Caco-2 cells with two concentrations well below critical micelle concentrations, 50 µM and 500 µM, to observe concentration-dependent effects.

Caspase-3/7 activation and cell viability were measured 24 hr after the addition of bile acids (Fig. 1A). Data are displayed as the ratio of caspase-3/7 activation relative fluorescence units (RFU) to cell viability RFU and relative to the no toxin control. A Student’s t-test was used to identify bile acids that significantly decreased caspase-3/7 activation relative to cells treated with DMSO and toxins. After plotting the mean difference in relative caspase-3/7 activation against -log10(*p* value), we observed a trend between caspase activation and bile acid conjugation (Fig. 1B); only conjugated bile acids decreased toxin induced caspase-3/7 activation whereas GLCA and several unconjugated bile acids exacerbated caspase-3/7 activation. TUDCA was the only bile acid treatment to significantly reduce caspase-3/7 activation at 50 µM (p=0.042). At 500 µM, TUDCA, TDCA, TLCA, TCDCA, GUDCA, and GCDCA significantly decreased caspase-3/7 activation (TUDCA *p*=0.02, TDCA *p*=0.0.011, TLCA *p*=0.006, TCDCA *p*=0.0.008, GUDCA *p*=0.014, GCDCA *p*=0.0.04). Conversely, at 50 µM, GLCA, LCA and DCA augmented caspase-3/7 activation but only GLCA was significant (GLCA *p*=0.03, LCA *p*=0.12, DCA *p*=0.17). GLCA also significantly induced caspase-3/7 activation in the absence of toxins (*p* <0.001; Fig. S1), indicating GLCA is toxic to Caco-2 cells on its own. At 500 µM, one conjugated (GLCA), four unconjugated (DCA, GLCA, LCA, CDCA) and four bile acid derivatives (oxoLCA, oxoDCA, iLCA and iDCA) increased caspase-3/7 activation, with iLCA, LCA, CDCA and GLCA causing a significant increase (iLCA *p*<0.0001, LCA *p*=0.02, CDCA *p*=0.02, GLCA *p*=0.014). Similar to the findings of previous reports (30), several unconjugated bile acids (CDCA, DCA, LCA, GLCA, oxoDCA, isoDCA, isoLCA, and oxoLCA) induced apoptosis on their own in the absence of toxins (Fig. S1). Conjugated bile acids, with the exception of GLCA, did not alter apoptosis in the absence of toxins at either concentration.

**Figure 1.**
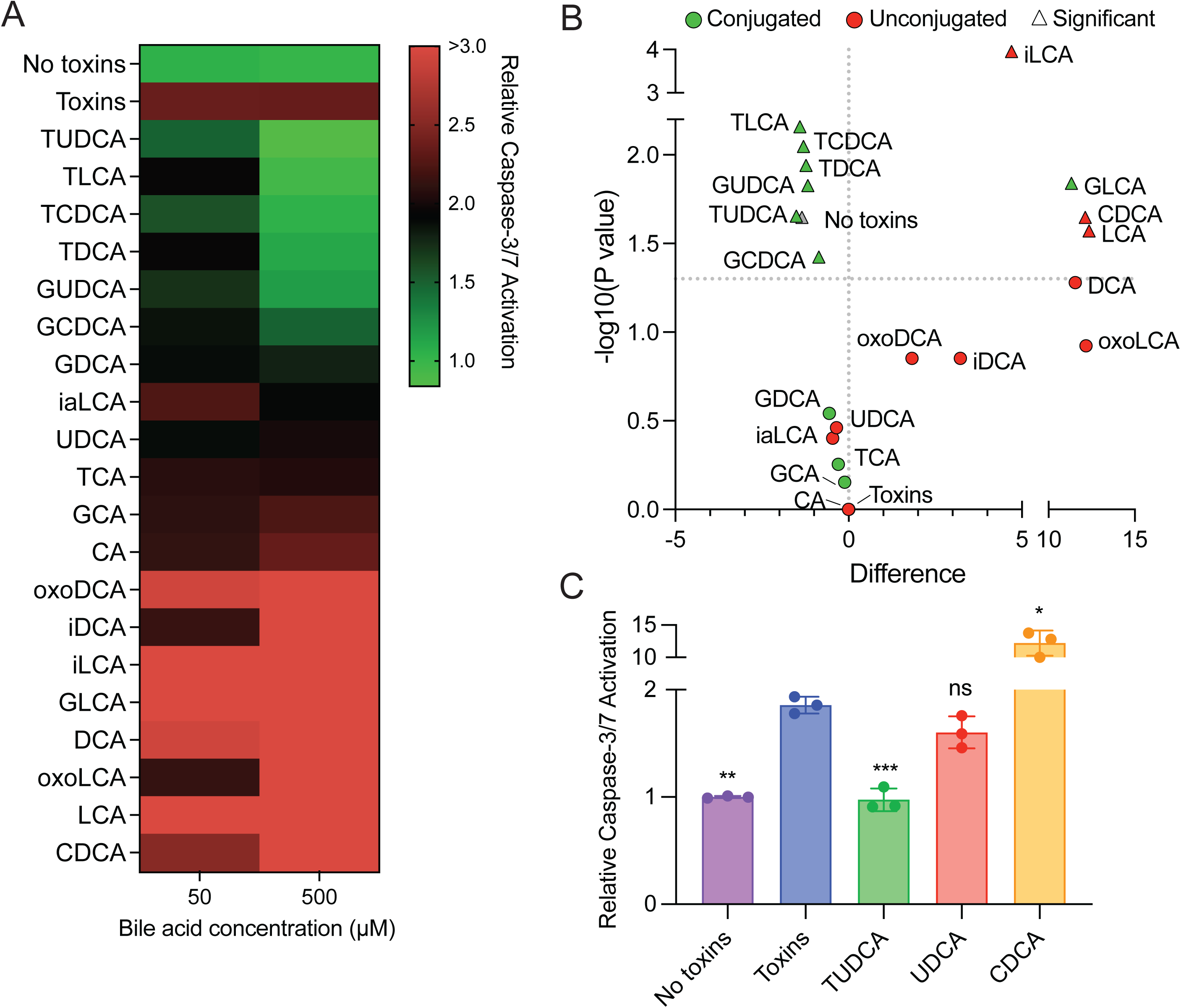
Conjugated bile acids inhibit toxin-induced caspase-3/7 activation in Caco-2. (A) Relative caspase-3/7 activation in Caco-2 cells after treatment with TcdA and TcdB (30 pmol) and the listed bile acids. Data are represented as the ratio of caspase activation RFU to cell viability RFU and relative to no toxins. The colors on the heatmap represent low (green) to high (red) relative caspase activation. (B) The mean difference in relative caspase-3/7 activation between intoxicated cells treated with and without bile acids (500 µM) plotted against -log10(*p* value). Green data points indicate conjugated bile acids and red data points indicate unconjugated bile acids. No toxins and toxins data points are in gray. Triangle shaped data points are significant (p <0.05; Student’s t-test). (C) Relative caspase-3/7 activation in HCT116 cells after treatment with TcdA and TcdB (30 pmol) and the listed bile acids (500 µM) for 48 h. Student’s t-test with Welch’s correction; * *p*<0.05, ** *p*<0.01, *** *p*<0.001. Abbreviations: TUDCA, tauroursodeoxycholic acid; TLCA, taurolithocholic acid; TCDCA, taurochenodeoxycholic acid; TDCA, taurodeoxycholic acid; GUDCA, glycoursodeoxycholic acid; GCDCA, glycochenodeoxycholic acid; GDCA, glycodeoxycholic acid, iaLCA, iso-allo-lithocholic acid; UDCA, ursodeoxycholic acid; TCA, taurocholic acid; GCA, glycocholic acid; CA, cholic acid; oxoDCA, 3-oxo-deoxycholic acid; iDCA, iso-deoxycholic acid; iLCA, iso- lithocholic acid; oxoLCA, 3-oxo-lithocholic acid; GLCA, glycolithocholic acid; DCA, deoxycholic acid; LCA, lithocholic acid; CDCA, chenodeoxycholic acid.

To validate these results, we repeated this assay using another human colorectal carcinoma cell line, HCT116. Using the same experimental set up as described previously, caspase-3/7 activation was measured in intoxicated HCT116 cells treated with TUDCA, UDCA or CDCA (500 µM). Similar to our findings in Caco-2 cells, TUDCA decreased, CDCA exacerbated and UDCA did not change toxin-induced caspase-3/7 activation (no toxins *p*=0.002, TUDCA *p*<0.001, CDCA *p*=0.01, UDCA *p*=0.07; Student’s t test; Fig. 1C). These findings suggest that several bile acids can inhibit toxin induced caspase-3/7 activation and that bile acid conjugation can influence the host response to toxins.

### TUDCA and not UDCA exhibit potent anti-apoptotic activity against *C. difficile* toxins

Of the 20 bile acids tested in our screen, TUDCA was the most inhibitory against toxin-induced apoptosis. Both TUDCA and UDCA have been established as potent inhibitors of apoptosis but their cytoprotective properties have not been leveraged against *C. difficile* toxins (25). We therefore further characterized the inhibitory effects of TUDCA and UDCA against toxins.

Several bile acids directly bind to TcdB, blocking TcdB from binding host cell receptors and being endocytosed (22), however it is unknown whether TUDCA or UDCA can bind to TcdB. We assessed the direct binding between bile acids and TcdB using differential scanning fluorimetry (Fig. 2A). The estimated EC50 (half maximal effective concentration) of TUDCA and UDCA binding was 338 µM and >1000 µM, respectively. Positive control TCDCA bound with an EC50 of 77 µM, while the EC50 of negative control dehydrocholate was >1000µM. To validate these findings, we tested the ability of UDCA and TUDCA to inhibit TcdB-induced cell rounding in IMR90 cells by measuring cell rounding index (CRI). IMR90 cells were treated with both TcdB and increasing concentrations of TUDCA or UDCA for 3 hr before cell rounding was measured. The EC50 of inhibition by TUDCA and UDCA were 684 µM and 662 µM, respectively (Fig. S2). Positive and negative control bile acids TCDCA and dehydrocholate had an EC50 of 75 µM and >400 µM, respectively. TUDCA and UDCA did not bind to TcdA or alter TcdA-induced cell rounding (data not shown).

**Figure 2.**
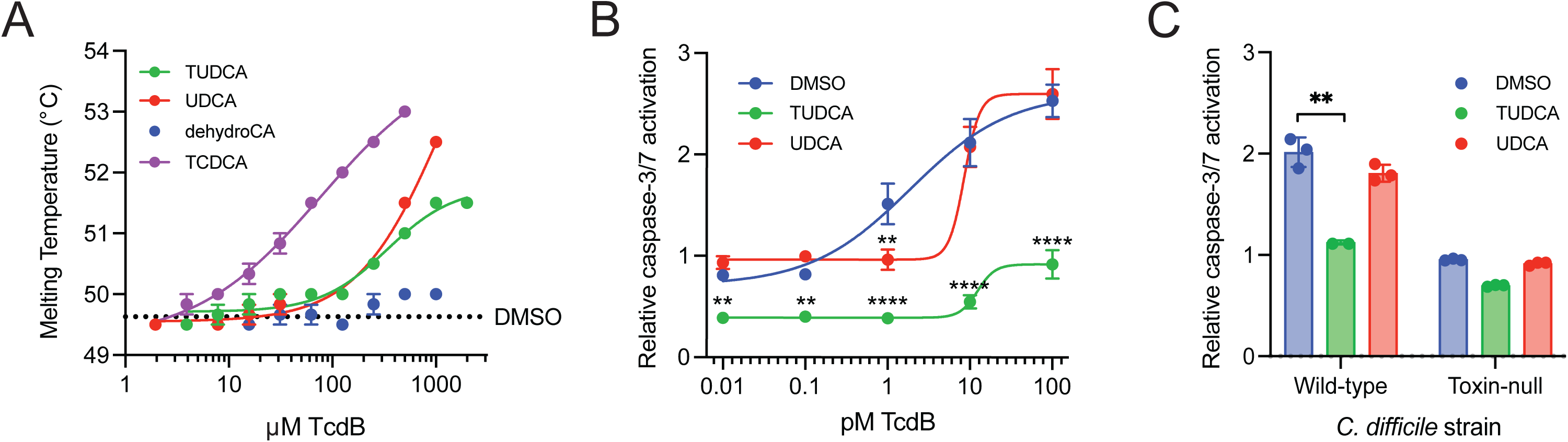
TUDCA and not UDCA exhibits potent anti-apoptotic activity against C. difficile toxins. (A) Titration curves of increasing bile acids by DSF. (B) Relative caspase-3/7 activation of Caco-2 cells treated with 500 μM of the listed bile acids and increasing concentrations of purified TcdB. (C) Relative caspase-3/7 activation of Caco-2 cells treated with 500 μM of the listed bile acids and filtered supernatants from wild-type or toxin-null C. difficile strains (1:500). Bars represent SEM of three separate experiments. Student’s t-test with Welch’s correction; ** p<0.01, *** p<0.001, **** p <0.0001.

We next evaluated the effects of a fixed dose of TUDCA and UDCA (500 µM) against increasing concentrations of TcdB (Fig. 2B). TUDCA significantly reduced caspase-3/7 activation in cells treated with all tested TcdB concentrations (0.01 pM, *p*=0.0.001; 0.1 pM, *p*=0.001; 1 pM. *p*<0.0001; 10 pM, *p*<0.0001; 100 pM, *p*<0.0001; Two-way ANOVA). UDCA was only inhibitory against 1 pM concentration of TcdB (*p*<0.0001). This suggests that UDCA is less potent than its taurine conjugated form. In addition to using recombinant purified toxins to induce apoptosis, we tested the anti-apoptotic activity of TUDCA and UDCA against endogenously produced toxins. Caco-2 cells were treated with filtered supernatants from either a wild-type (*C. difficile* R20291) or toxin-null strain of *C. difficile* (1:500) for 8 hr. DMSO, TUDCA, or UDCA (500 µM) were added and caspase activation was measured 24 hr later (Fig. 2C). Compared to cells treated with DMSO, TUDCA and not UDCA significantly decreased caspase activation (TUDCA, *p*= 0.001; UDCA, *p*=0.058; Student’s t test). The supernatant from the toxin-null strain did not induce apoptosis, indicating that TUDCA’s inhibitory effect was directly against toxins.

### TUDCA alters the transcriptome of toxin-treated Caco-2 cells

To determine how TUDCA alters the host response to toxins, we compared gene expression profiles of untreated, toxin treated, and TUDCA+toxin treated cells (referred to as no toxins, toxins and TUDCA, respectively). RNA isolated from Caco-2 cells was analyzed using the NanoString Host Response panel. We first assessed how toxins altered Caco-2 gene expression relative to no toxins. After processing the raw data using nSolver, 438 transcripts were above threshold level (probe count > 20). 247 genes had a significant fold change in expression (*p* <0.05; Student’s t test) with 169 transcripts being increased and 78 transcripts being decreased (Fig. 3A and 3C, Table S3). Gene set enrichment analysis identified cytokine-mediated signaling, defense response, cell surface receptor signaling, immune system process, and response to stress as enriched gene ontogeny (GO) biological process terms in differentially expressed genes (DEGs) with increased abundance (Fig. 3E, Table S4). Pathway enrichment analysis identified TNF signaling, apoptosis, NOD-like receptor signaling, MAPK signaling and autophagy as enriched KEGG pathways in DEGs with increased abundance (Fig. 3E). This transcriptional profile is in line with previous findings that toxins induce cytotoxic and inflammatory signaling pathways.

**Figure 3.**
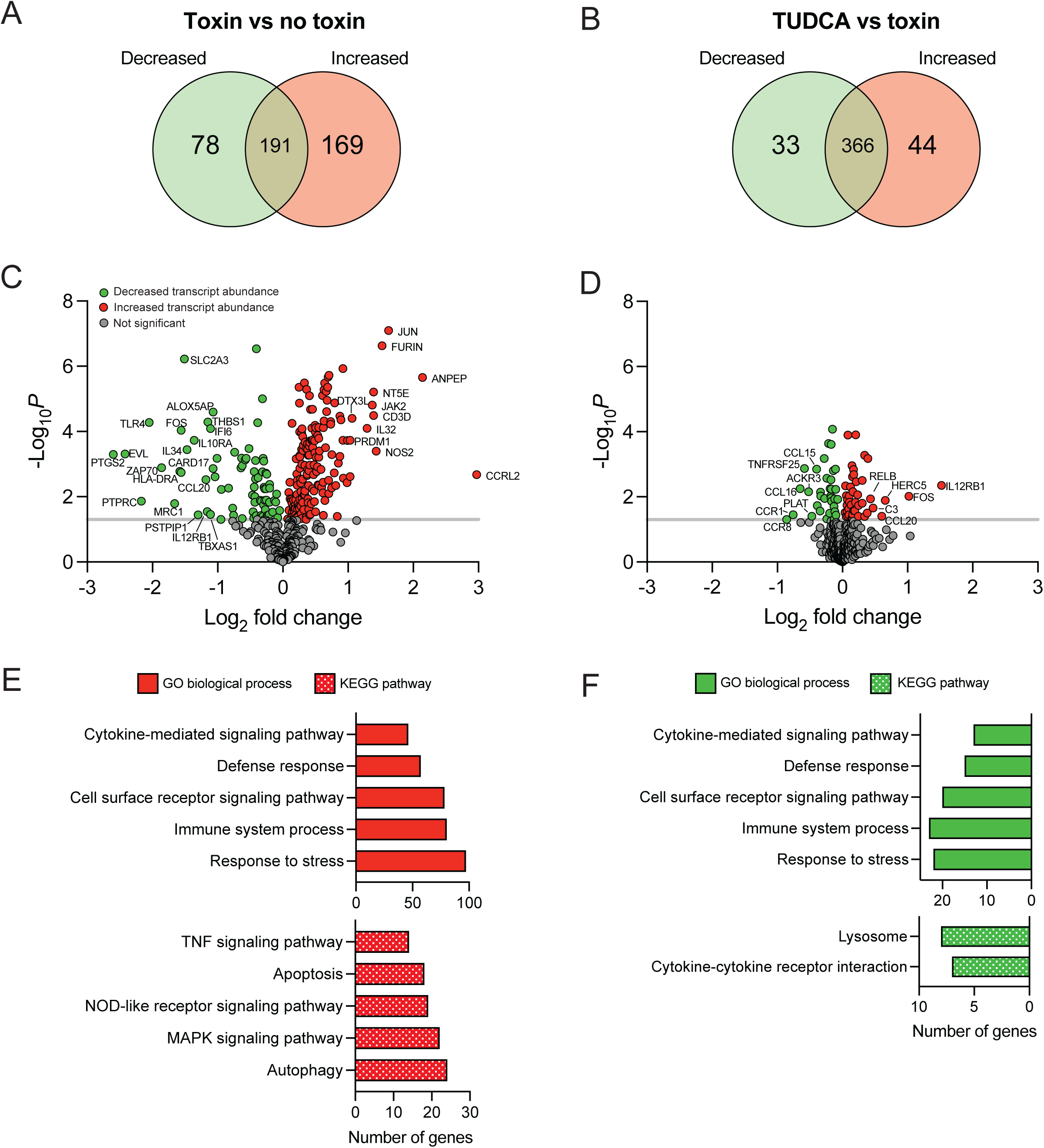
TUDCA alters the transcriptome of toxin-treated Caco-2 cells. (A, B) Venn diagram depicting the significant differentially expressed genes (DEGs) in intoxicated Caco-2 cells treated with (left) toxins relative to no toxins and (right) TUDCA relative to toxins (increased expression, yellow; decreased expression, purple). (C, D) Volcano plots displaying log2 fold change plotted against -log10(*p* value) of DEGs in intoxicated Caco-2 cells treated with (left) toxins relative to no toxins and (right) TUDCA relative to toxins. Red data points highlight genes with increased fold change, green data points highlight genes with decreased fold change, and gray data points highlight genes with insignificant fold changes. (E, F) GO biological terms and KEGG pathways of (left) toxin relative to no toxin DEGs with increased abundance and (right) TUDCA relative to toxin DEGs with decreased abundance (FDR <0.05).

We next compared gene expression profiles of Caco-2 cells treated with TUDCA and toxins relative to toxins alone. nSolver detected 476 transcripts above threshold level (probe count > 20). A Student’s t test identified 77 genes that had a significant fold change in expression, with increased expression in 33 genes and decreased expression in 44 genes (Fig. 3B and 3D, Table S5). However, the fold change were relatively low with only two genes being above a 1-fold change (*FOS* and *IL12RB1*). The GO biological process terms identified among the DEGs with decreased abundance matched the GO terms associated with toxin treatment (Fig. 3F and Table S6), suggesting TUDCA decreased the expression of several genes involved in cell stress and immune signaling pathways induced by toxins. The lysosome and cytokine-cytokine receptor interaction signaling KEGG pathways were enriched in DEGs with decreased abundance (Fig. 3F).

Decreased expression of several genes involved in lysosome function, including vacuolar ATPases *TCIRG1* and *ATP6V0D1,* led us to speculate that TUDCA may alter the pH of endosomal compartments. Acidification of endocytic vesicles is required for the release of the cytotoxic glycosyltransferase domain from the endosome (31). Several small molecules inhibit endosomal acidification thereby preventing pore formation and translocation of toxins into the cytosol (32). To assess the effect of TUDCA and UDCA on endosomal pH, we quantified lysotracker staining in IMR90 cells treated with increasing concentrations of bile acids (Fig. S3). We observed no change in lysotracker staining, indicating that TUDCA does not alter cellular pH and may instead modulate other lysosome associated processes like autophagy.

Several transcripts (*CCR1*, *CCL15*, *CCL16* and *MYC*) were selected for further qRT-PCR analysis to validate the NanoString results (Fig. S4). Relative to toxin-treated cells, TUDCA treatment significantly decreased expression of *CCR1*, *CCL15* and *CCL16* and increased expression of *MYC* (*CCR1*, *p*=0.046; *CCL15*, *p*=0.046; CCL16, *p*=0.041; *MYC*, *p*=0.003; Student’s t test). These findings were similar to the trends observed in the NanoString data.

### Differential effects of TUDCA and UDCA on *C. difficile* pathogenesis *in vitro*

UDCA alters *C. difficile* spore germination, vegetative cell growth and toxin activity *in vitro* but TUDCA’s influence on *C. difficile* pathogenesis *in vitro* has not been investigated (23). To determine the effect of TUDCA on spore germination, *C. difficile* spores were added to BHI containing 1% of the spore germinant, TCA, and either CDCA (0.04%, a potent inhibitor of spore germination), TUDCA or UDCA. Spore germination was measured as the decrease in OD600 measured every 20 min over 1 hr. Cultures grown in TCA had a significant drop in OD600 after 20 min whereas the OD600 of cultures grown in CDCA did not change over the 1 hr period (Fig. 4A). Similar to CDCA, cultures grown in either UDCA and TUDCA did not drop in OD600 over 1 hr, indicating that both bile acids inhibit TCA-induced spore germination.

**Figure 4.**
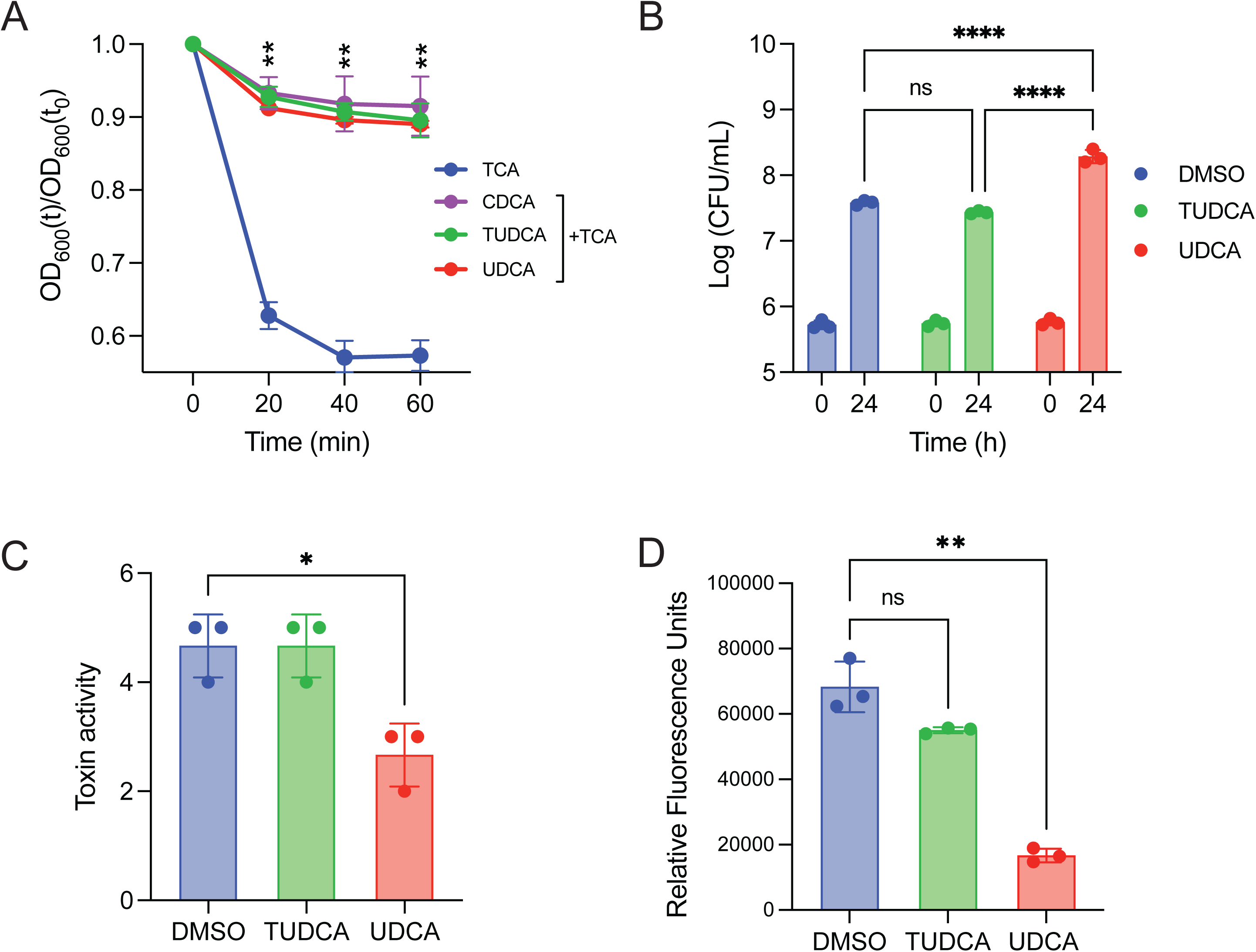
TUDCA does not affect *C. difficile* pathogenesis *in vitro*. (A) *C. difficile* spores were grown in the presence of the spore germinant, TCA, and 500 μM of the listed bile acids. Germination was monitored by plotting the ratio of the OD600 at the listed time points to the OD600 at time zero. (B) Culture aliquots of *C. difficile* taken at 0 and 24 hr were enumerated on TBHI plates to obtain CFU/ml of total vegetative cells and spores. (C) Filtered culture supernatants after 24 hr of growth were used for a Vero cell cytotoxicity assay. Toxin activity is expressed as log10 reciprocal dilution toxin per 100 μl of *C. difficile* culture supernatant. (D) *C. difficile* carrying a P*tcdA-mCherry* reporter plasmid was grown for 24 hr with the listed treatments. Data are displayed as the ratio of fluorescence to absorbance. (B) two-way ANOVA with Sidak’s multiple comparison test, (C and D) Student’s t-test with Welch’s correction. All experiments were performed in triplicate. * p <0.05, ** p < 0.005, **** p <0.0001.

We next tested the ability of TUDCA to alter *C. difficile* growth by enumerating CFU/ml after 24 hr of anaerobic growth in BHI containing either DMSO, TUDCA or UDCA (500 µM; Fig. 4B). Similar to previous findings, UDCA significantly increased *C. difficile* growth (*p* <0.0001; two-way ANOVA with Sidak’s multiple comparison test). TUDCA had no effect on growth (*p* =0.1436). From the same cultures, we measured toxin activity from *C. difficile* supernatant using a Vero cell rounding assay. Supernatants were filtered, serially diluted and added to Vero cells in a 96 well plate. Cell rounding was assessed 24 hr later (Fig. 4C). Compared to the DMSO control, UDCA significantly decreased toxin activity while TUDCA had no effect on toxin activity (UDCA *p*=0.0132, TUDCA, *p* >0.9999; Student’s t test).

Our DSF results revealed that UDCA has an EC50 >1000 µM, thus the decreased toxin activity is not attributed to direct binding and neutralization of TcdB. To determine how UDCA could be altering toxin activity, we tested the effect of UDCA on toxin expression using a fluorescent reporter plasmid. *C. difficile* carrying the pDSW1728 plasmid encoding the *tcdA* promoter upstream of *mCherry* was grown in BHI + thiamphenicol supplemented with DMSO, TUDCA or UDCA (500 µM) for 24 hr. Data are represented as the ratio of fluorescence to OD600 (Fig. 4D). *C. difficile* carrying a plasmid with no P*tcdA-mCherry* cassette was used as a negative control and produced no fluorescence after 24 hr of growth. We observed a significant reduction in normalized fluorescence in *C. difficile* grown in UDCA compared to the DMSO control (*p*=0.004, Student’s t test). No significant reduction in fluorescence was observed in *C. difficile* grown in TUDCA compared to DMSO (*p*= 0.09). These data demonstrate that TUDCA and UDCA exert different effects on *C. difficile* pathogenesis *in vitro*.

## Discussion

Inflammation is suggested to benefit *C. difficile*, as patients with Inflammatory bowel disease are four times more likely to acquire CDI, and a shift in the nutrient availability in the inflamed mouse gut may offer an alternative nutrient source for *C. difficile* (33–35). Immune cell infiltration is necessary for mounting an early defense against CDI (36, 37), however a robust inflammatory response can be detrimental, as prolonged exposure to proinflammatory mediators can exacerbate tissue damage (9, 38, 39). Thus, therapies that target inflammation and the host response during CDI could potentially decrease disease severity and disease recurrence. Bile acids act as an important communication system between the intestinal lumen and the liver, influencing bile acid synthesis, barrier maintenance and inflammatory pathways (24). The *C. difficile* lifecycle is sensitive to different bile acids, yet the dynamics between bile acids and host cellular responses during CDI have been given little attention. We found that several bile acids alleviated toxin-induced caspase-3/7 activation in Caco-2 cells. Similar to previous findings, caspase inhibition was dependent on bile acid conjugation (30, 40). Several taurine and glycine conjugated bile acids were protective whereas unconjugated bile acids exacerbated toxin-induced apoptosis. CDCA, DCA and LCA can activate the intrinsic apoptosis pathway at concentrations below their critical micelle concentrations by inducing oxidate stress and altering mitochondria membrane permeability (41). This could explain why several unconjugated bile acids augmented toxin-induced apoptosis.

It is well established that antibiotics disrupt the gut microbiota, depleting secondary bile acids and increasing susceptibility to CDI (18). Given that secondary bile acids regulate multiple host cell processes in the gut (24), the loss of these key cell signaling molecules is expected to also impact the host response, yet this association in the context of CDI has not been explored. We found that conjugated secondary bile acids (TUDCA, GUDCA, TCDCA, GCDCA, GDCA, TLCA) decrease toxin induced caspase-3/7 activation. This class of bile acid is lost after antibiotic use, thus restoring these protective bile acids could be used as a treatment strategy to attenuate cellular damage and inflammation caused by *C. difficile* toxins.

Of our screen of 20 bile acids, TUDCA was the most inhibitory against toxin-induced apoptosis. UDCA was the only unconjugated bile acid to not increase toxin-induced apoptosis, possibly due to its low hydrophobicity. GUDCA treatment also yielded significant protection, signifying that bile acid conjugation can have varying effects on the host response. Previous work from our group has shown that mice pretreated with an oral gavage of UDCA had an attenuated host inflammatory response in a mouse model of CDI (23). Metabolomics of stool from the same study found that mice treated with UDCA had an almost 5-fold increase in TUDCA compared to UDCA. Mice do not produce glycine conjugated bile acids, thus only an increase in taurine conjugation was observed. This elevated concentration of TUDCA is presumably due to the conjugation of taurine to UDCA by conjugative liver enzymes. Due to the extensive increase in TUDCA, it is unclear which bile acid is responsible for altering the host inflammatory response during CDI. Our results suggest that TUDCA may partly contribute to the protection conferred by UDCA administration. Interestingly, both TUDCA and UDCA have been reported to attenuate apoptosis but only TUDCA was inhibitory against toxin induced caspase-3/7 activation (23). UDCA and TUDCA both modulate mitochondria permeabilization and release of pro-apoptotic signals. Mitochondria permeabilization partly contributes to toxin-induced apoptosis, thus it is unclear why UDCA was not protective (42, 43). UDCA decreased toxin activity and *tcdA* gene expression which is in line with *in vivo* findings that UDCA administration significantly inhibited toxin activity in the mouse cecum on day 4 (23). Taken together, our data suggests administration of UDCA may offer a dual method of protection against CDI, with TUDCA modulating the host cell response and UDCA targeting toxin production.

We used NanoString to understand how toxins and TUDCA alter Caco-2 cell gene expression. Relative to untreated cells, toxin treated cells caused significant fold changes in 247 genes. In DEGs with increased abundance, autophagy, apoptosis, TNF signaling and NOD-like receptor signaling KEGG pathways were enriched. The addition of TUDCA to intoxicated Caco-2 cells resulted in significant fold changes in only 77 genes. This relatively low number of DEGs may indicate that TUDCA acts at the protein level rather than the transcriptional level to inhibit toxin-induced apoptosis. TUDCA has endoplasmic reticulum chaperone activity and can modify Rac1 GTPase *in vitro*, corroborating this hypothesis (44, 45). Of the DEGs with decreased abundance, five genes were in the cytokine-cytokine receptor interaction KEGG pathway (*CCR1*, *CCL15*, *CCL16*, *IL1R2*, *IL17RA*), several of which have been implicated in inflammatory intestinal diseases. CCL15 and CCL16 are the ligands of receptor CCR1; receptor binding mediates chemotaxis of neutrophils, monocytes and lymphocytes (46). CCR1^-/-^ mice had less ileal epithelial cell damage and immune cell infiltration in response to TcdA (47). IL-17RA is the receptor for the proinflammatory cytokine IL-17. The role of IL-17RA signaling in inflammation has produced conflicting results, as IL-17RA^-/-^ mice were protected from trinitrobenzenesulfonic induced colitis, whereas mice given a neutralizing antibody against IL-17RA experienced exacerbated inflammation in response to DSS-induced colitis (48, 49). TUDCA is anti-inflammatory against several diseases, but has not been previously shown to alter expression of this set of genes. TUDCA treatment also decreased expression of eight lysosome-associated genes, including *LAMP1*. While TUDCA’s direct role in lysosome function has not been explored, TUDCA has been shown to modulate autophagy. Fusion between lysosomes and autophagosomes is a key step in the progression of autophagy. Two studies reported that TUDCA decreased autophagy and protein levels of LAMP1 in both TGF-β1 treated fibroblasts and in a mouse model of Parkinson’s Disease (50, 51). TUDCA may attenuate autophagy caused by *C. difficile* toxins, although further proteomic and phenotypic analyses are needed to pinpoint the direct mechanism between TUDCA and these pathways.

While this study is limited in that bile acid mediated protection was only studied in Caco-2 cells, several pieces of evidence suggest that TUDCA has therapeutic applications against inflammatory diseases in both mouse models and humans. UDCA is currently in a phase 4 clinical trial for recurrent CDI and TUDCA is in a phase 1 trial for ulcerative colitis. Several studies have shown that in dextran sodium sulfate-induced mouse models of colitis, TUDCA decreases inflammation and improves survival rates (52–55). Mechanistically, these same studies found that TUDCA decreased apoptosis, the unfolded protein response and production of inflammatory markers.

A recent preprint corroborated our findings that conjugated bile acids are protective against cytotoxic compounds (56). Unconjugated bile acids increased the permeability of Caco-2 monolayers whereas conjugated bile acids prevented this disruption of membrane integrity by forming micelles around unconjugated bile acids. To increase levels of conjugated bile acids in the intestine, rats fed a choline-deficient, high-fat diet were given the bile salt hydrolase inhibitor AAA-10. AAA-10 rats had elevated cecal TUDCA which was associated with improved barrier integrity and decreased intestinal and hepatic inflammation. A similar strategy could be applied to mouse models of CDI to assess the protective role of conjugated bile acids *in vivo*. Manipulating the bile acid pool is currently being explored as a therapeutic strategy for CDI. Our findings demonstrate that alterations to the bile acid pool, specifically deconjugation, may have detrimental effects on the host. While deconjugation is the first step to produce secondary bile acids that are inhibitory against *C. difficile* growth, an increase in unconjugated bile acids could potentially damage intestinal epithelial cell membranes or activate proapoptotic pathways, leading to exacerbated cell death and inflammation. When designing therapies that alter the bile acid pool, careful consideration is needed to avoid the production of toxic concentrations of unconjugated bile acids in the intestine.

## Materials and Methods

### Stains, Cell Lines and Reagents

*C. difficile* strains R20291 and R20291Δ*tcdR* were used in this study (35). *C. difficile* spores were maintained on brain heart infusion (BHI) medium supplemented with 100 mg/liter L-cysteine and 0.1% taurocholate (Sigma-Aldrich, T4009). Cultures were started by inoculating a single colony from the plate into BHI liquid medium supplemented with 100 mg/liter L-cysteine and grown anaerobically. Caco-2 (ATCC, HTB-37), HCT116 (ATCC, CCL-247), IM90 (ATCC, CCL-186) and Vero cells (ATCC, CCL-81) were cultured in DMEM supplemented with 2 mm L-glutamine and 10% FBS and incubated in 5% CO2 at 37 °C. CellEvent™ Caspase-3/7 Green Detection Reagent (C10723) and PrestoBlue™ Cell Viability Reagent (A13261) were purchased from ThermoFisher Scientific. All bile acids were dissolved in DMSO to a final concentration of 40 mM. Vendors and catalog numbers of bile acids used in this study are listed in Table S1. Antitoxin for the Vero cell cytotoxicity assay was purchased from TechLabs (T5000). The plasmid pDSW1728-P*tcdA*::*mCherry* was a kind gift from Dr. Craig Ellermeier (University of Iowa). All strains, plasmids and primers used in this study are listed in Table S2.

### Protein Expression and Purification

Plasmid pHis1522 encoding his-tagged TcdB was a kind gift from Hanping Feng. Expression and isolation of recombinant TcdB was as described by Yang et al (57). Briefly, transformed *Bacillus megaterium* was inoculated into LB containing tetracycline and grown to an A600 of 1.6, followed by overnight xylose induction at 30°C. Bacterial pellets were collected, resuspended with 20 mM Tris pH 8/0.1 M NaCl, and passed twice through an EmulsiFlex C3 microfluidizer (Avestin, Ottawa, ON) at 15,000 psi. The resulting lysate was clarified by centrifuging at 18,000 ×g for 20 min. TcdB was purified by nickel affinity chromatography followed by anion exchange chromatography using HisTrap FF Crude and HiTrap Q columns (Cytiva), respectively. Fractions containing TcdB or TcdA were verified by SDS-PAGE, then pooled and diafiltered with a 100,000 MWCO ultrafiltration device (Corning) into 20 mM Tris PH 7.5/150 mM NaCl. Finally, glycerol was added to 5% v/v, the protein concentration was estimated by A280, divided into single use aliquots, and stored at - 80°C.

### Caspase-3/7 activation assay

Activation of Caspases 3 and 7 was assessed using CellEvent Caspase-3/7 Green Detection Reagent according to the manufacturer’s instructions. Caco-2 cells or HCT116 cells were seeded in 96 well plates for 7 days or until confluent. DMEM containing recombinantly purified TcdA and TcdB (30 pmol) or filtered *C. difficile* supernatants (1:500) were added to the wells and the plates were incubated for 8 hr at 37 °C before the media was replaced with fresh DMEM containing bile acids. After 24 hr of incubation, the media was replaced with 1X PBS containing 5% FBS and 2µM of CellEvent™ Caspase-3/7 Green Detection Reagent. The plates were incubated for 1 hr at 37 °C and green fluorescence was detected at excitation/emission wavelengths of 485/530 nm using a ThermoFisher Fluoroskan Plate Reader. PrestoBlue™ Cell Viability Reagent was then added to the wells and incubated for 30 min at 37 °C. Red fluorescence was detected at excitation/emission wavelengths of 550/610 nm. Data are displayed as the ratio of caspase activation RFU to to cell viability RFU and normalized to the DMSO control.

### Differential Scanning Fluorometry

DSF was performed in a similar manner as described previously (58). TcdB protein was diluted to 0.05 µg/µL using phosphate buffer (100 mM KPO4, 150 mM NaCl, pH 7) containing 5X SYPRO Orange (Invitrogen S6650), and a serial dilution of test compound. A Biorad CFX96 qRT-PCR thermocycler was used to establish a temperature gradient from 42°C to 65°C in 0.5°C increments, while simultaneously recording the increase in SYPRO Orange fluorescence as a consequence of binding to hydrophobic regions exposed on unfolded proteins. The Bio-Rad CFX Manager 3.1 software was used to integrate the fluorescence curves to calculate the melting point.

### Arrayscan high content imaging

IMR90 cells were grown in EMEM (Wisent) supplemented with 10% FBS and penicillin-streptomycin (complete EMEM) and were seeded in 96 well Cellbind plates (Corning) at a density of 3,000 cells/well. After 48 h, the media was exchanged with serum free EMEM (SFM) containing 1 µM Celltracker Orange CMRA (Invitrogen C34551). After 60 min, excess dye was removed by media exchange with 90 μL SFM. An Agilent Bravo liquid handler was used to deliver 0.4 μL of compound from the compound plate to the cell plate, immediately followed by 10 μL of 5 pM TcdB (diluted in SFM), representing a final concentration of toxin (0.5 pM) previously established as EC99 levels of cytopathology. The cell plates were returned to the incubator for 3.5 hr before imaging. Celltracker-labelled cells were evaluated on a Cellomics ArrayScan VTI HCS reader (Thermo Scientific, Waltham, MA) using the Target Acquisition mode, a 10× objective and a sample rate of at least 150 objects per well. After recording all image data, the cell rounding and shrinking effects of TcdB intoxication were calculated using the cell rounding index (CRI), a multiple of the length to width ratio (LWR) and area parameters. The percent inhibition was calculated as the ratio between the sample well and the average toxin-untreated controls after subtracting the average DMSO control values. Dose response curves were created and evaluated using Prism software (Graphpad Software, La Jolla, Ca).

### RNA extraction from Caco-2 cells

RNA was extracted from Caco-2 cells grown in 24 well plates for 21 days using the PureLink RNA Mini kit (Thermo Fisher, 12183025) following the manufacturer’s protocol. The RNA was treated with Turbo DNase (Thermo Fisher, AM2239) to remove genomic DNA contamination. RNA was then column purified according to the manufacturer’s instructions (Zymo, R1019) and RNA integrity was analyzed via qubit and nanodrop.

### NanoString analysis

RNA from Caco-2 cells was submitted to the DELTA Translational Recharge Center at the University of North Carolina at Chapel Hill for quantification of transcripts via NanoString technology. The nCounter Human Host Response gene expression panel was purchased from NanoString Technologies (Seattle, WA). The raw data were normalized to six internal reference genes using nSolver software following the manufacturer’s instructions (NanoString Technologies). Log2 fold changes and significant DEGs were identified using nSolver Advanced analysis software. Enriched GO biological terms and KEGG pathways were identified using ShinyGO with a FDR cut off of 0.05 (59).

### Quantitative reverse transcription PCR

RNA from Caco-2 cells was used as the template in reverse transcription reactions using the High-Capacity cDNA Reverse Transcription Kit (ThermoFisher 4368814) following the manufacturer’s protocol. The resulting cDNA was diluted 1:5 in deionized water and used as the template for quantitative PCR with the SsoAdvanced Universal SYBR Green Supermix (Bio-Rad). For relative quantification, the ΔΔCt method was used to normalize *CCR1*, *CCL15*, *CCL16 and MYC* genes to that of the internal control, *GAPDH*.

### Lysotracker assay

Endosomal pH neutralization was assayed as described by Slater et al. (60); IMR90 cells in complete EMEM were plated at 14,000 cells/well (∼95% confluency). After 24 hr the media was changed to serum free media for 60 min, then bile acid was added and incubated at 37°C for 2 hr. Lysotracker red DND-99 (Life Technologies) was added to 0.1 μM, and incubated for 30 min. Excess dye was removed by media change and the fluorescence at ex/em 574/594 was measured on a Biotek Neo plate reader (Agilent Technologies).

### Spore preparation

Spores were prepared as in Edwards and McBride (61). Mid-log phase cultures were spread onto 70:30 agar plates and incubated at 37 °C for 4 days. The bacterial lawns were scraped off and suspended in 10 mL sterile PBS, mixed 1:1 with 96% ethanol, vortexed for 30 sec, and incubated at room temperature for 1 hr. The suspension was centrifuged at 3000 rpm for 10 min. The pellet was suspended in 10 ml fresh sterile PBS, and centrifuged again; this was repeated twice. The final pellet was suspended in 1 ml PBS and serial dilutions were plated on BHI agar with 0.1% of taurocholate for spore CFU enumeration.

### *C. difficile* in vitro spore germination assay

This assay was performed as previously described (62). Purified spores were enumerated and tested for purity before use and were subjected to heat treatment (65°C for 20 min) to eliminate any vegetative cells. UDCA and TUDCA were dissolved in DMSO, passed into the anaerobic chamber, and added to BHI broth containing 0.1% TCA. CDCA (0.04%), a known inhibitor of TCA-mediated spore germination, was used as a negative control. The spore stock was resuspended in BHI broth containing bile acids to an OD600 of 0.3. The OD600 was measured at 0, 20, 40 and 60 min using a Tecan Infinite F200 Pro plate reader. Data are plotted as ratios of the OD600 at the indicated time points to the OD600 at time zero against time.

### C. difficile growth assay

*C. difficile* R20291 was started from a single colony and grown anaerobically in 5 ml of BHI overnight, subcultured 1:5 in BHI, and grown to mid-log phase (OD, 0.3 to 0.5). Cultures were then diluted to an OD of 0.01 in fresh BHI medium supplemented with either DMSO, UDCA or TUDCA (500 µM) and incubated for 24 hr. At 0 and 24 hr time points, culture aliquots were enumerated on BHI plates to obtain total CFU per milliliter.

### *C. difficile in vitro* Vero cell cytotoxicity assay

Culture supernatants from the anaerobic growth assay were filtered through a 0.22 micron filter and 10-fold dilutions, to a maximum of 10^−6^, were generated. Sample dilutions were incubated 1:1 with 1X PBS (for all dilutions) or antitoxin (performed for 10^−1^ and 10^−4^ dilutions only) for 40 min at room temperature. Following incubation, these mixtures were added to the Vero cells seeded in 96 well plates, and the plates were incubated overnight at 37°C. Vero cells were viewed under 200× magnification for rounding after overnight incubation. The cytotoxic titer was defined as the reciprocal of the highest dilution that produced rounding in 80% of Vero cells for each sample. Vero cells treated with purified *C. difficile* TcdA and TcdB (List Biological Labs, 152C and 155C) and antitoxin were used as controls.

### mCherry fluorescence reporter assay

R20291 strains carrying either pDSW1728-P*tcdA*::*mCherry* or pDSW1728 were grown anaerobically on BHI agar + 10 μg/ml thiamphenicol. A single colony was inoculated into a 5 ml culture of BHI broth + 10 μg/ml thiamphenicol and grown overnight at 37°C. The overnight culture was diluted 1:100 into a fresh 3 ml culture of BHI broth + 3 μg/ml thiamphenicol + either DMSO UDCA or TUDCA (500 µM) and grown at 37°C. After 24 hr of growth, a 500-μl aliquot of culture was mixed with 120 μl of a 5X fixation cocktail: 100 μl of 16% paraformaldehyde aqueous solution and 20 μl of 1 M NaPO4 buffer (pH 7.4). The sample was incubated at room temperature for 30 min, removed from the anaerobic chamber and incubated in the dark for 30 min on ice. The fixed cells were washed three times with 1X PBS, resuspended in 30 μl of PBS and left in the dark at 4°C overnight to allow for chromophore maturation. Red fluorescence was measured at excitation/emission wavelengths of 550/610 nm using a Tecan Infinite F200 Pro plate reader. Absorbance at 600 nm was measured on the same plate reader. Data are displayed as the ratio of fluorescence to absorbance.

### Statistical analysis

All statistical tests were performed in GraphPad Prism 8. Statistical significance was set at a *p* value of <0.05 for all analyses. A Student’s t test was used to test for significance in the apoptosis screen, mCherry fluorescence assay, and *C. difficile* germination assays. A two-way ANOVA was used to test for significance in the *C. difficile* growth assay. For Nanostrings analysis, a Student’s t test was used to calculate significance in gene expression fold changes.

## Acknowledgements

C.M.P. and C.M.T. are funded by the National Institute of General Medical Sciences of the National Institutes of Health under award number R35GM119438. This project was also funded by an intramural grant from the North Carolina State University College of Veterinary Medicine.

## Abbreviations

TCA: taurocholic acid
GCA: glycocholic acid
CA: cholic acid
TCDCA: taurochenodeoxycholic acid
GCDCA: glycochenodeoxycholic acid
CDCA: chenodeoxycholic acid
TDCA: taurodeoxycholic acid
GDCA: glycodeoxycholic acid
DCA: deoxycholic acid
oxoDCA: 3-oxo-deoxycholic acid
iDCA: iso-deoxycholic acid
TLCA: taurolithocholic acid
GLCA: glycolithocholic acid
LCA: lithocholic acid
oxoLCA: 3-oxo-lithocholic acid
iLCA: iso-lithocholic acid
iaLCA: iso-allo-lithocholic acid
TUDCA: tauroursodeoxycholic acid
GUDCA: glycoursodeoxycholic acid
UDCA: ursodeoxycholic acid

---

**Supplemental Figure 1.**
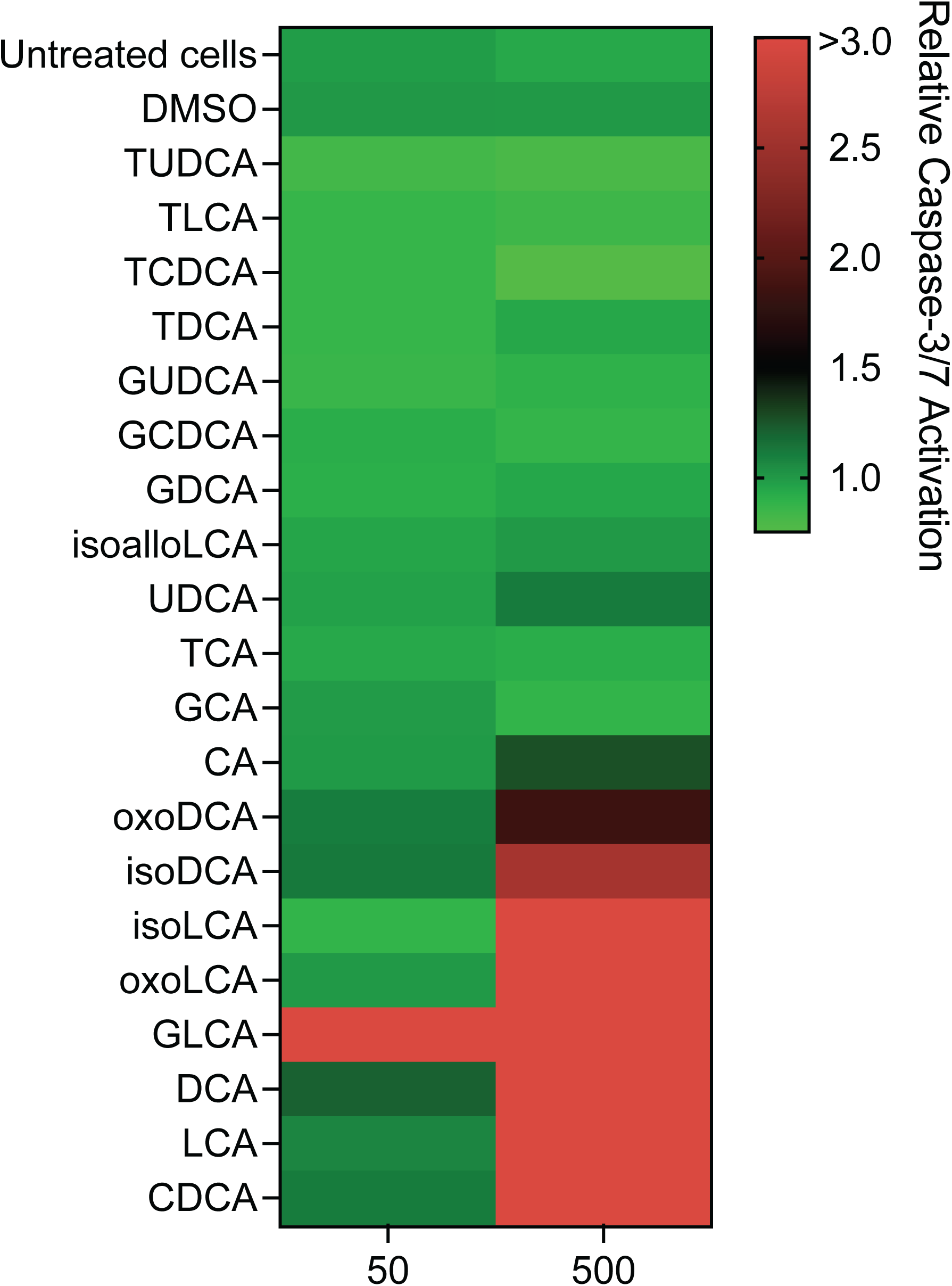
Effect of bile acids on Caco-2 apoptosis without the addition of toxins. Relative caspase-3/7 activation in Caco-2 cells 24 hr after treatment with the listed bile acids. Data are represented as the ratio of caspase activation RFU to cell viability RFU and relative to DMSO. The colors on the heatmap represent low (green) to high (red) relative caspase activation.

**Supplemental Figure 2.**
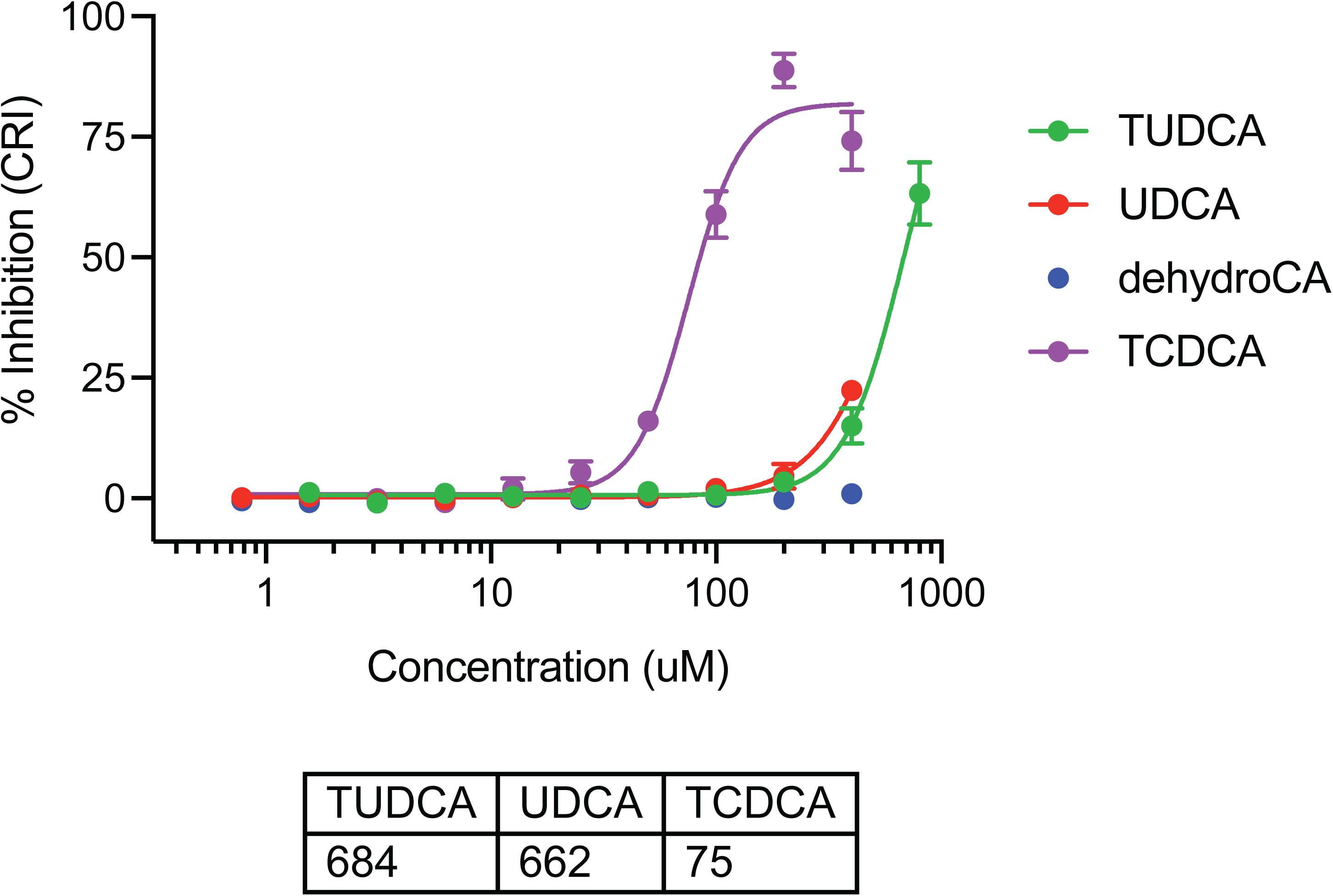
Intoxication of human IMR90 cells measured by cell rounding index (CRI). The EC50s of inhibition by TUDCA and UDCA were 684 µM and 662 µM, respectively. Positive and negative control bile acids TCDCA and dehydrocholate had an EC50s of 75 µM and >400 µM, respectively. Bars represent SEM of three separate experiments.

**Supplemental Figure 3.**
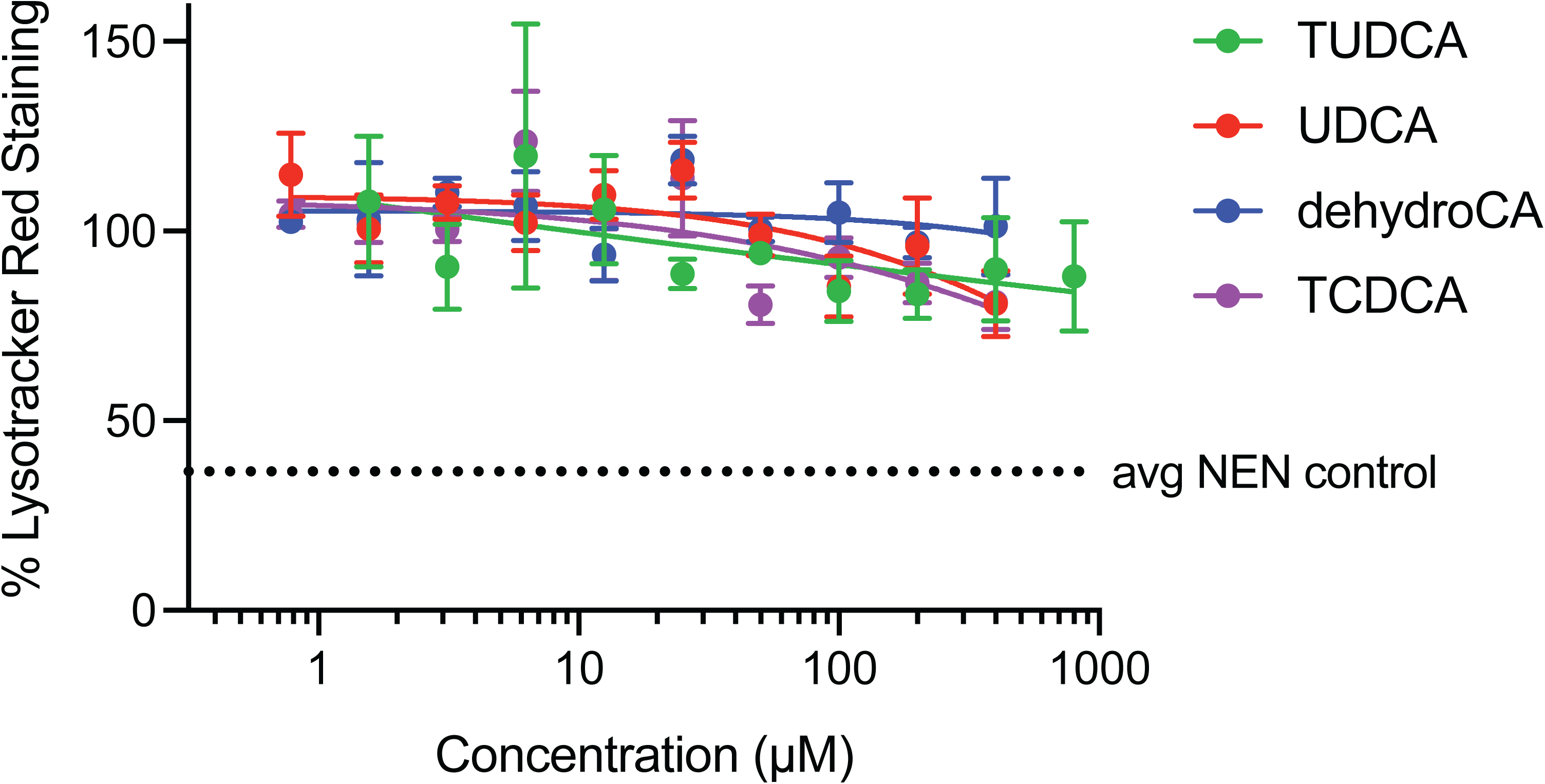
TUDCA and UDCA did not affect lysosomal pH of IMR-90 cells, with EC50s >400 µM. Bars represent SEM of three separate experiments. The dashed line represents % lysosomal staining for control compound Niclosamide (ethanolamine salt form; 4 µM).

**Supplemental Figure 4.**
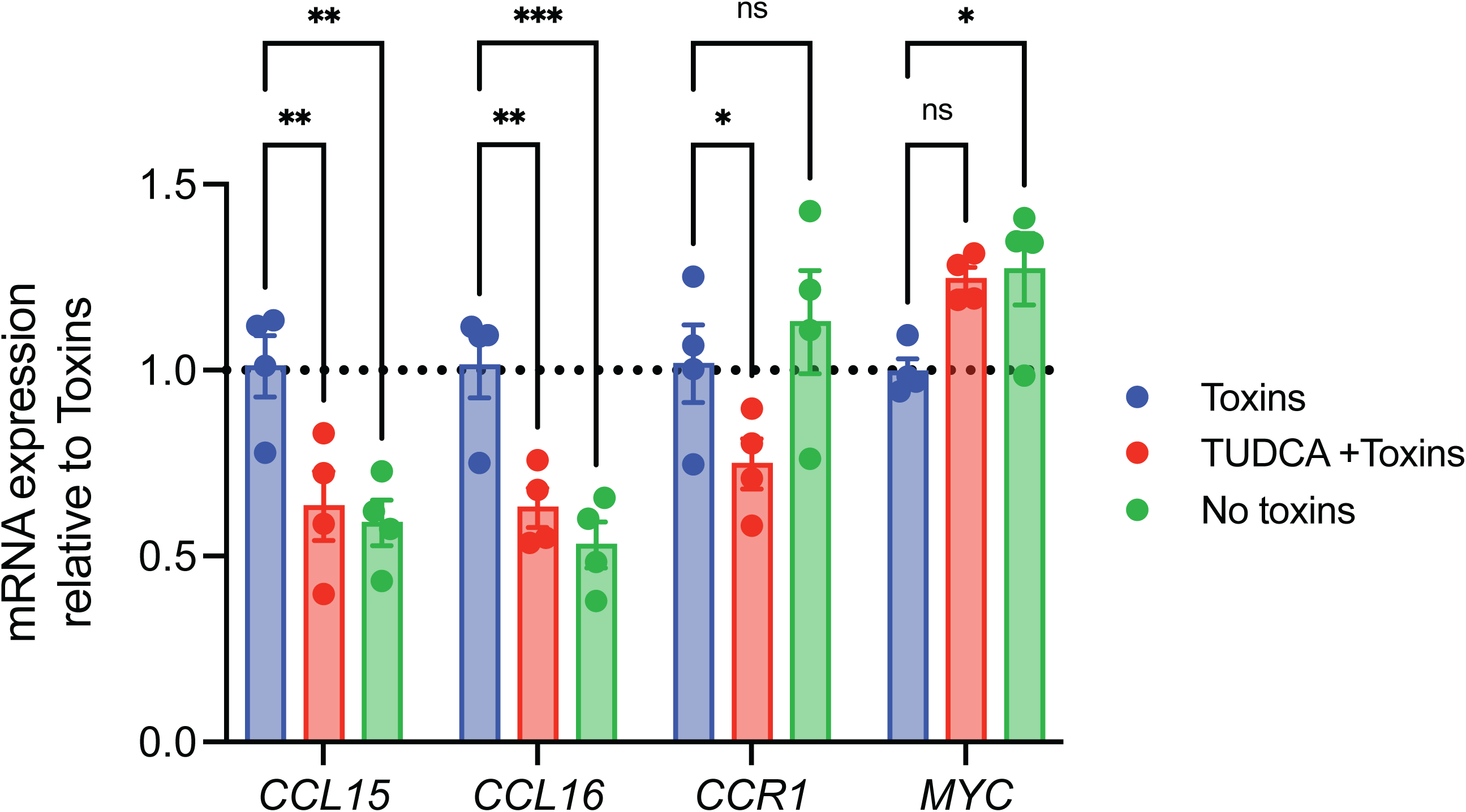
Validation of differentially expressed Caco-2 transcripts by qRT-PCR. Expression was quantified from cDNA generated from RNA isolated from Caco-2 cells. Each point represents a biological replicate. All data are presented as the mean and error bars indicate the sd. * p <0.05, ** p < 0.005, *** p <0.0005; Student’s t test; n=4.

**Supplemental Table 1.**
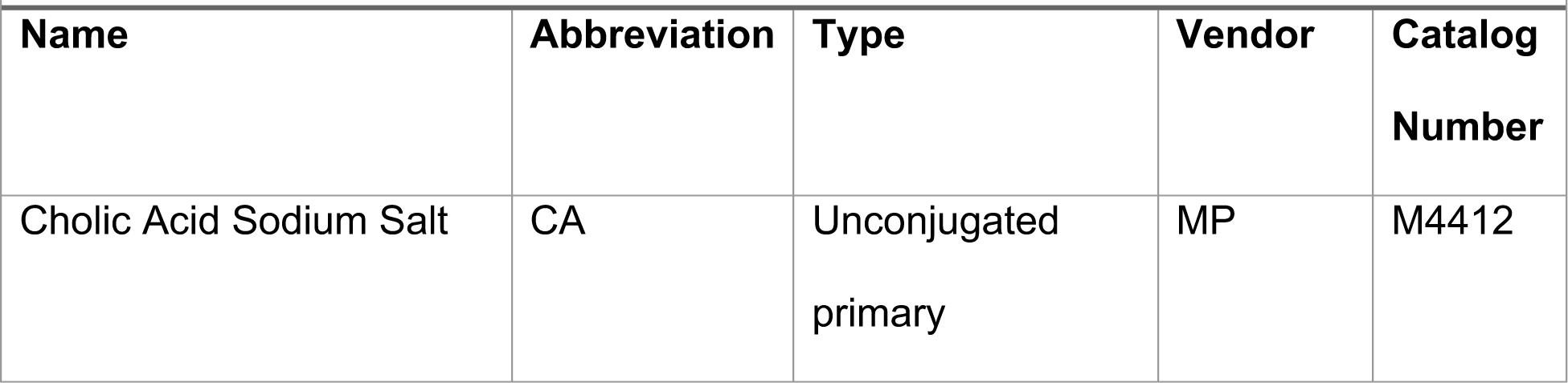
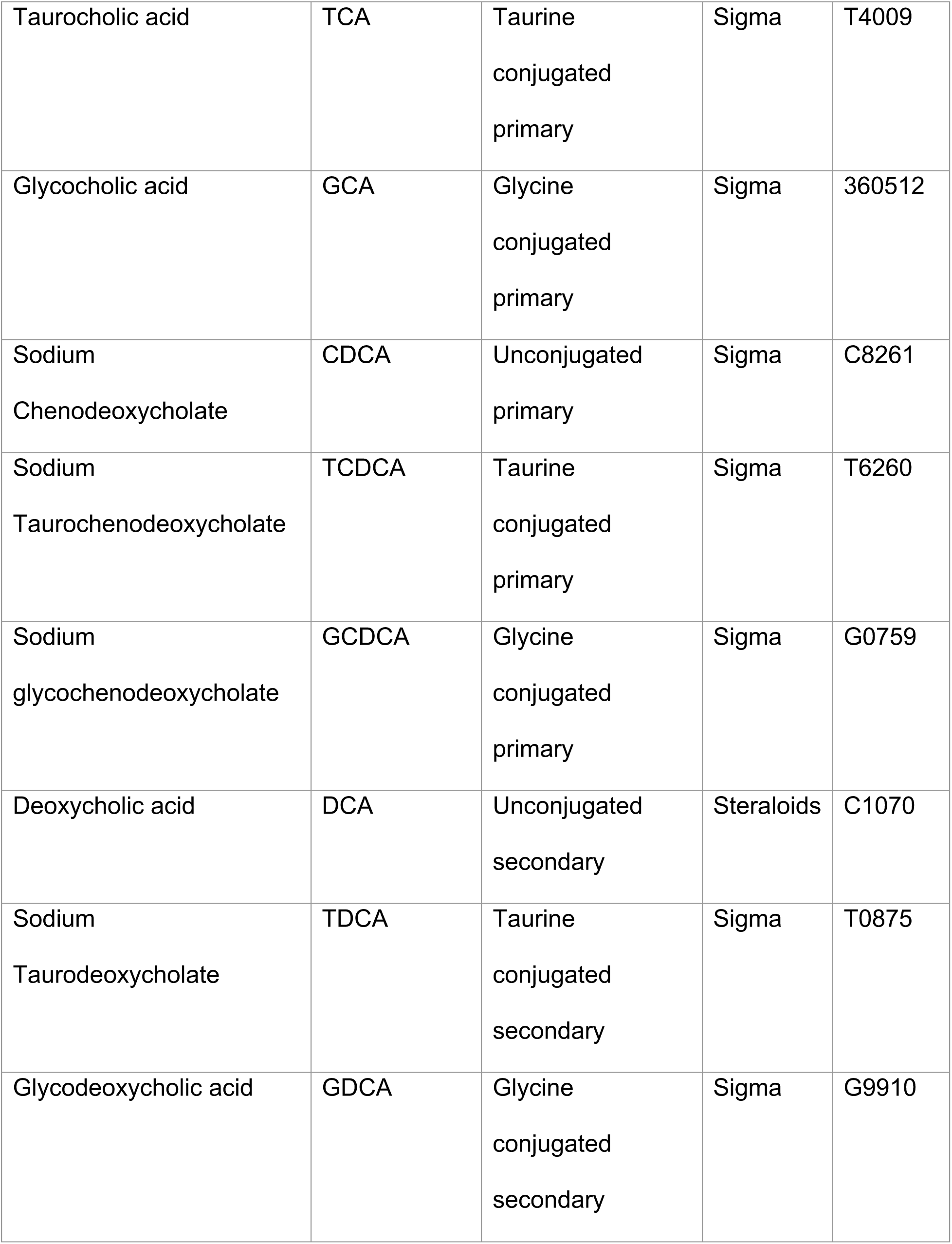
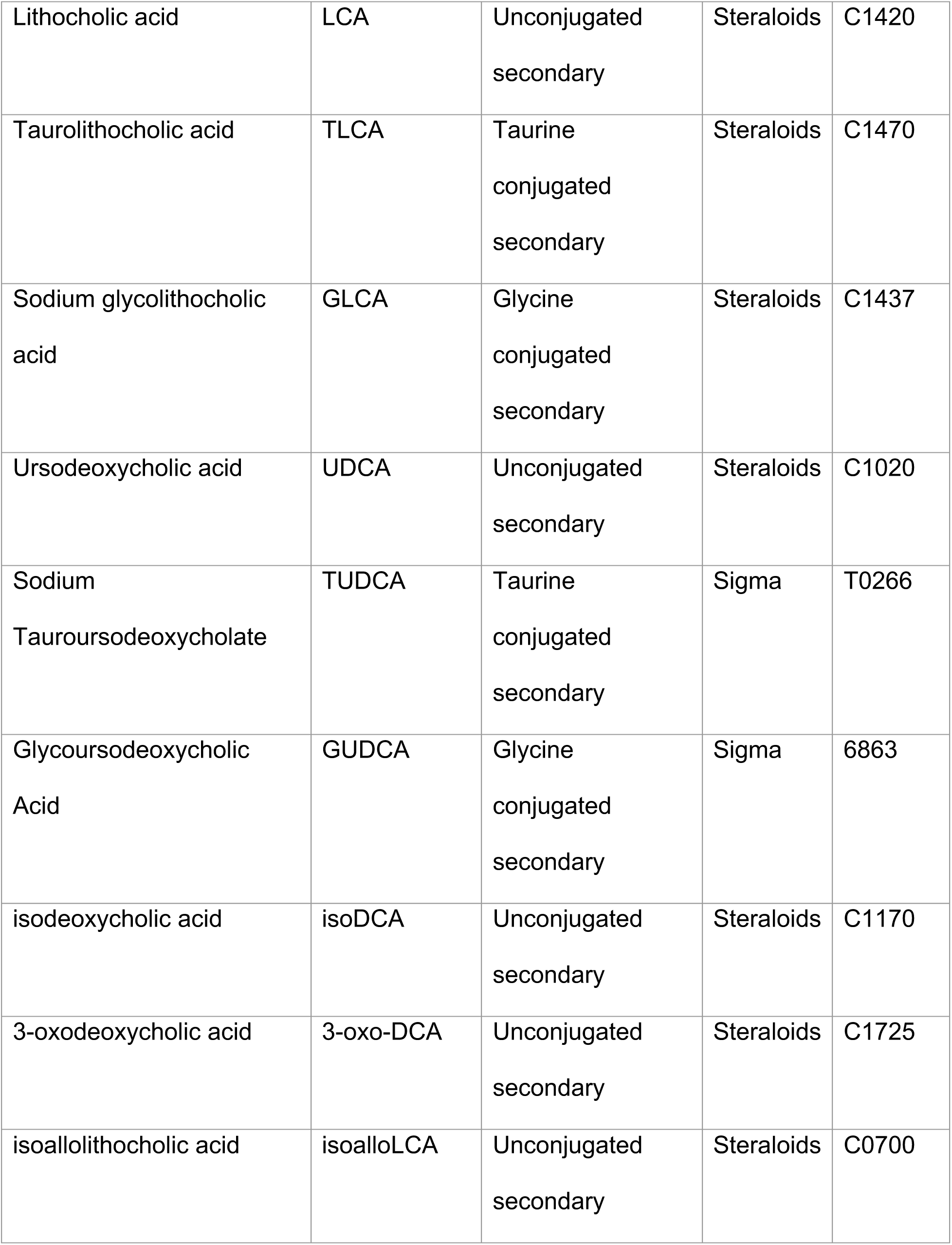
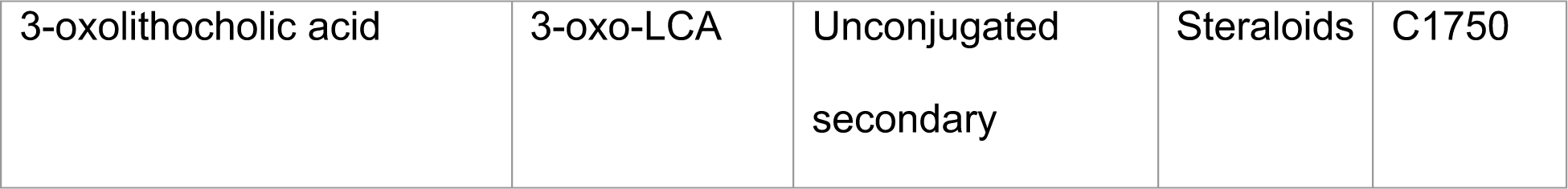
Bile acids used in this study.

**Supplemental Table 2.**
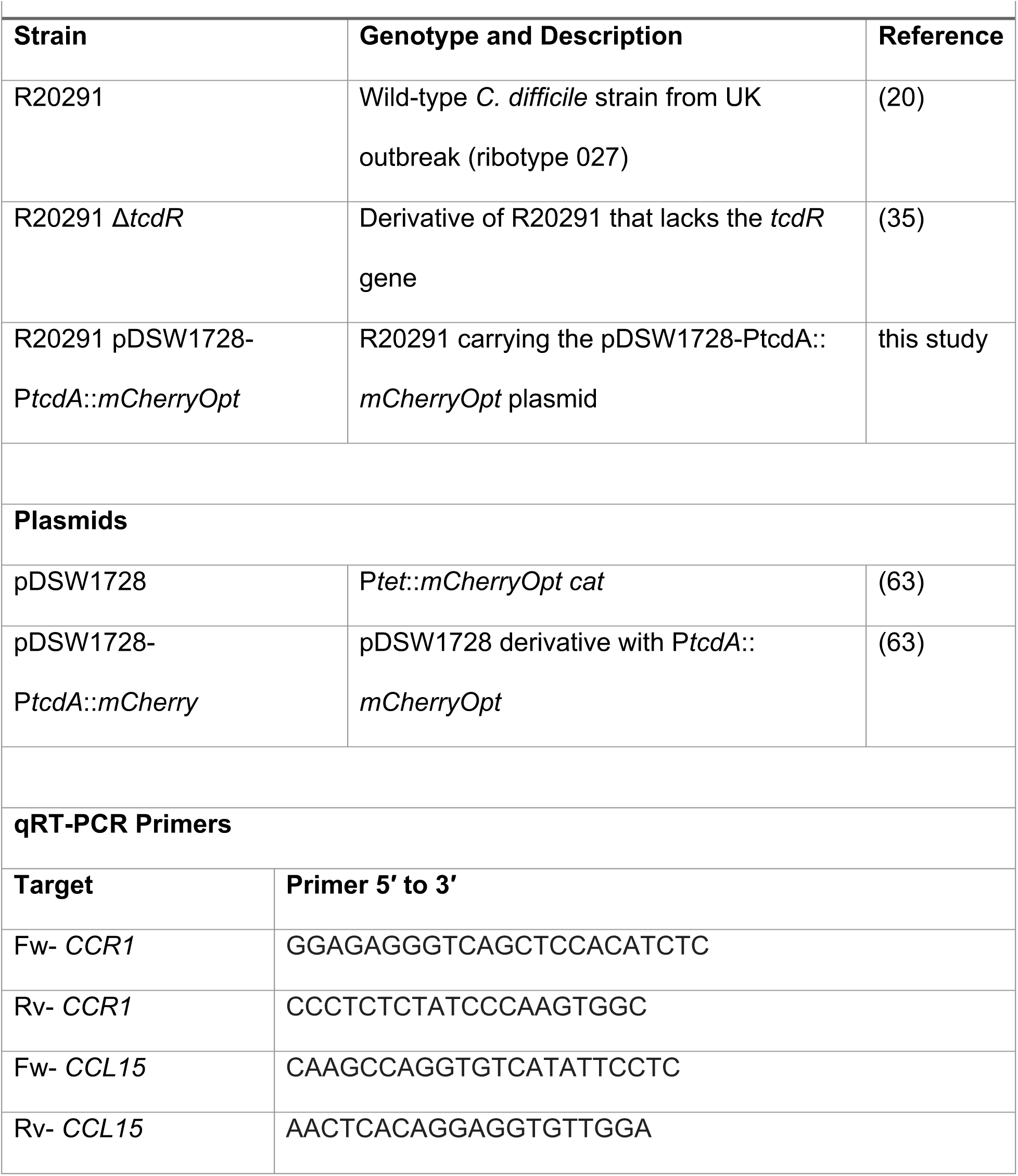
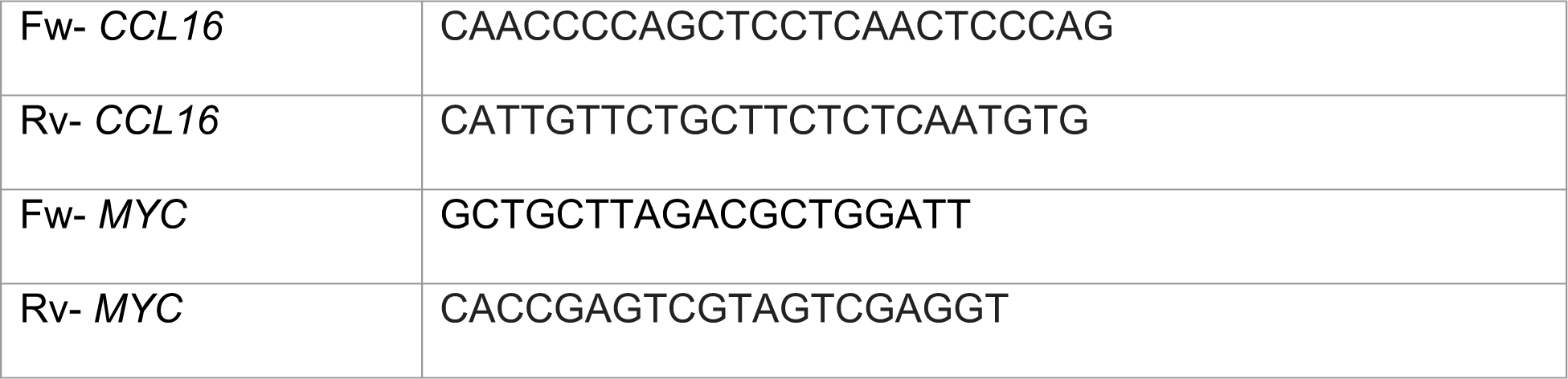
Strains, plasmids and primers used in this study.

**Supplemental Table 3.**
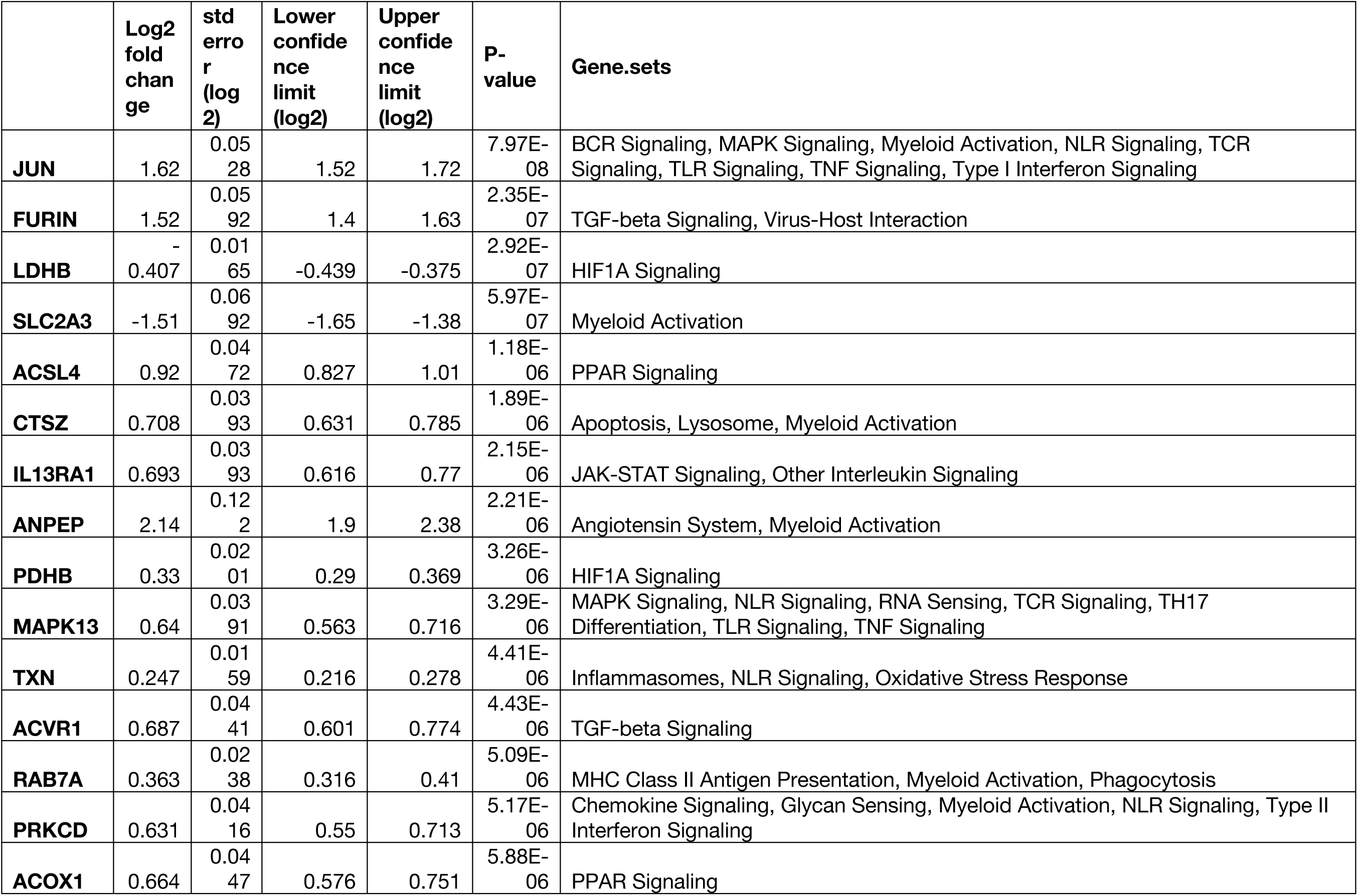
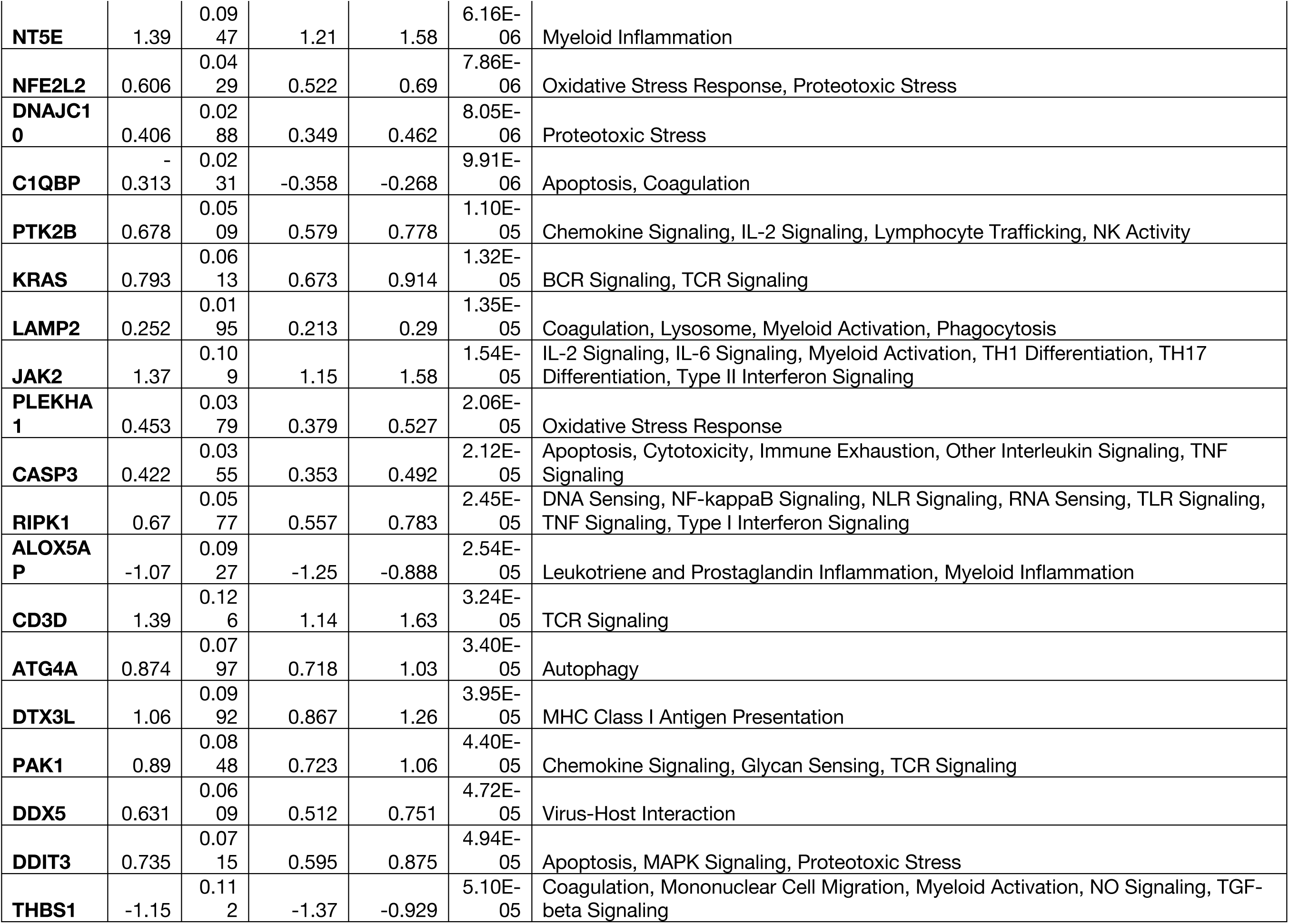
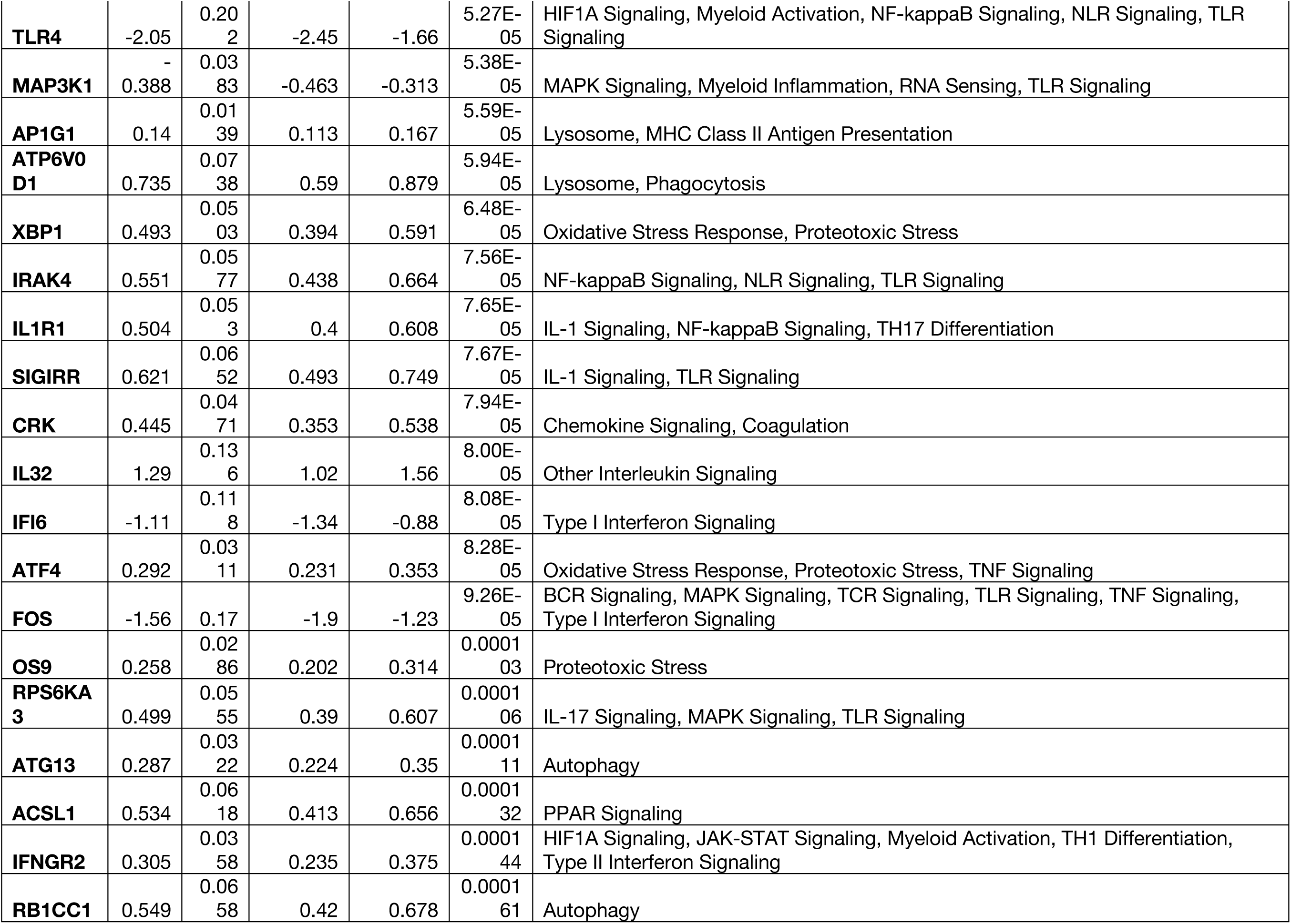
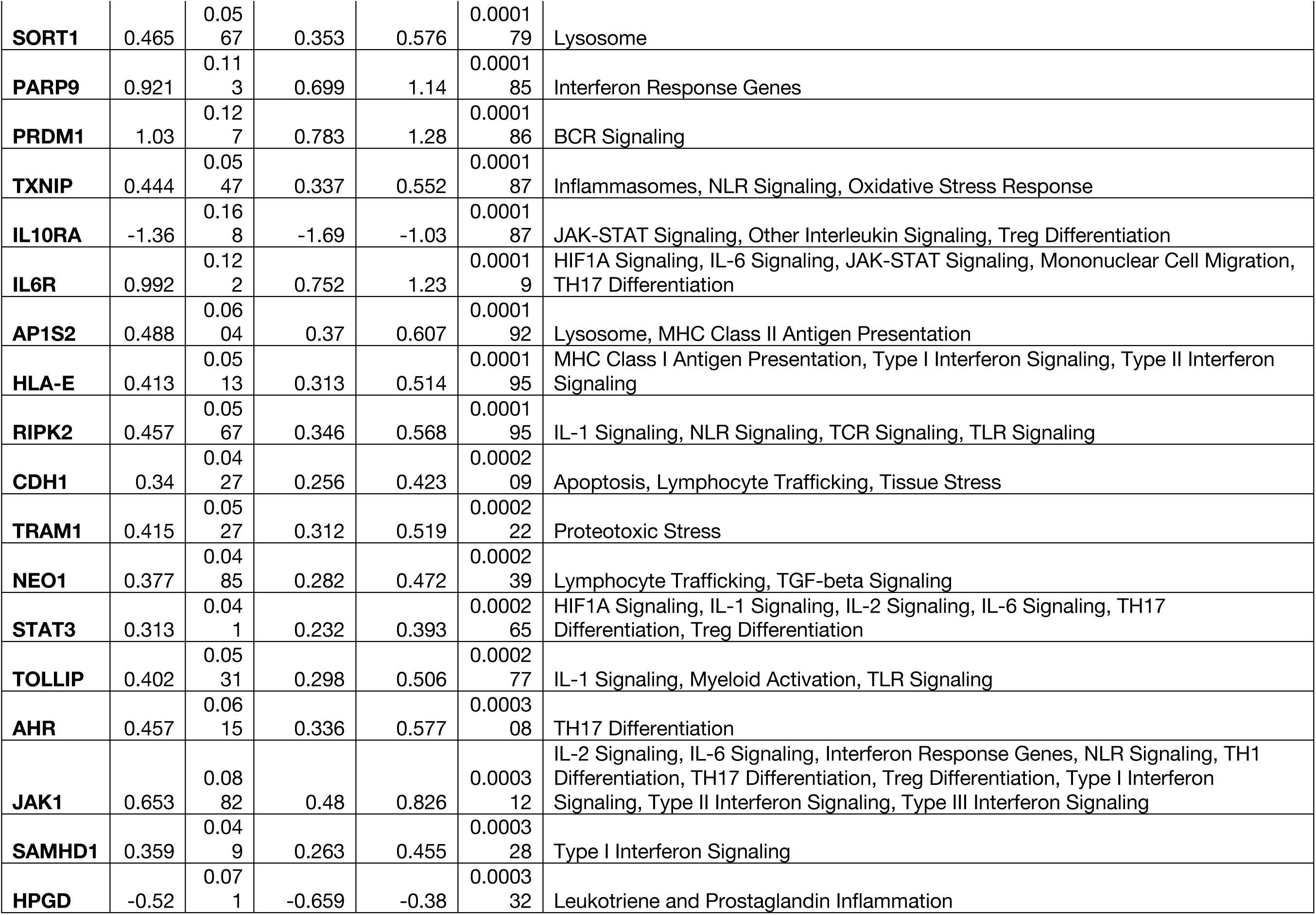
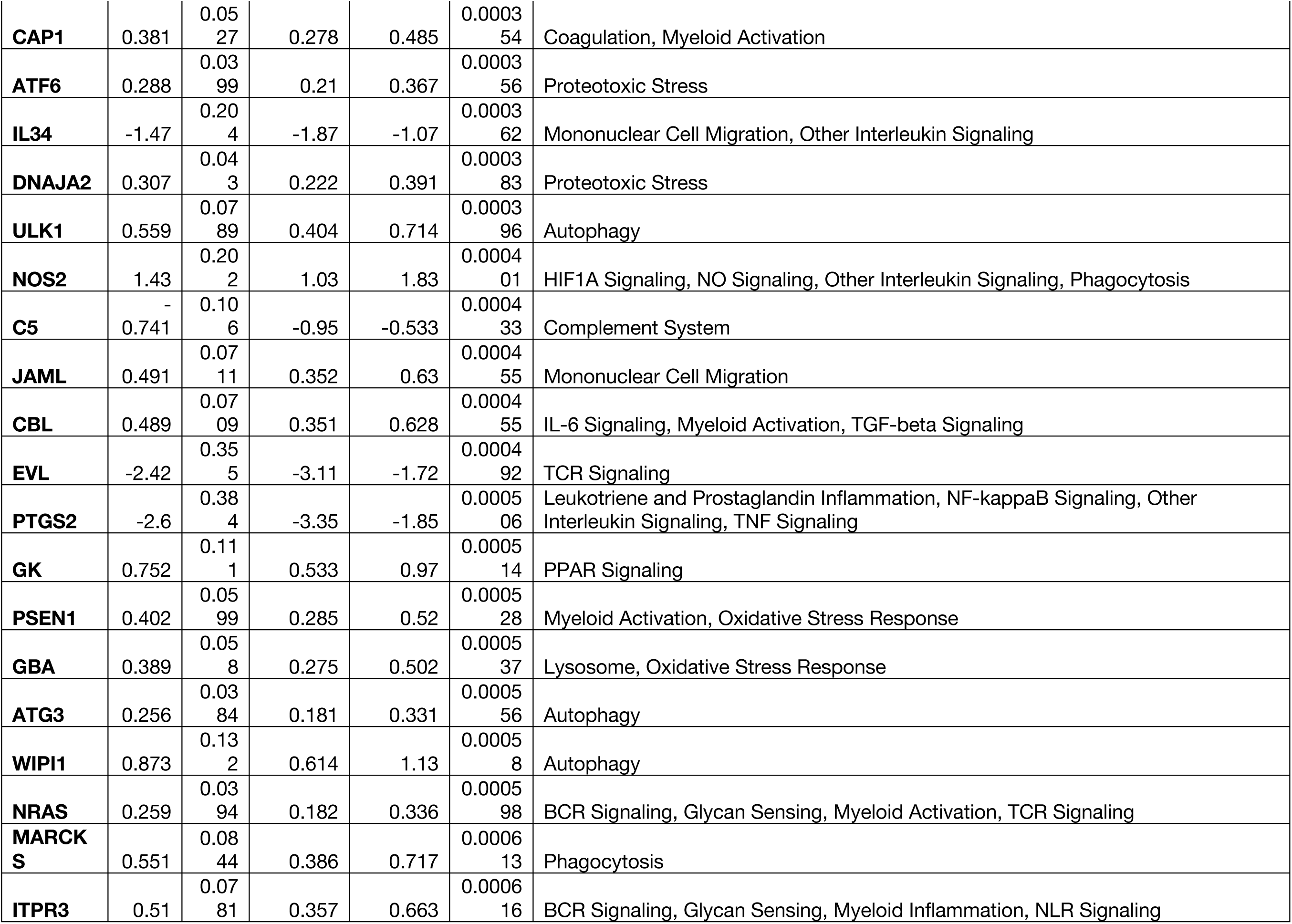
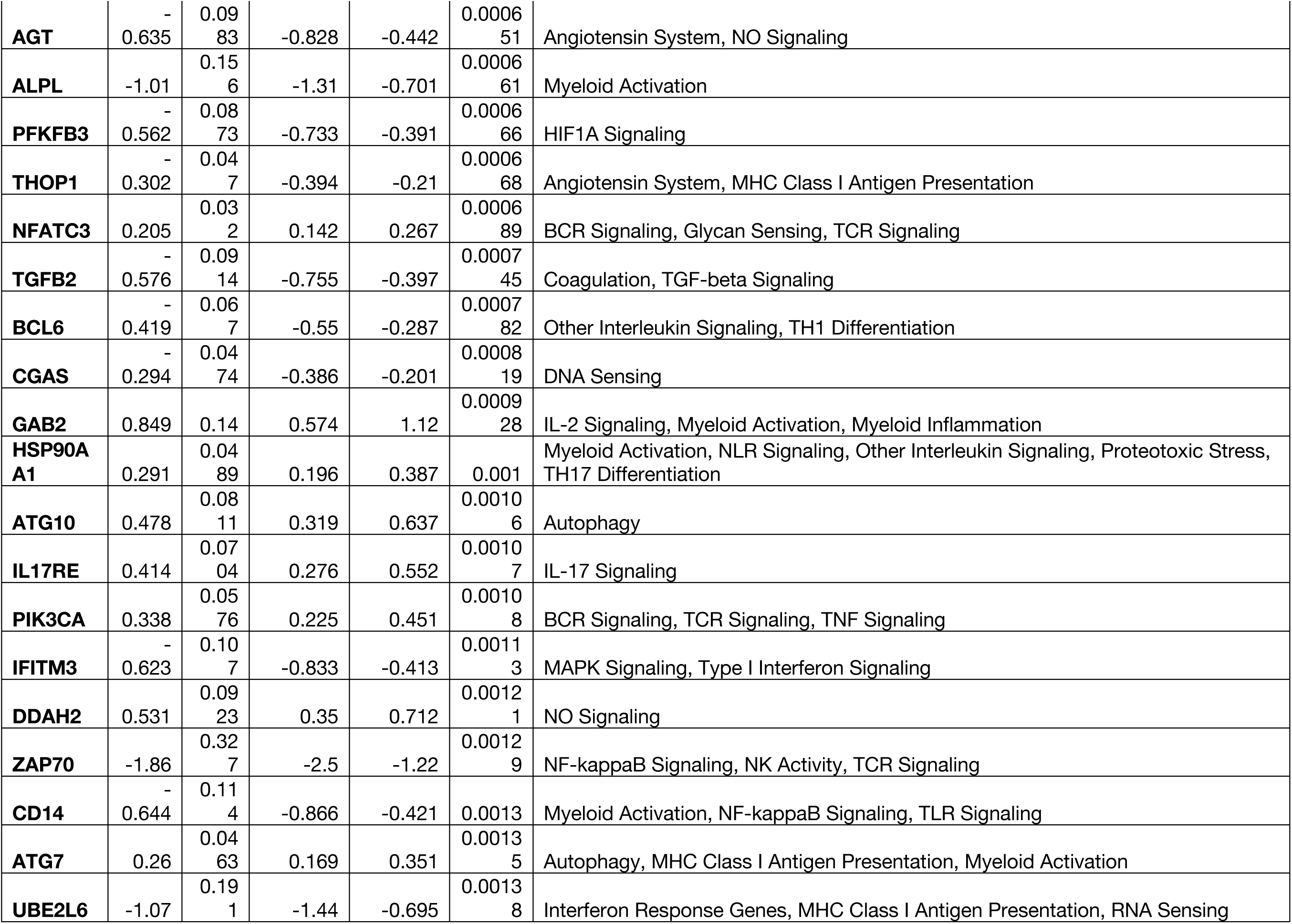
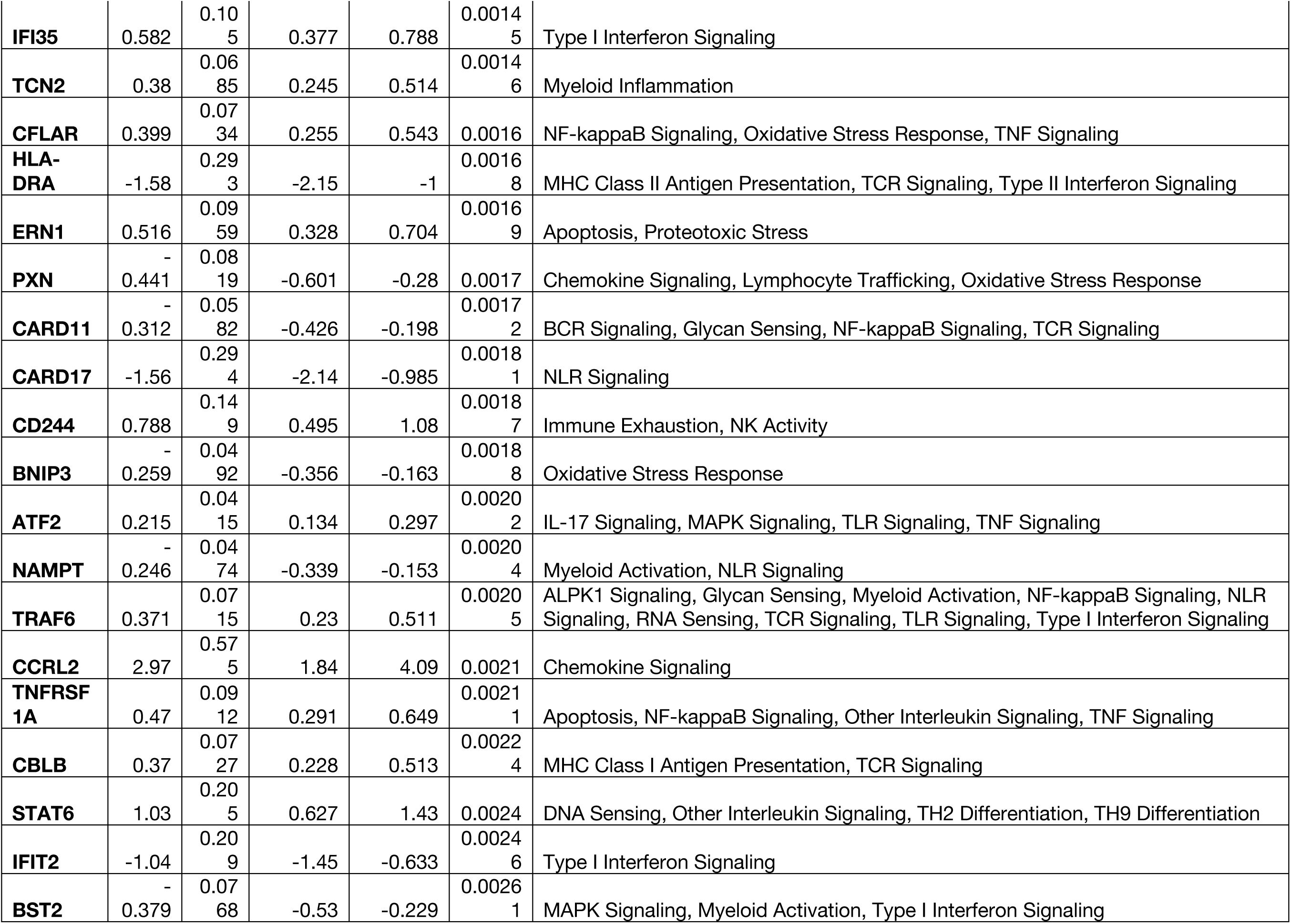
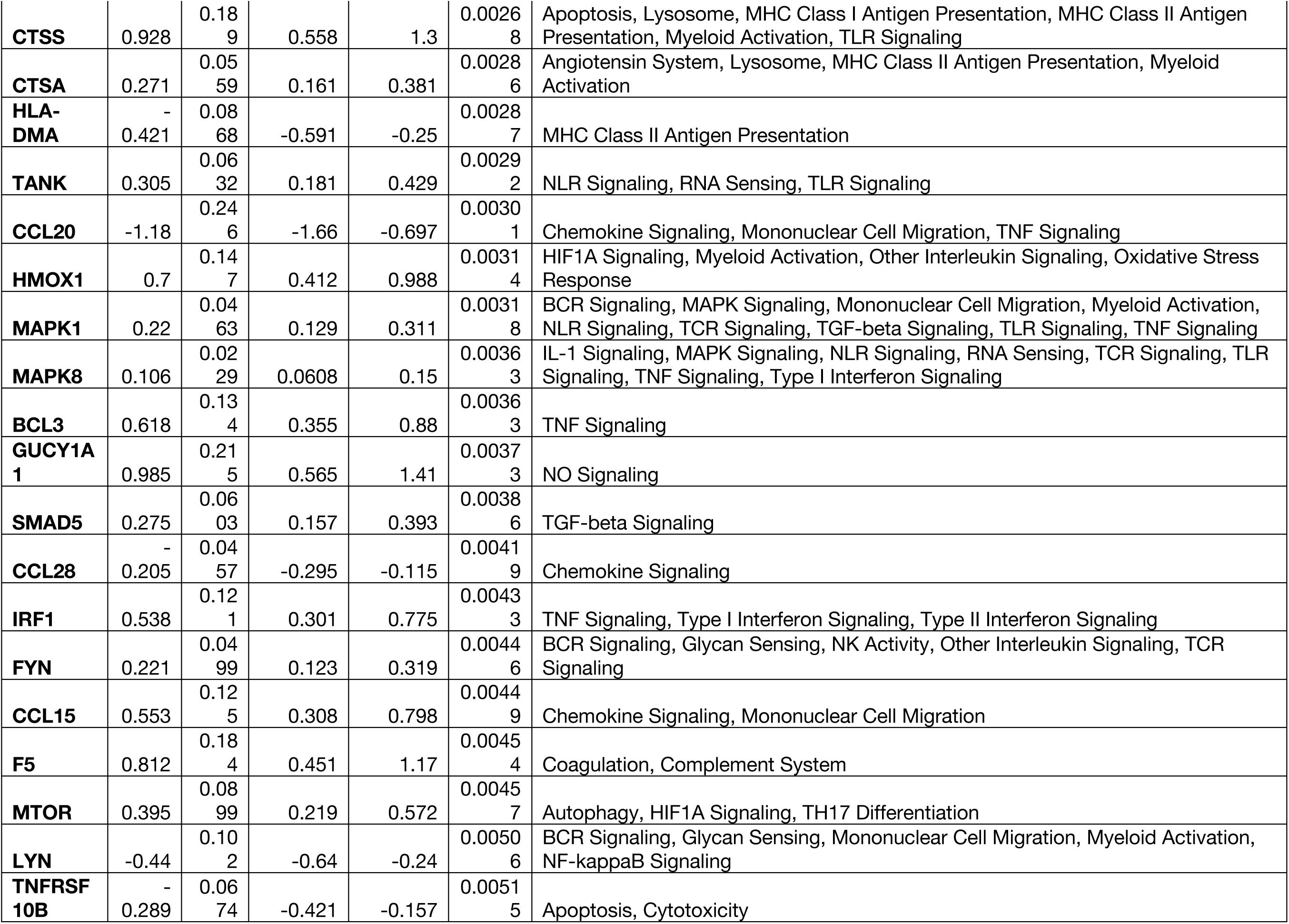
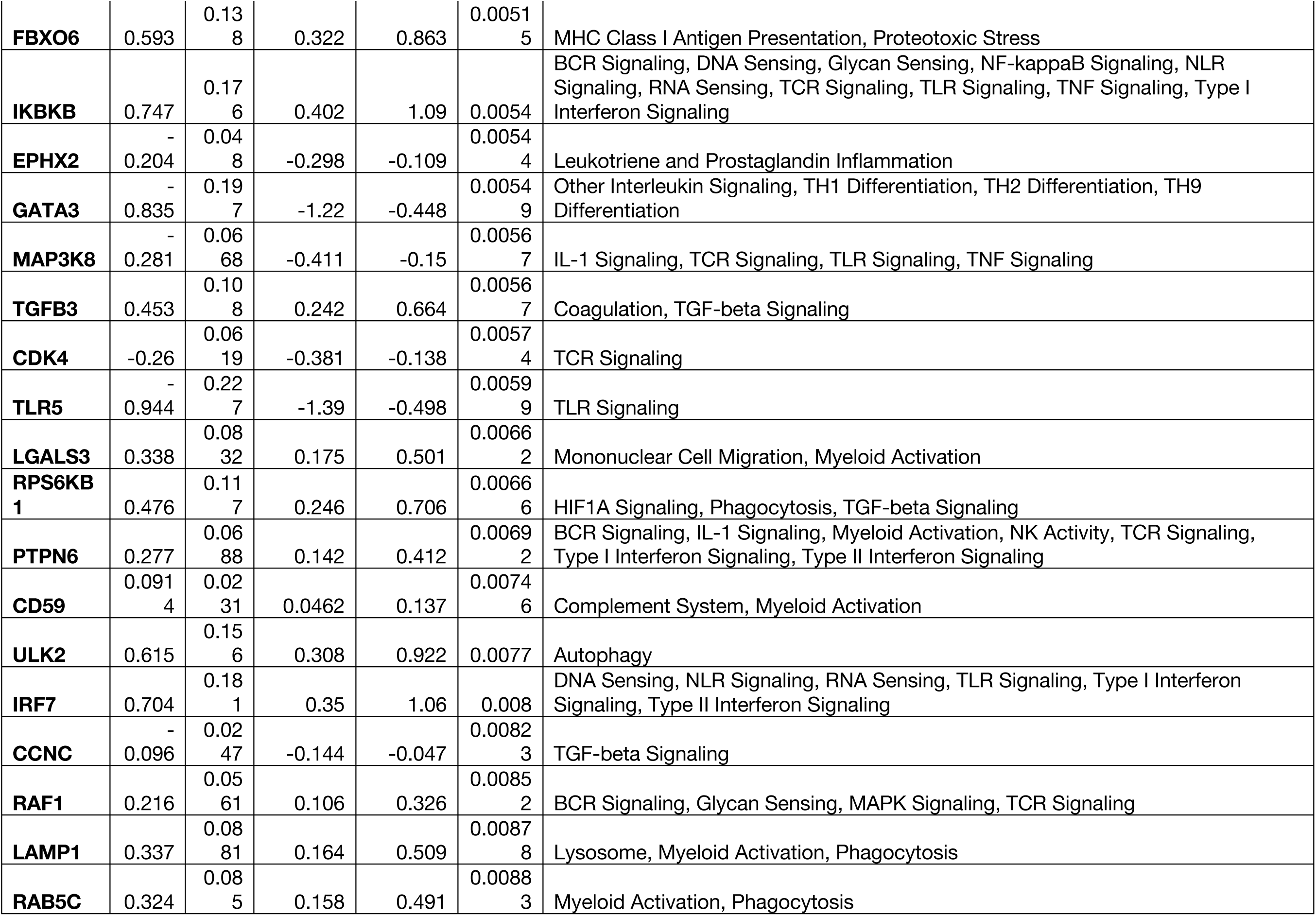
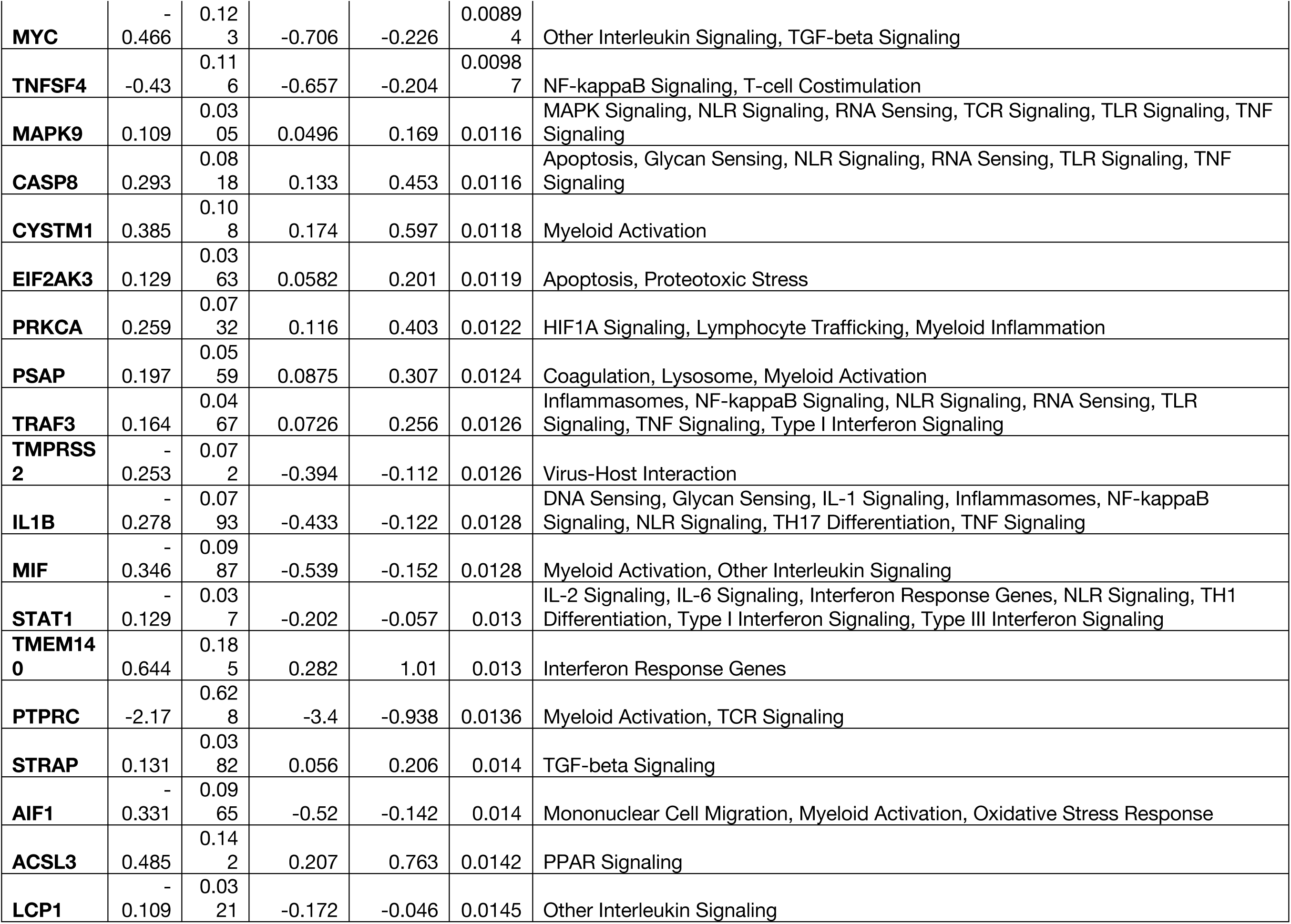
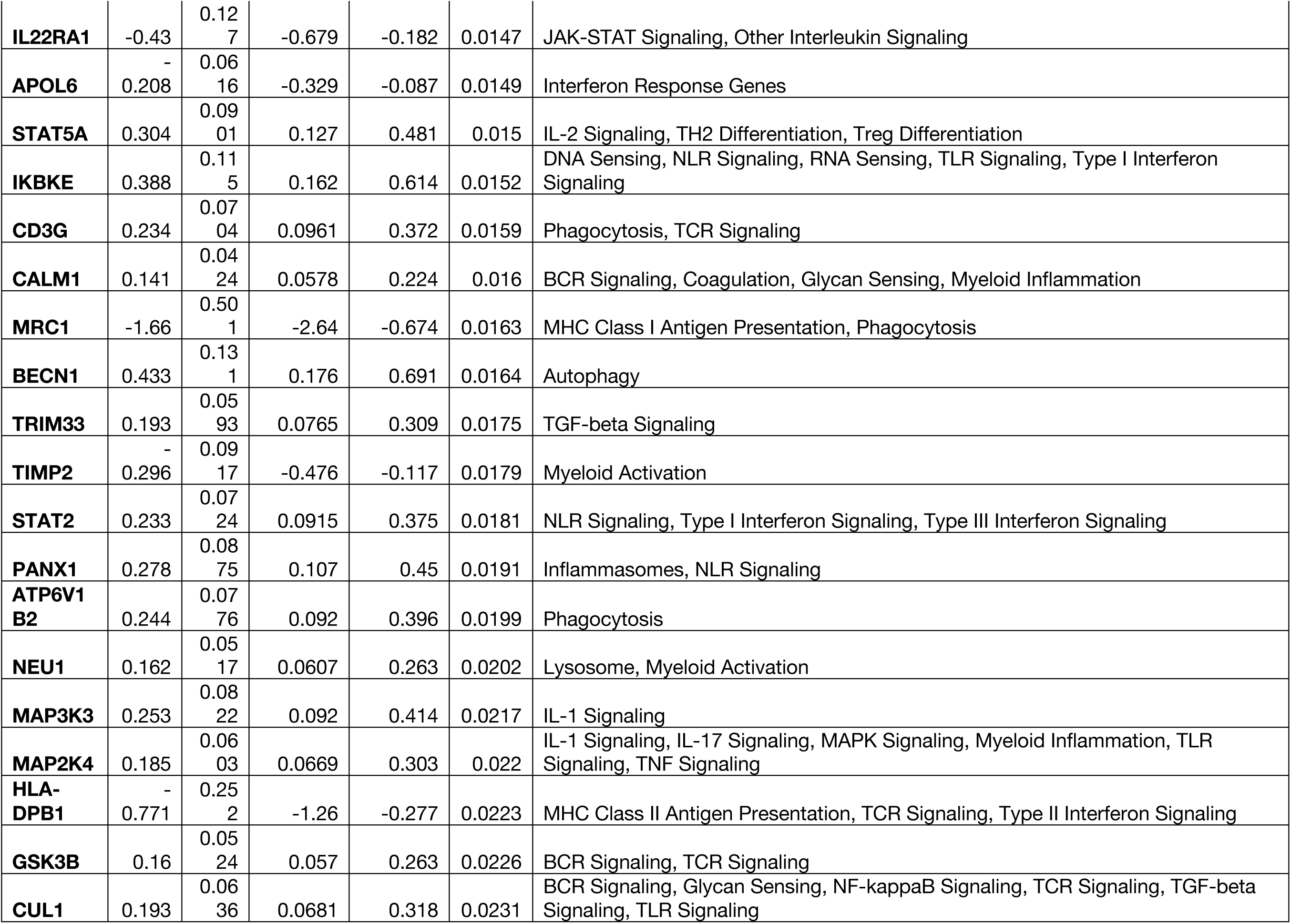
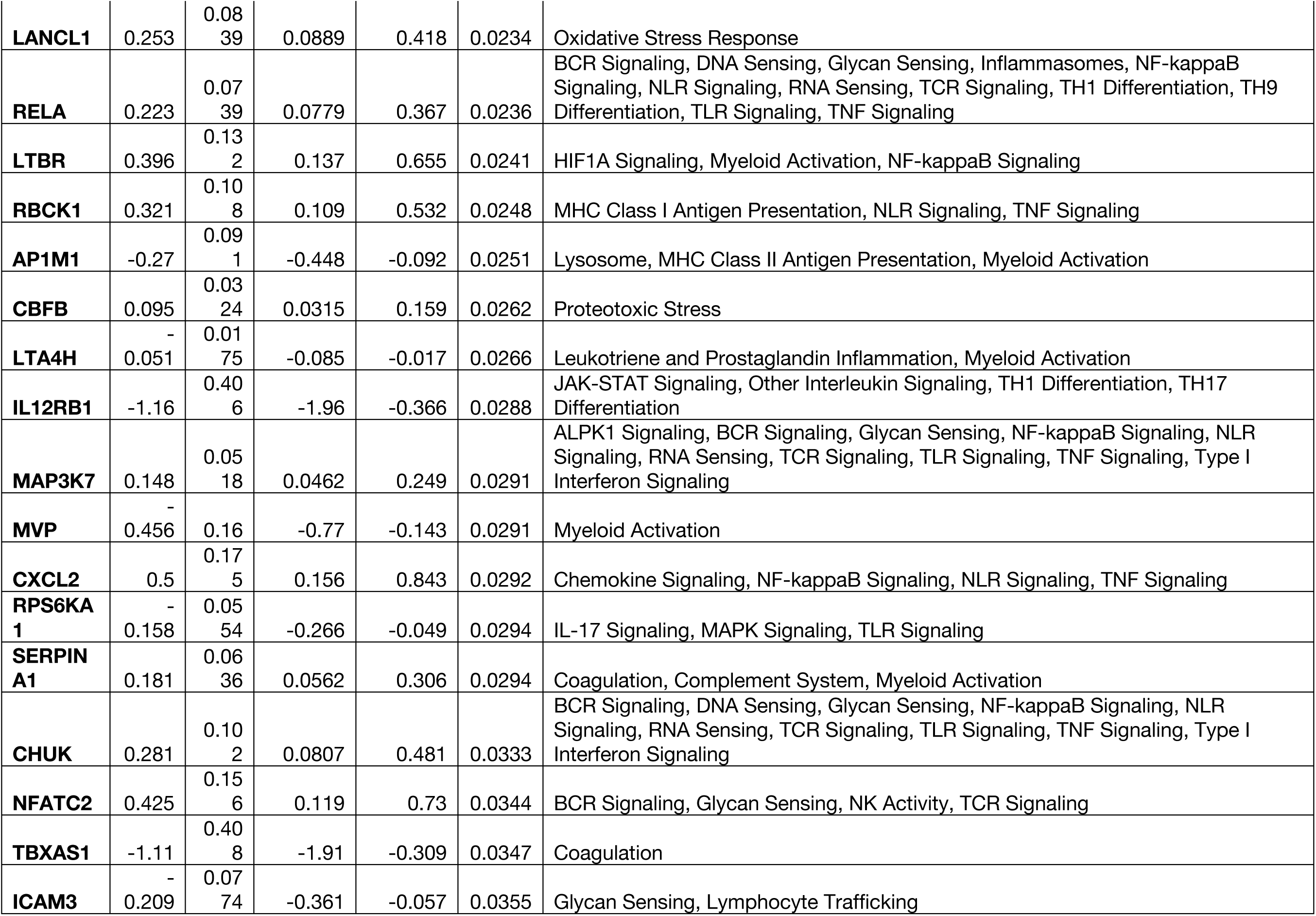
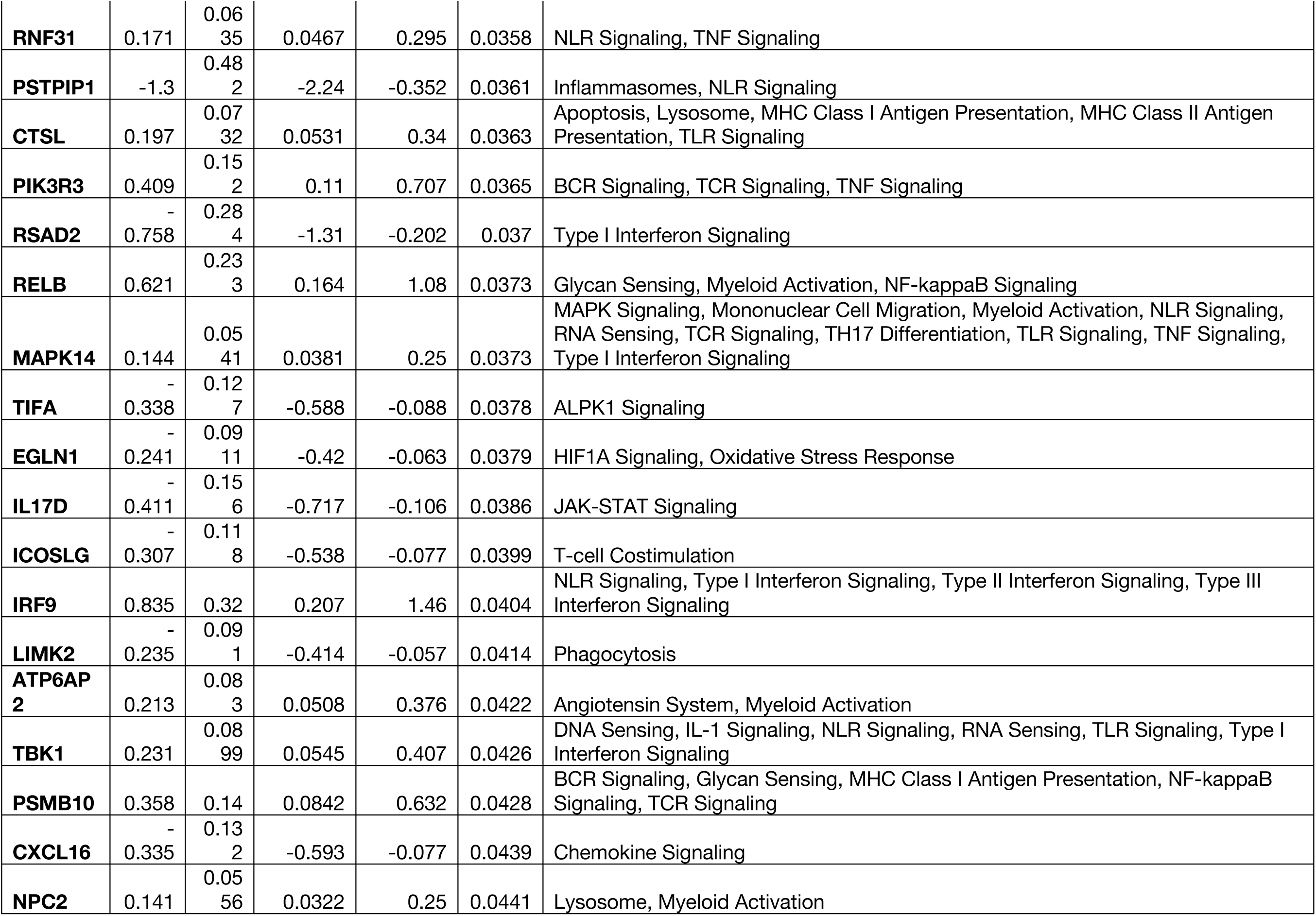
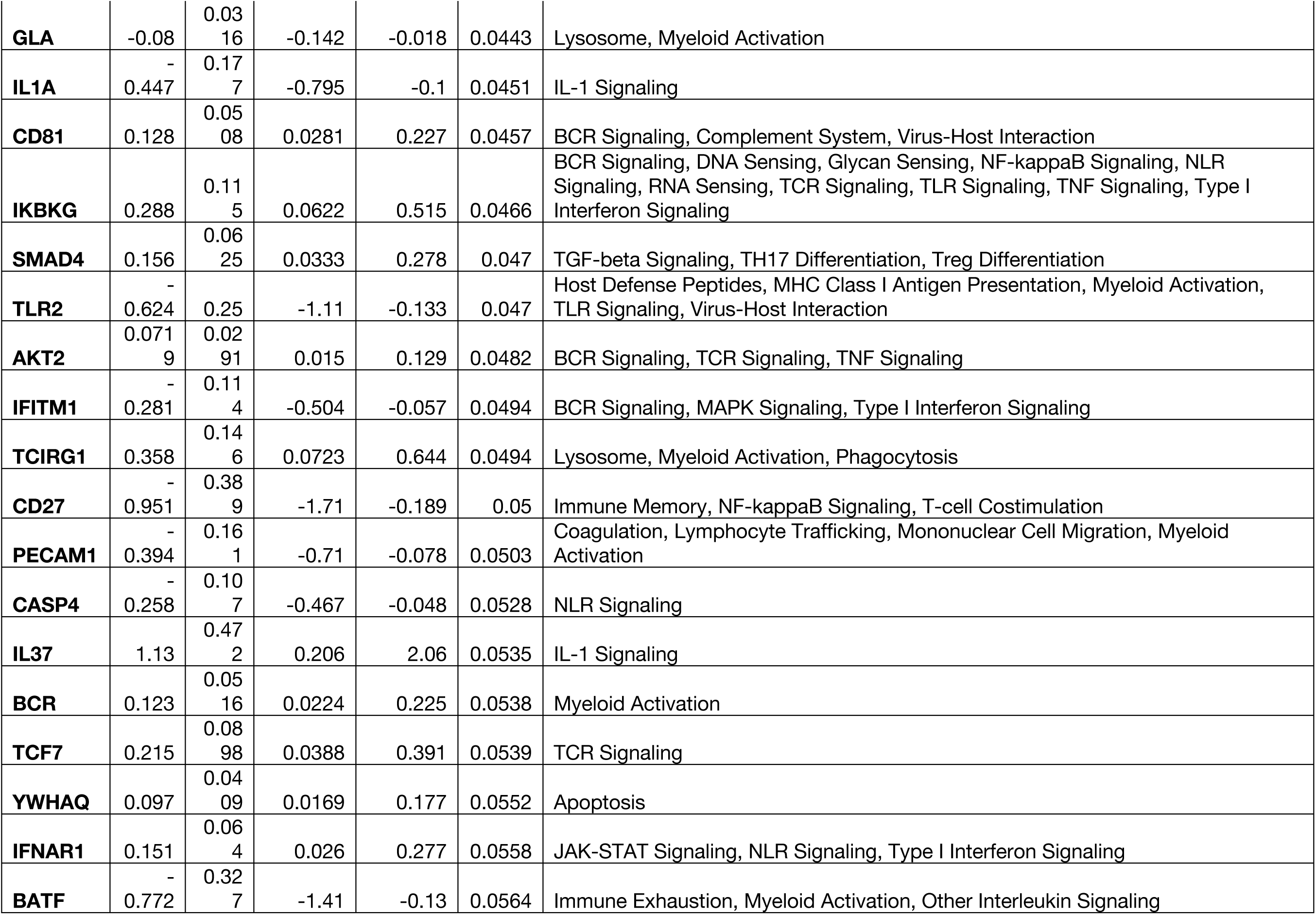
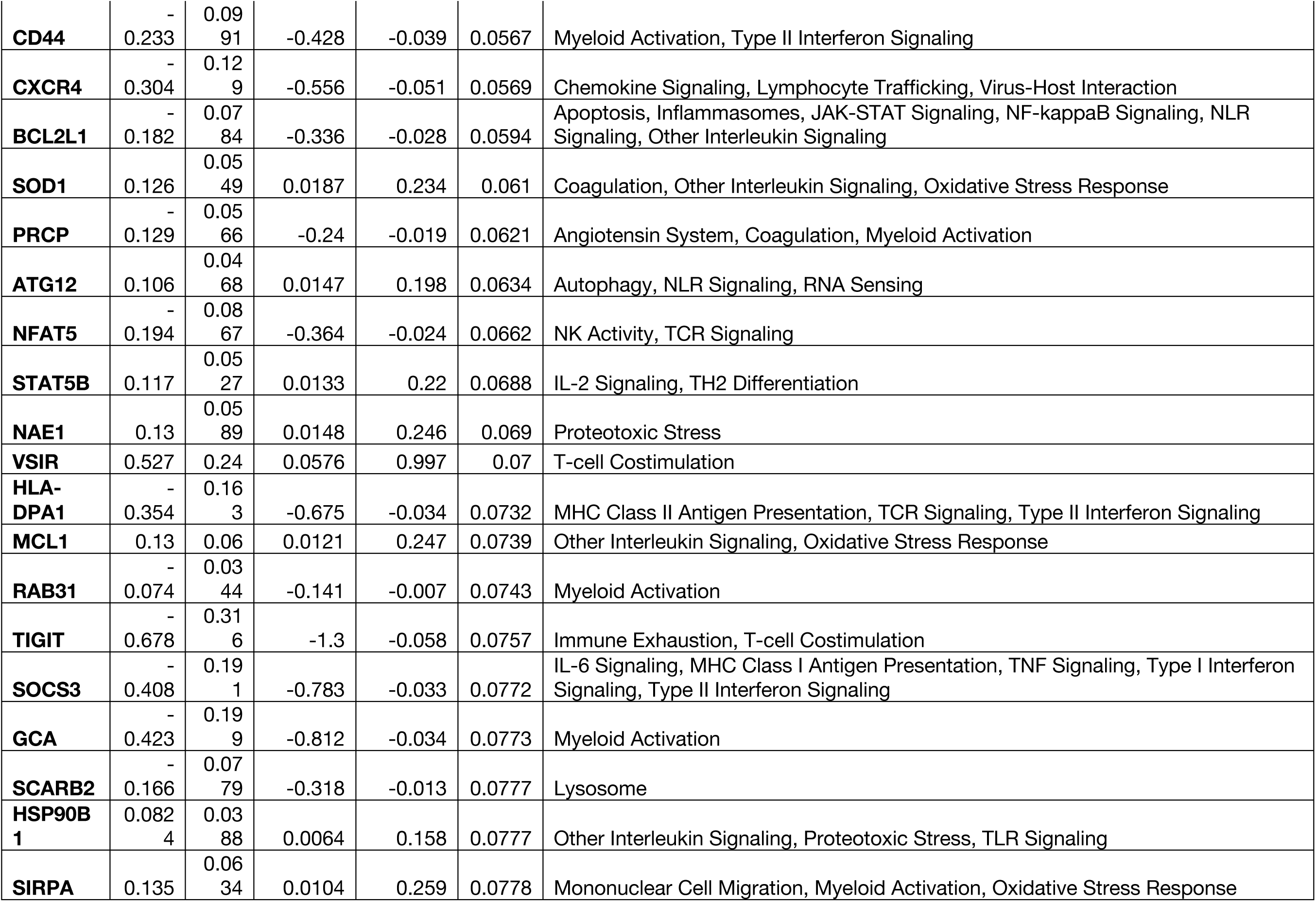
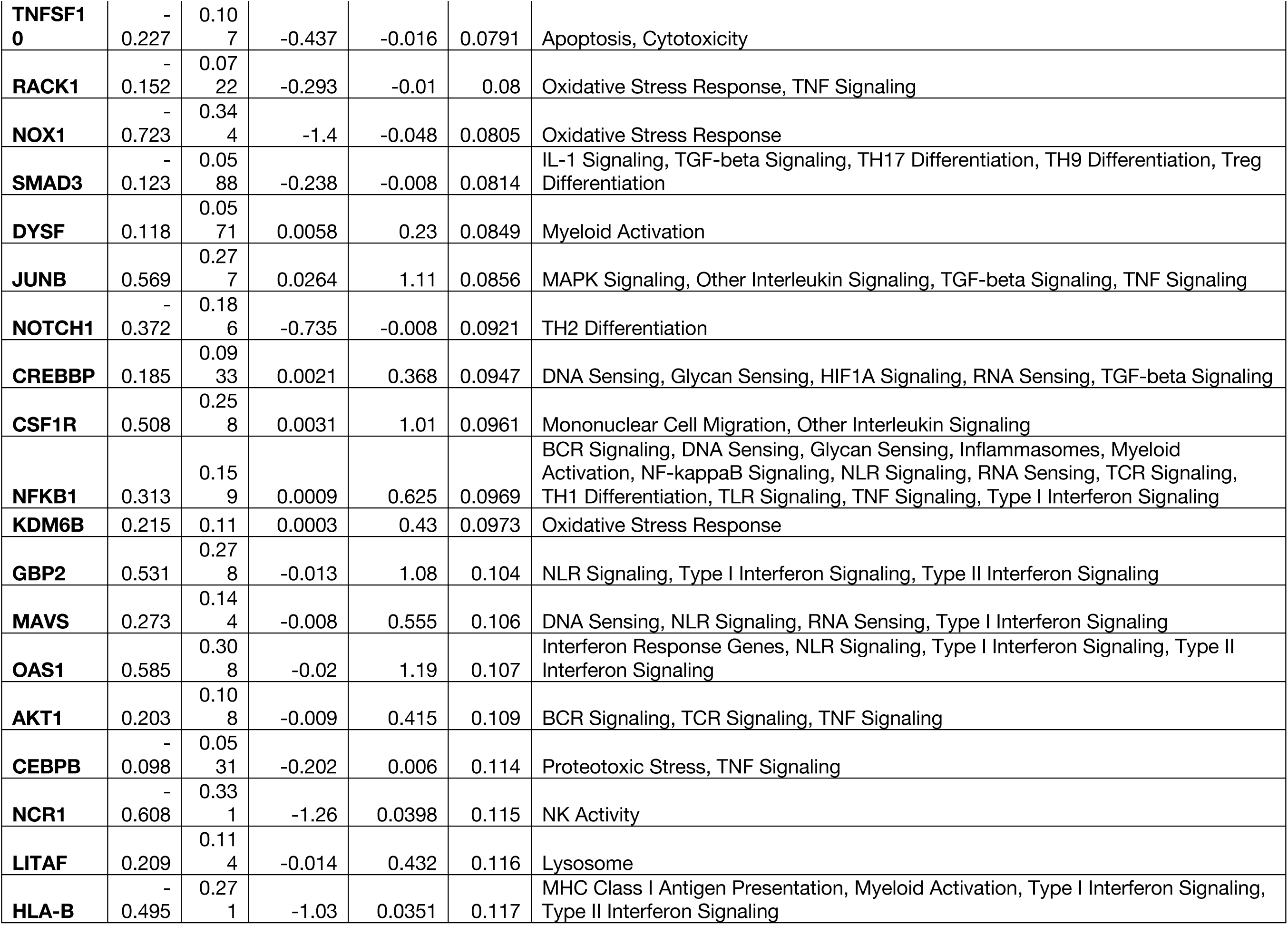
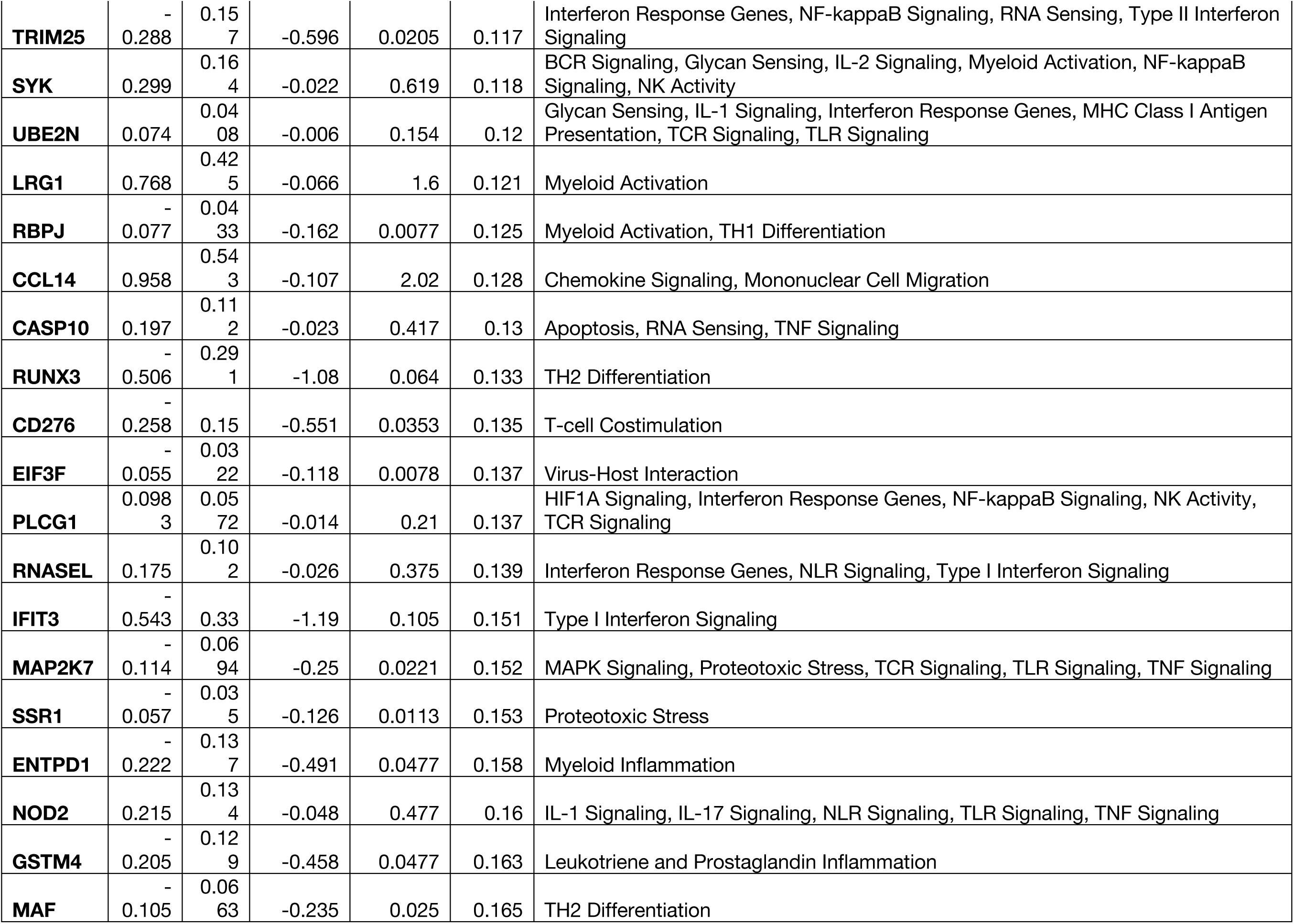
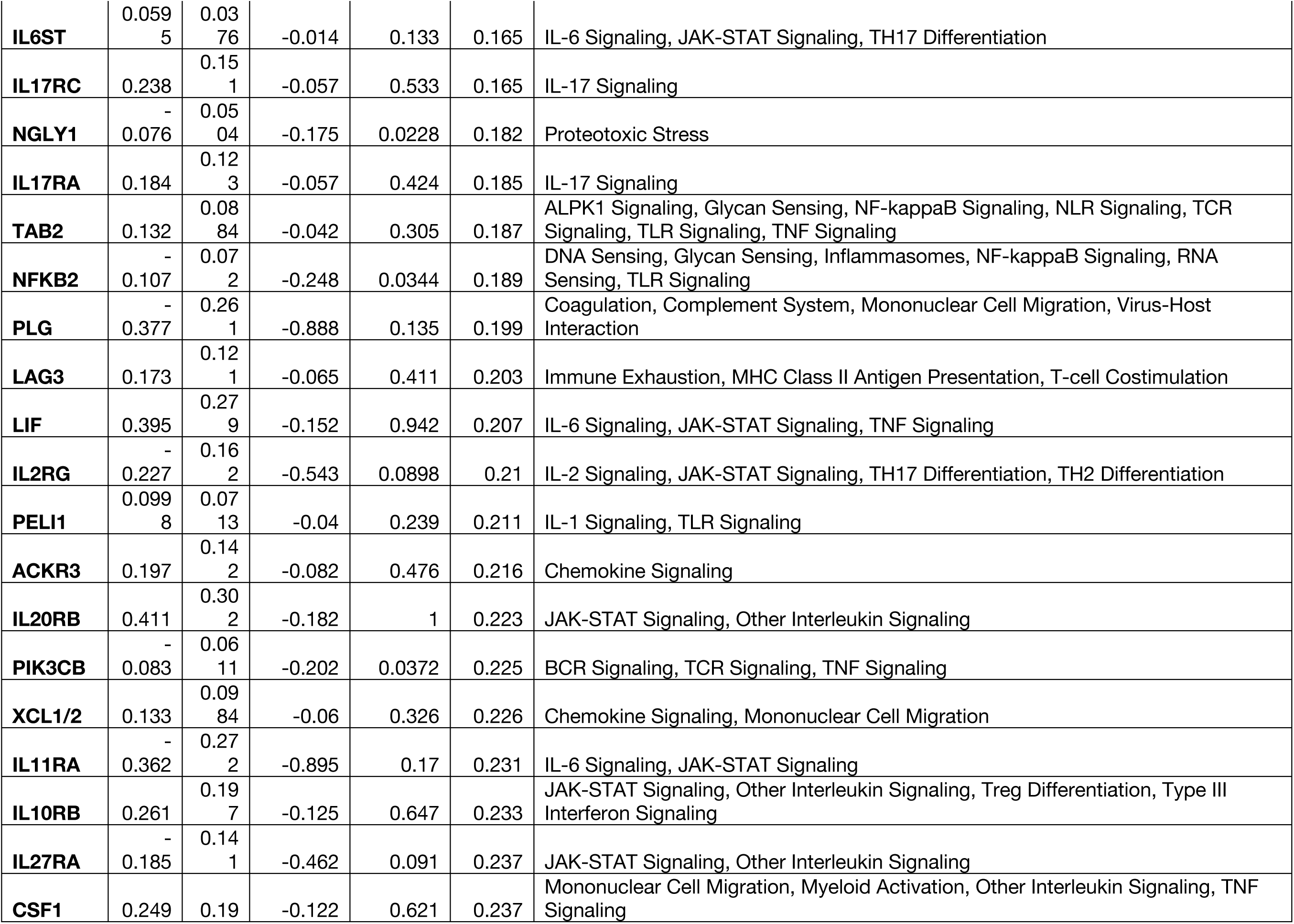
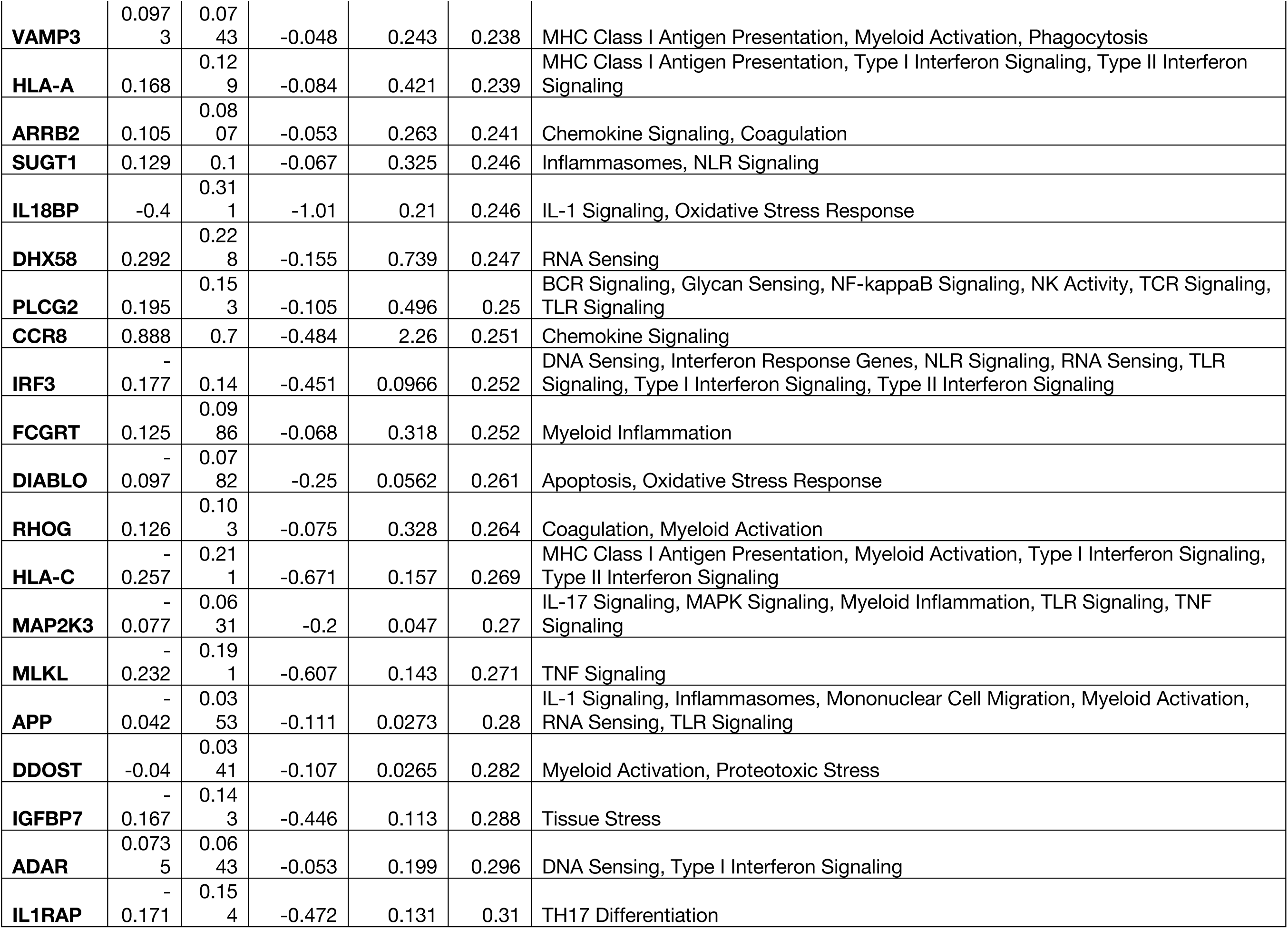
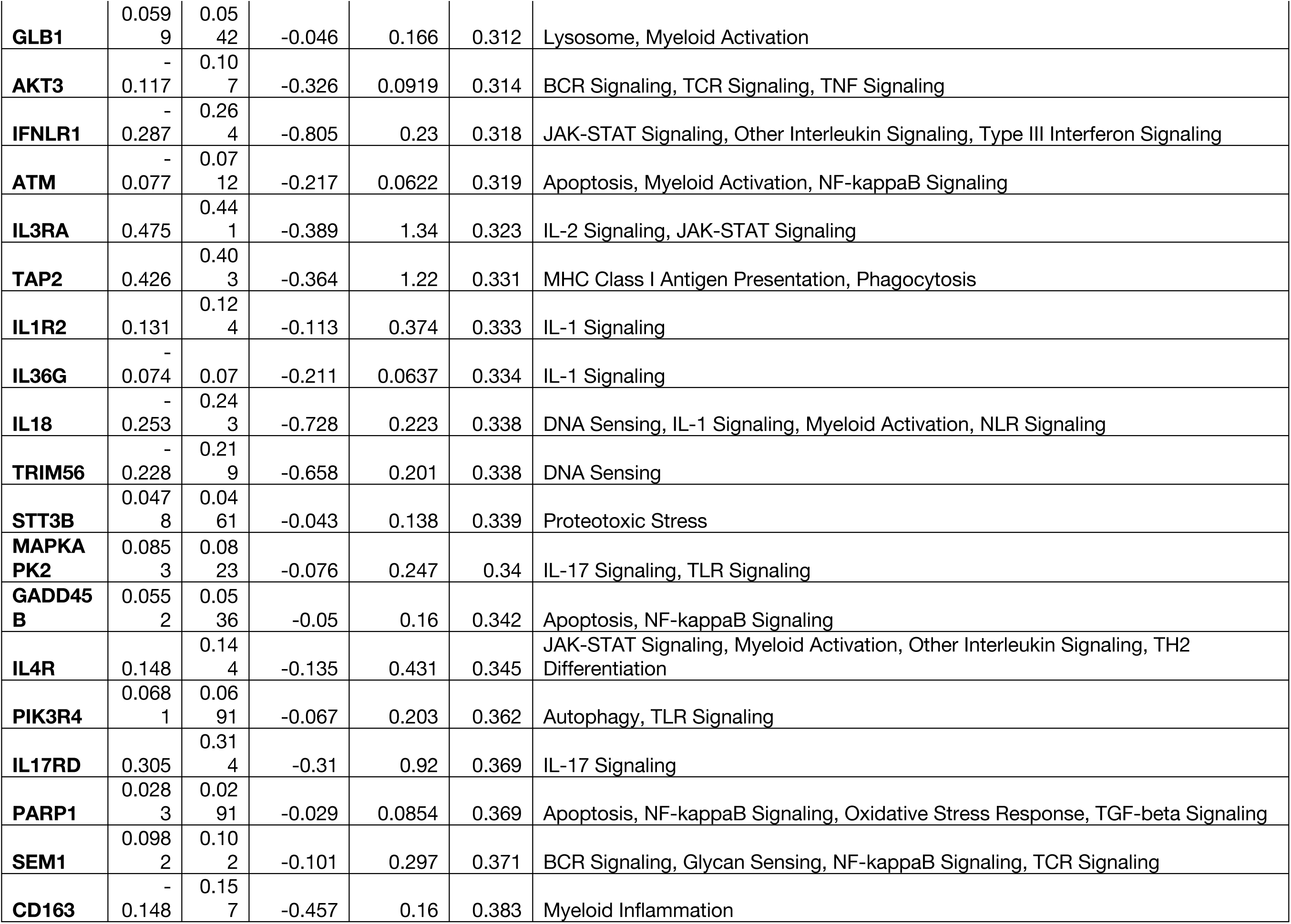
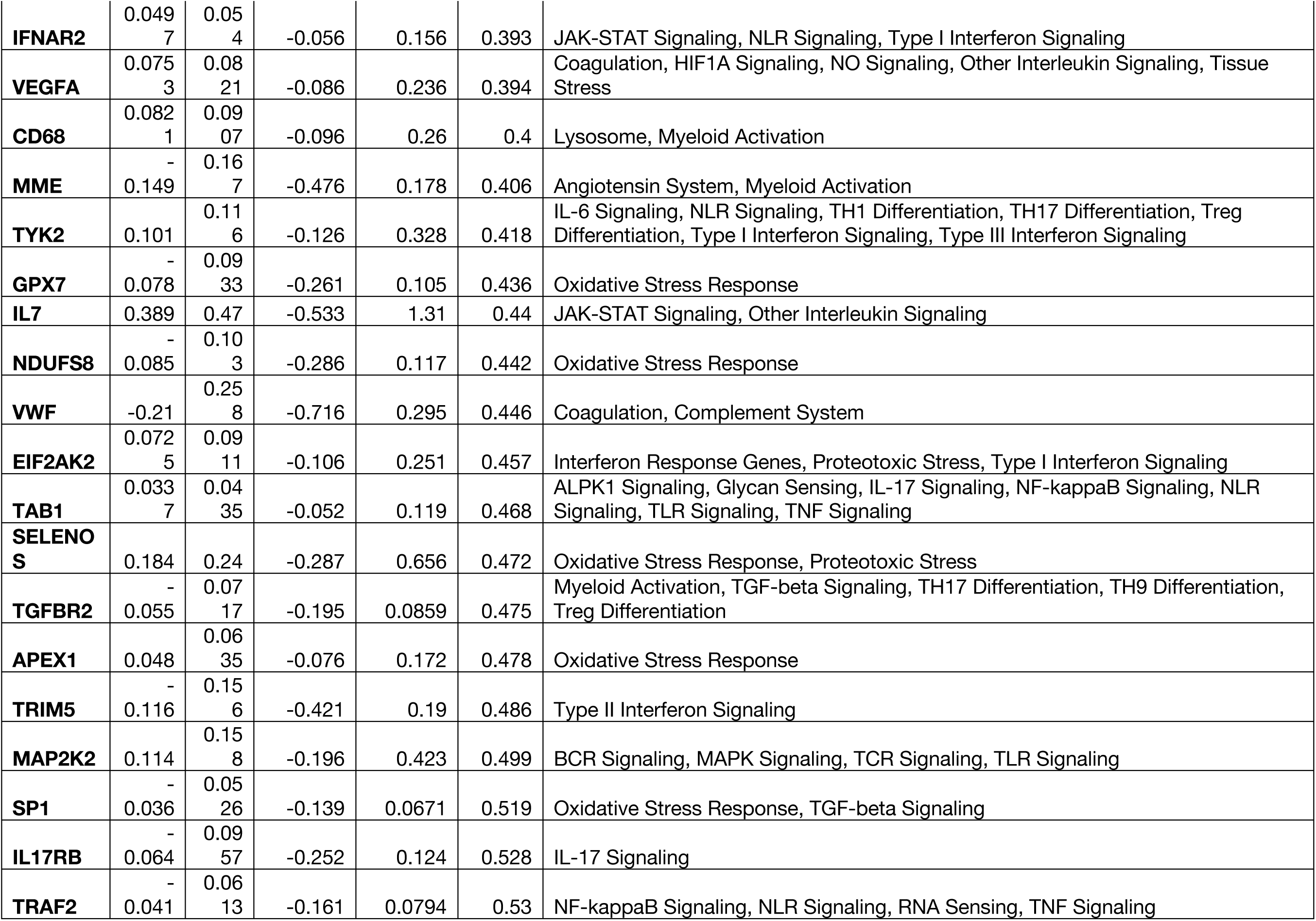
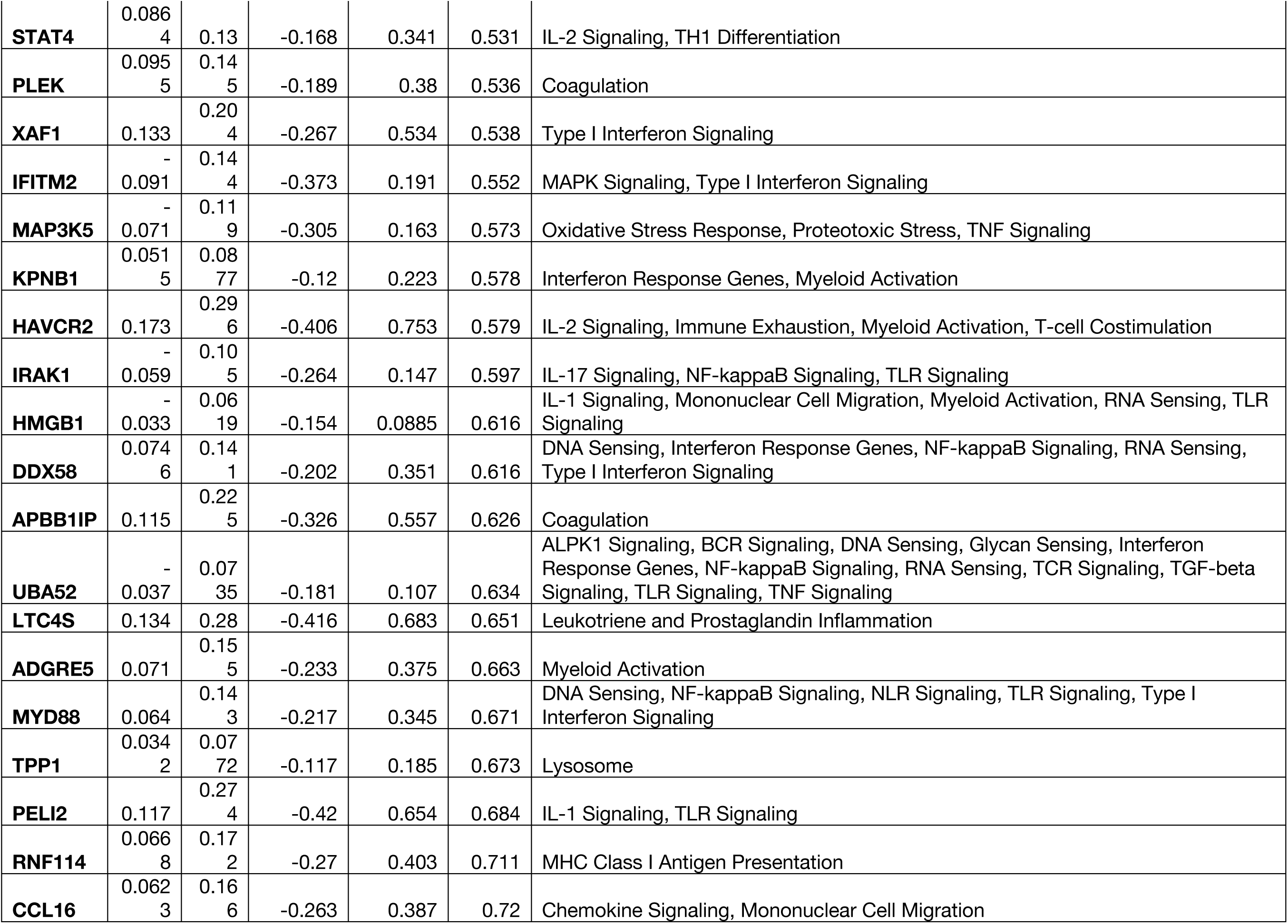
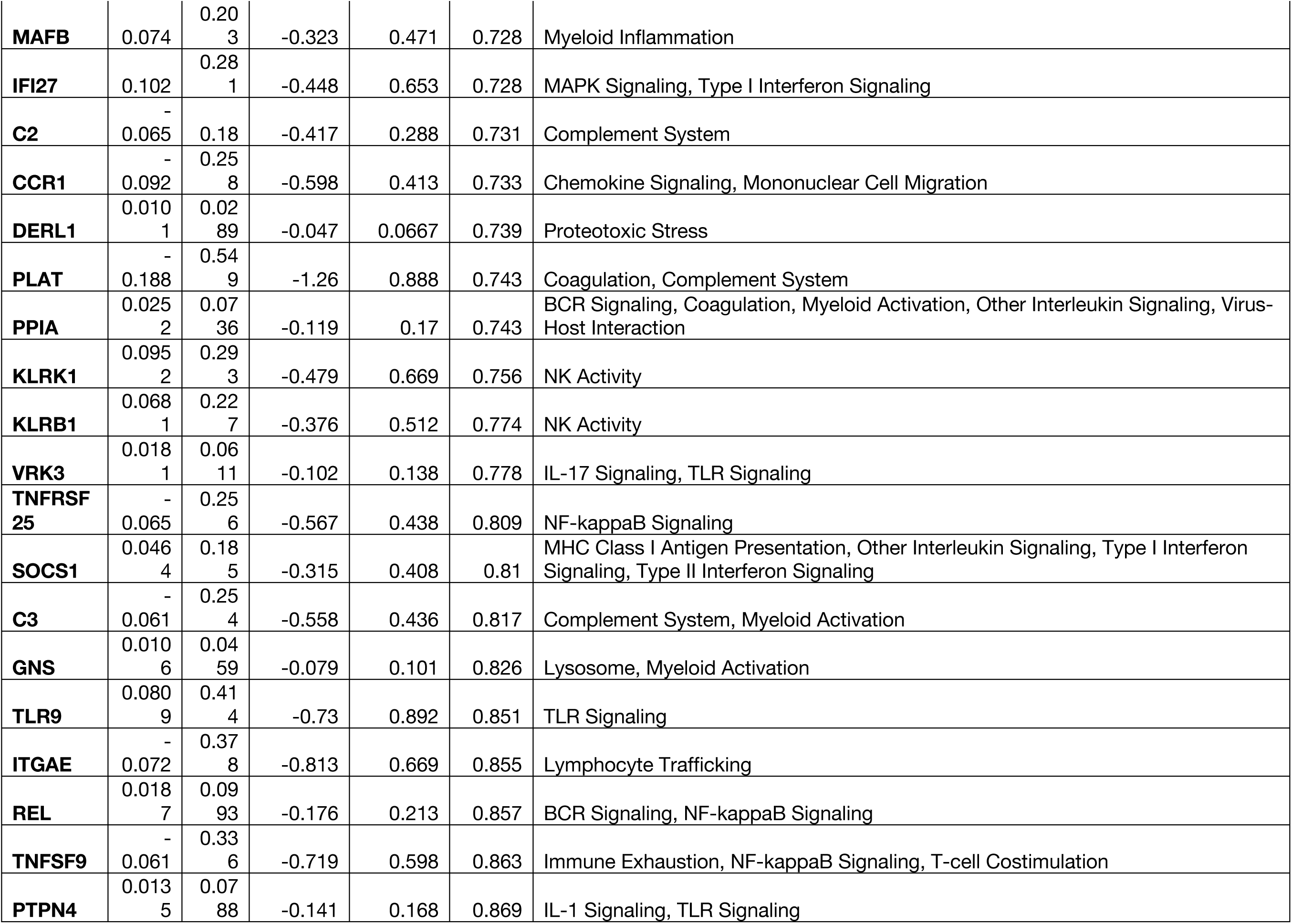
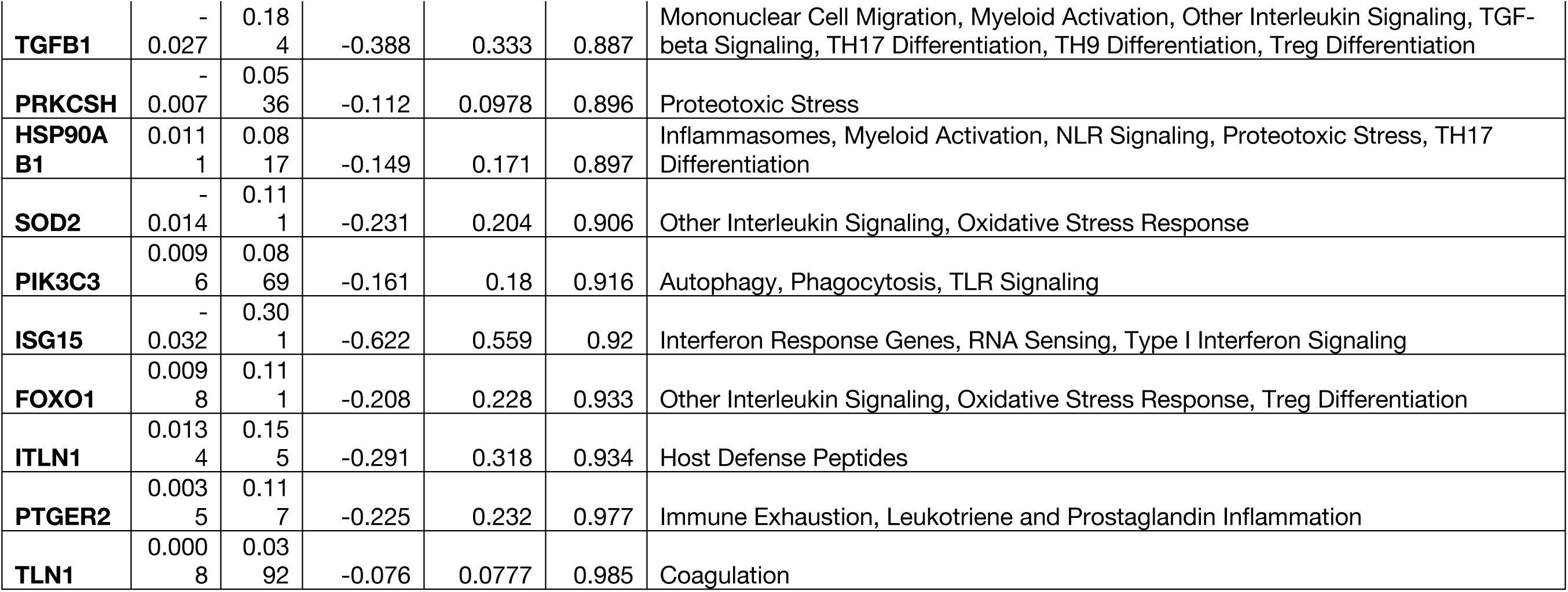
Differentially expressed genes of Caco-2 cells treated with toxins relative to no toxins

**Supplemental Table 4.**
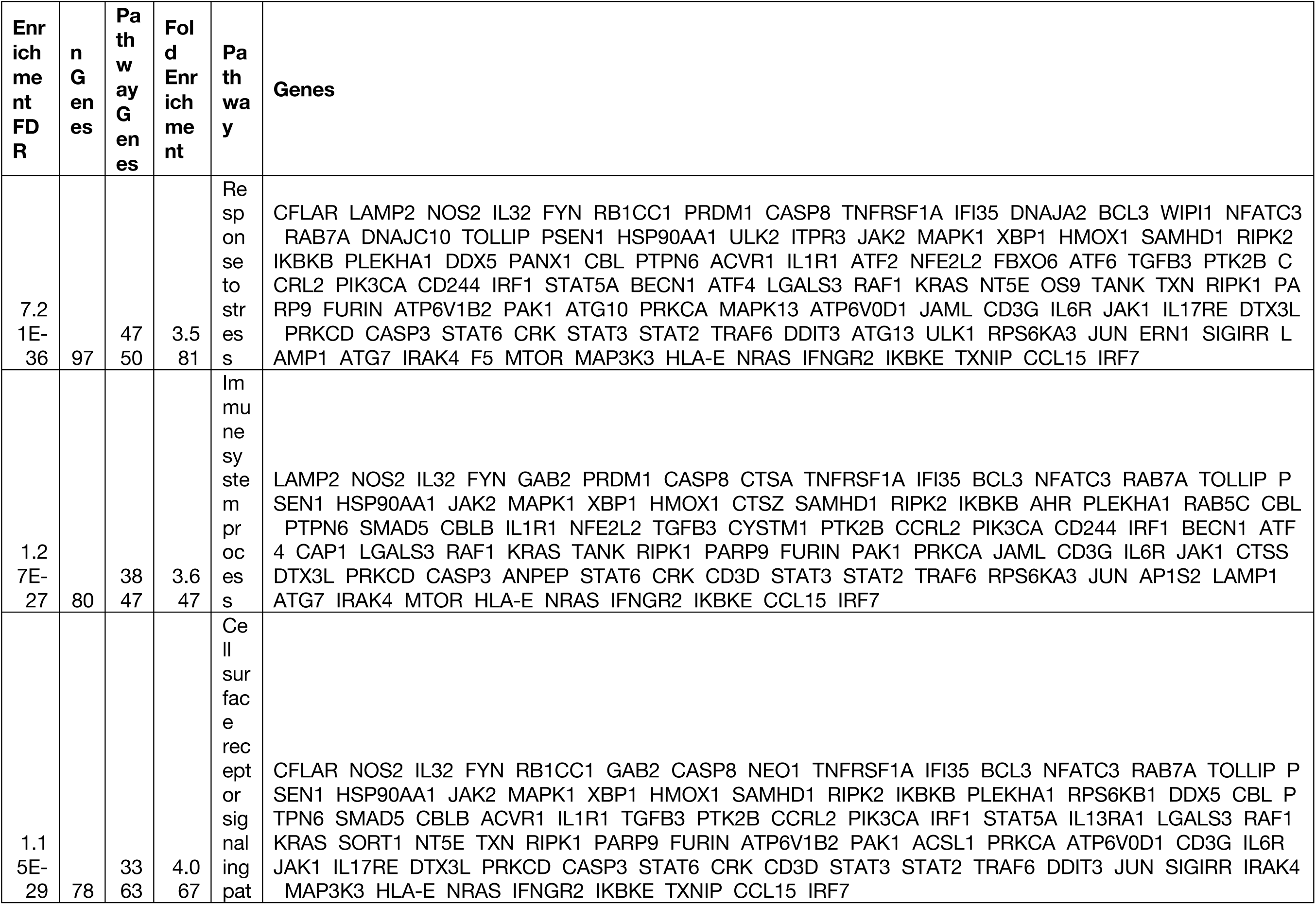
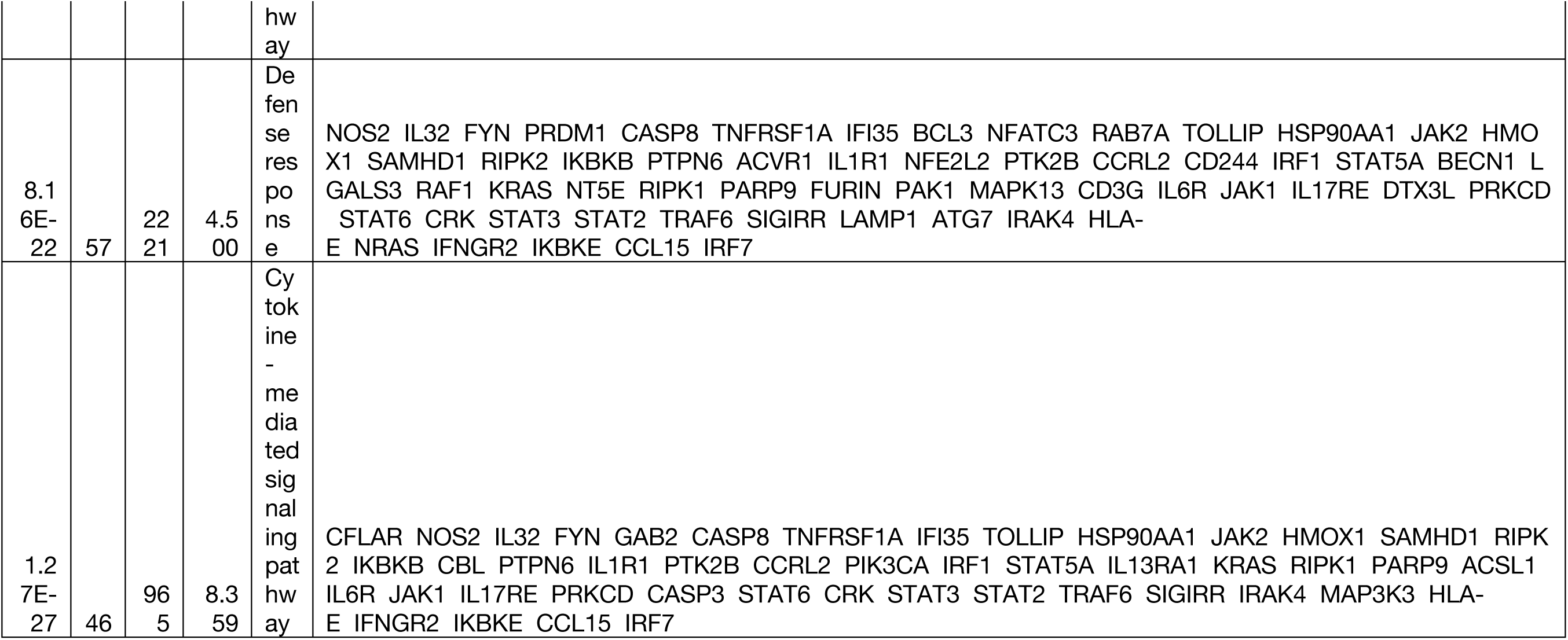
Genes in biological process GO terms from Caco-2 cells treated with toxins relative to no toxins

**Supplemental Table 5.**
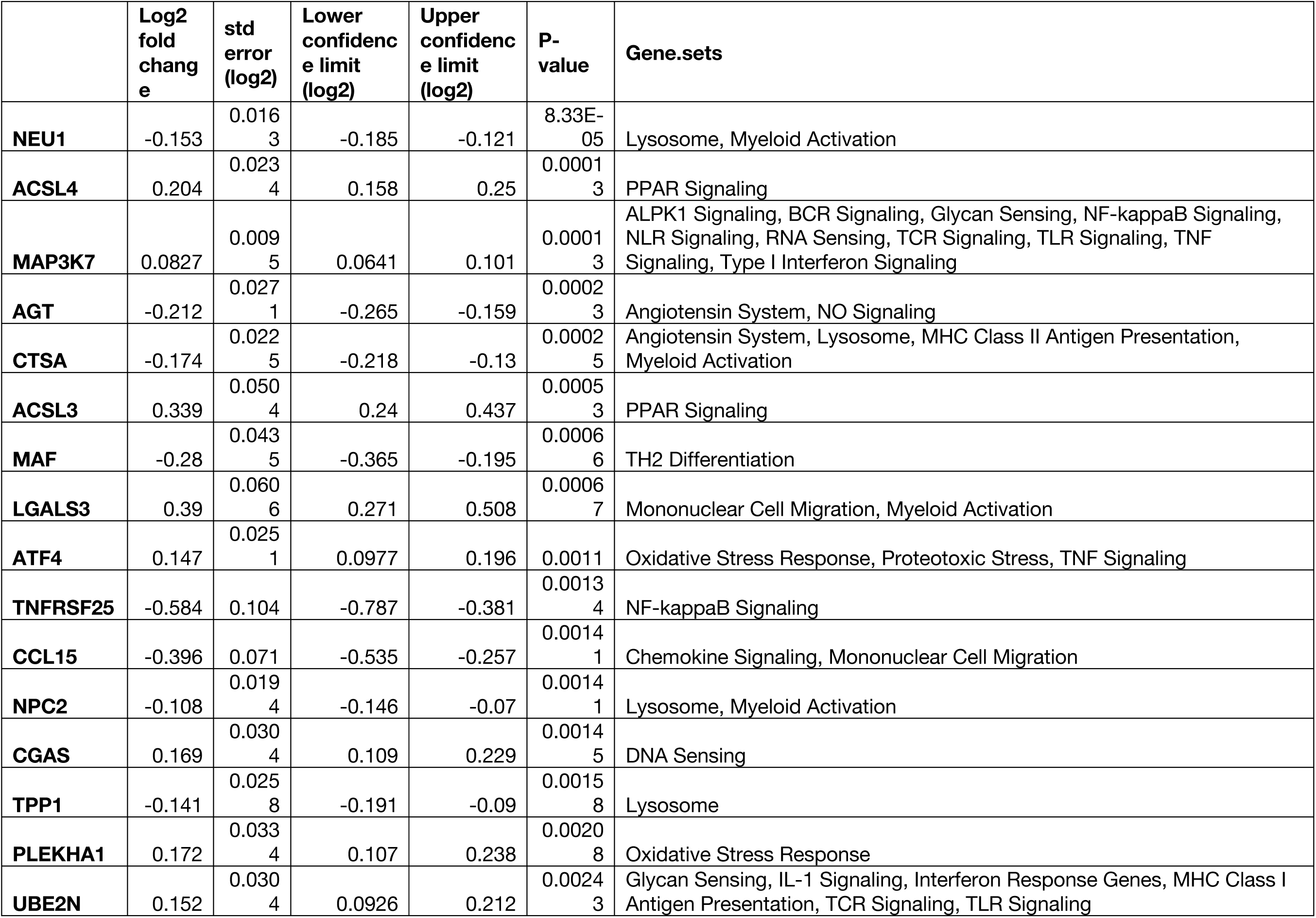
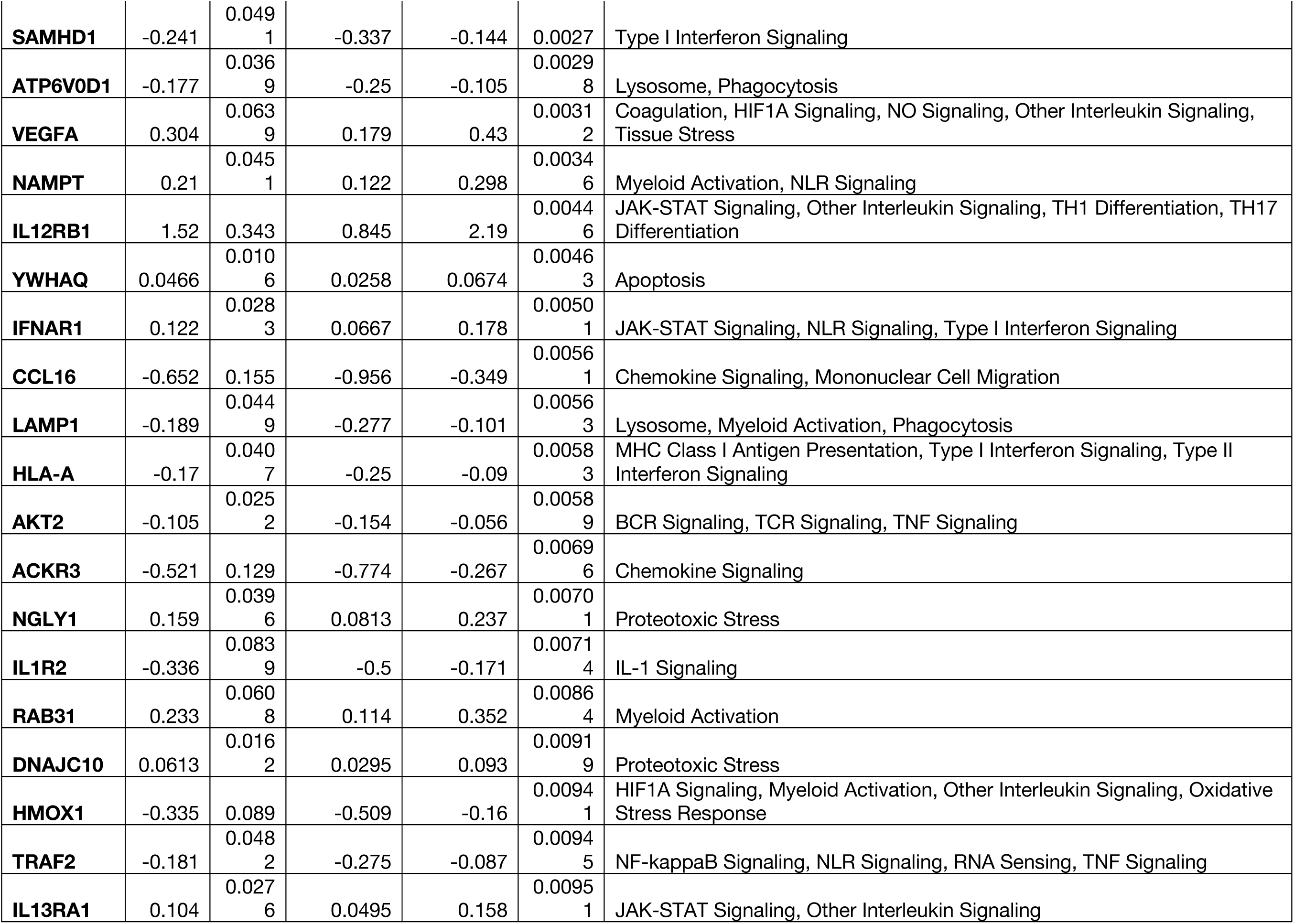
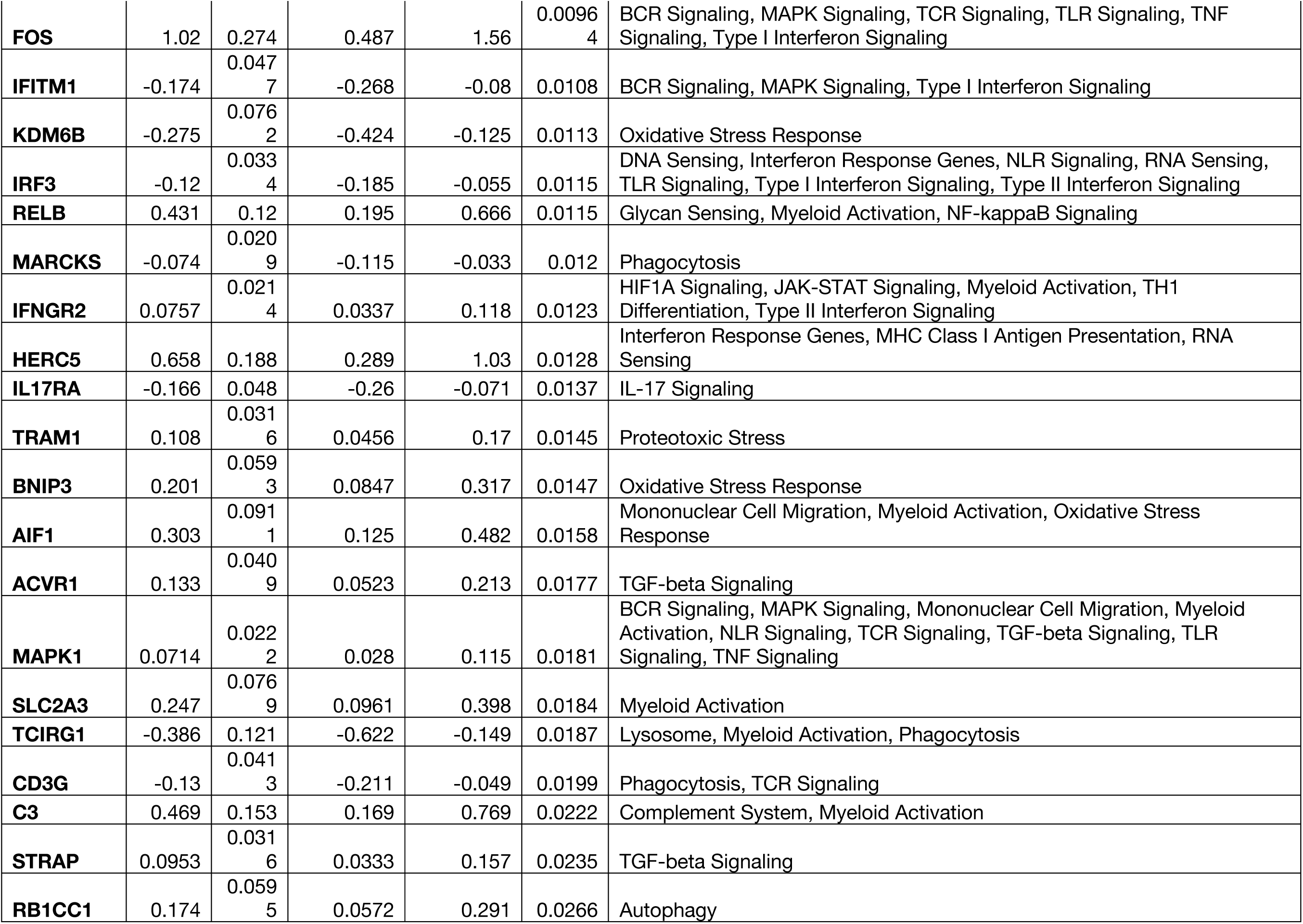
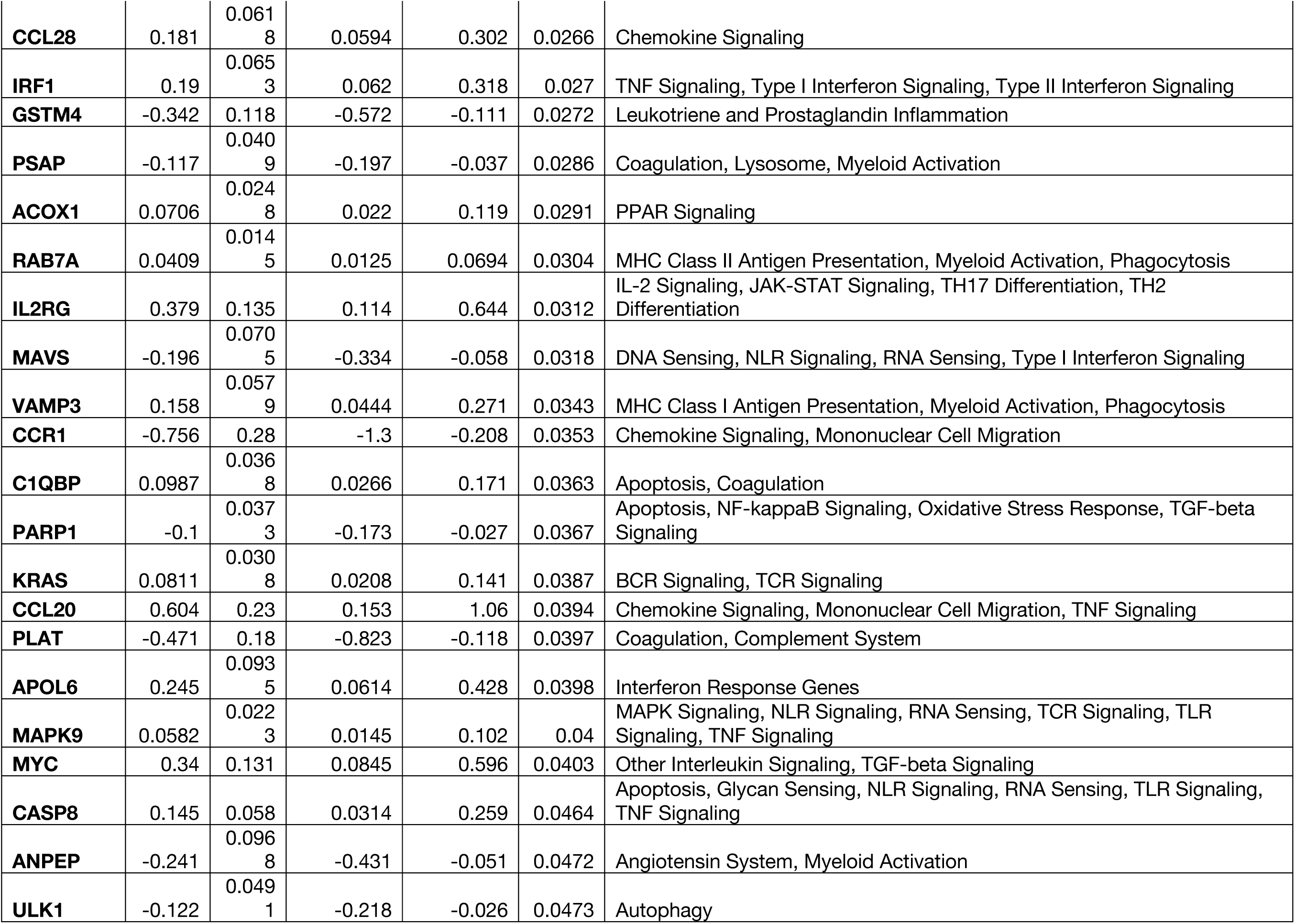
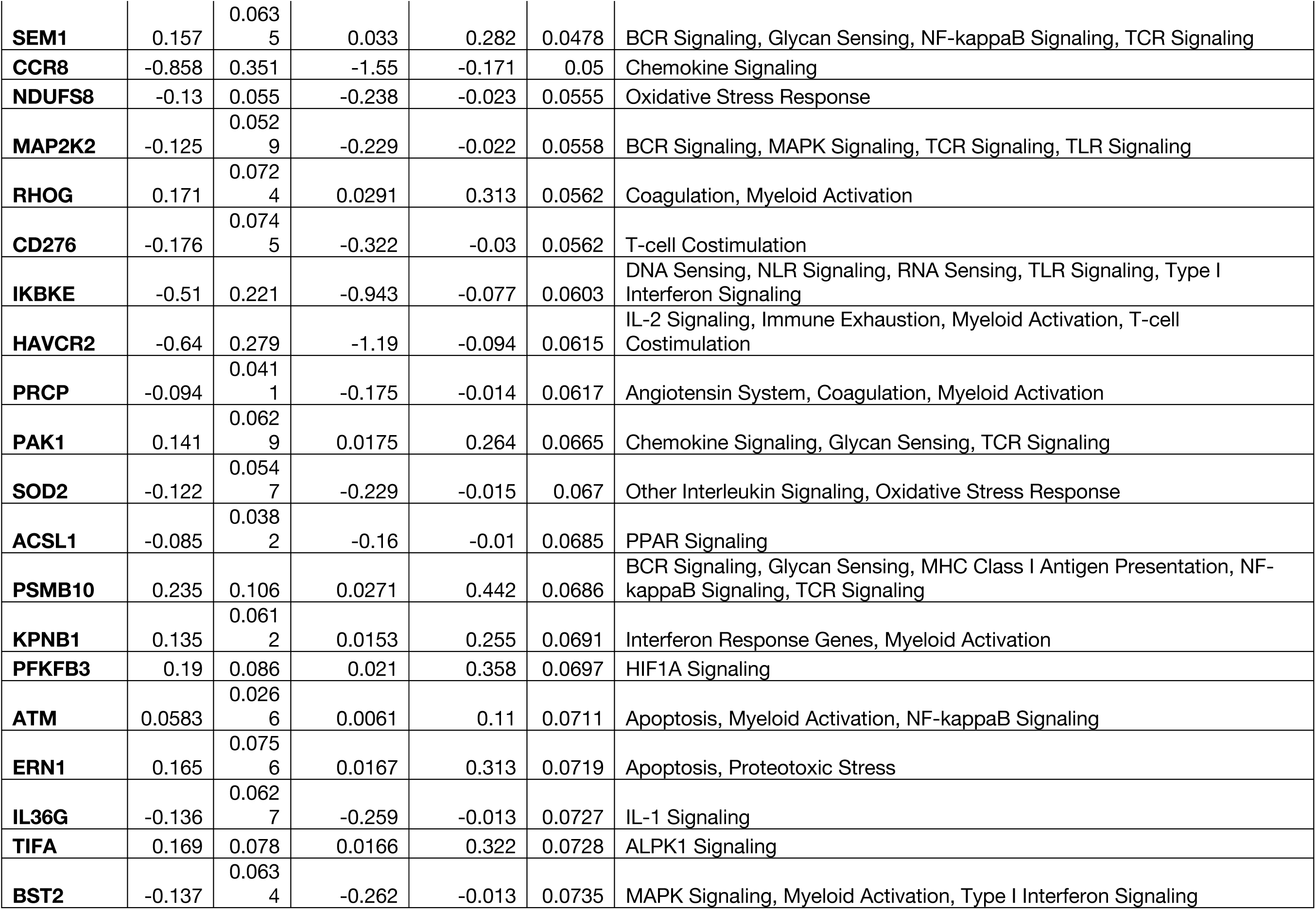
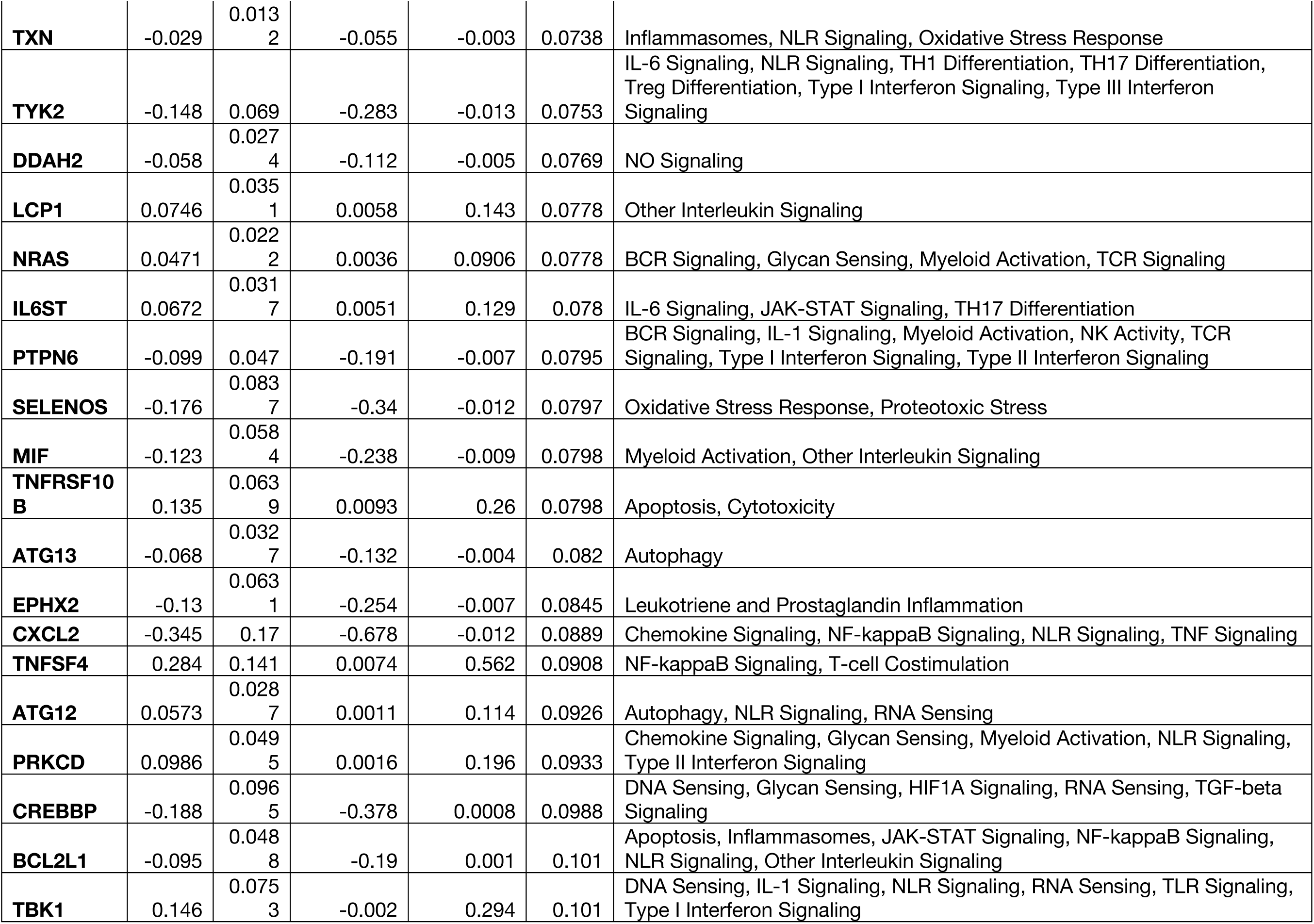
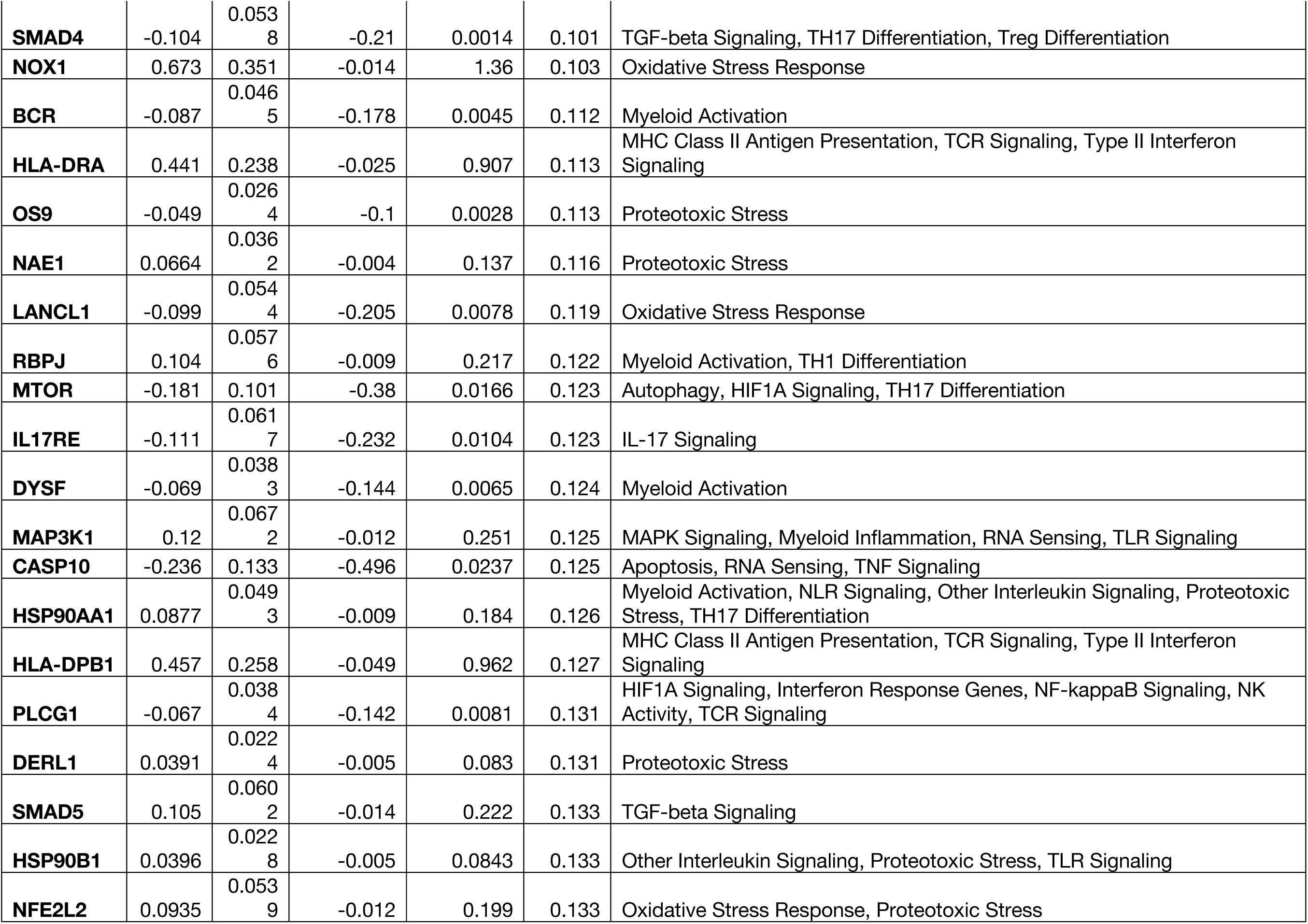
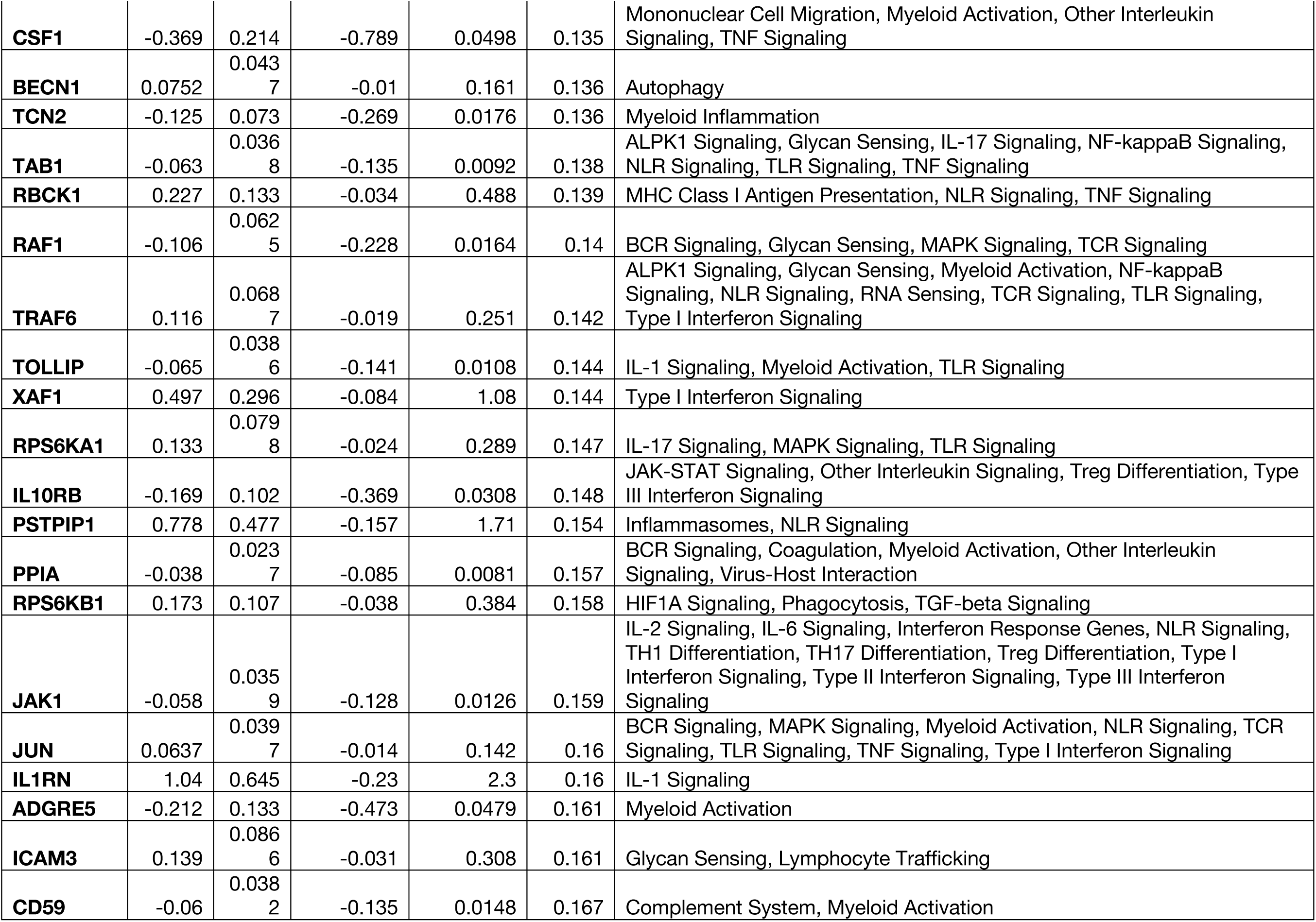
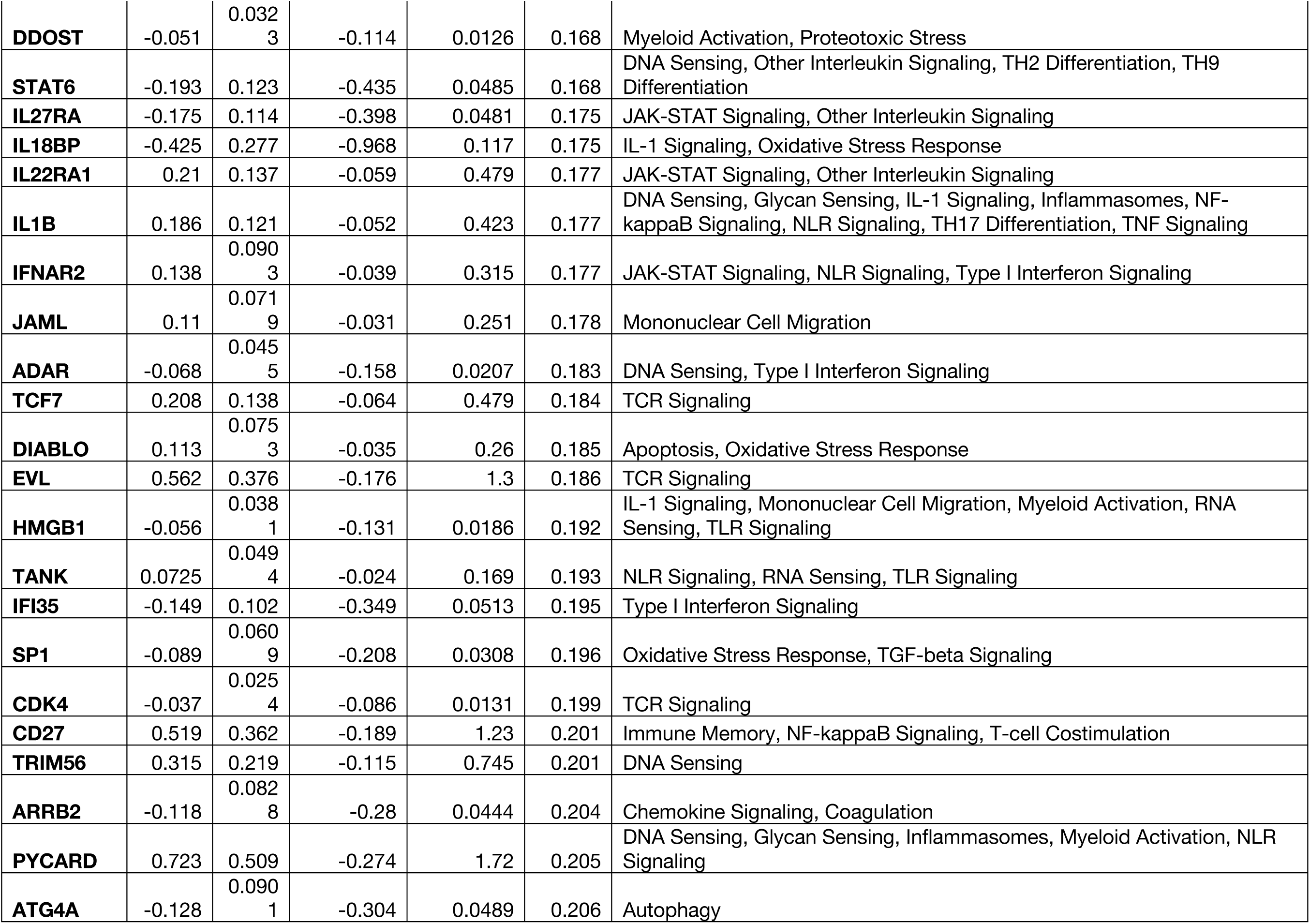
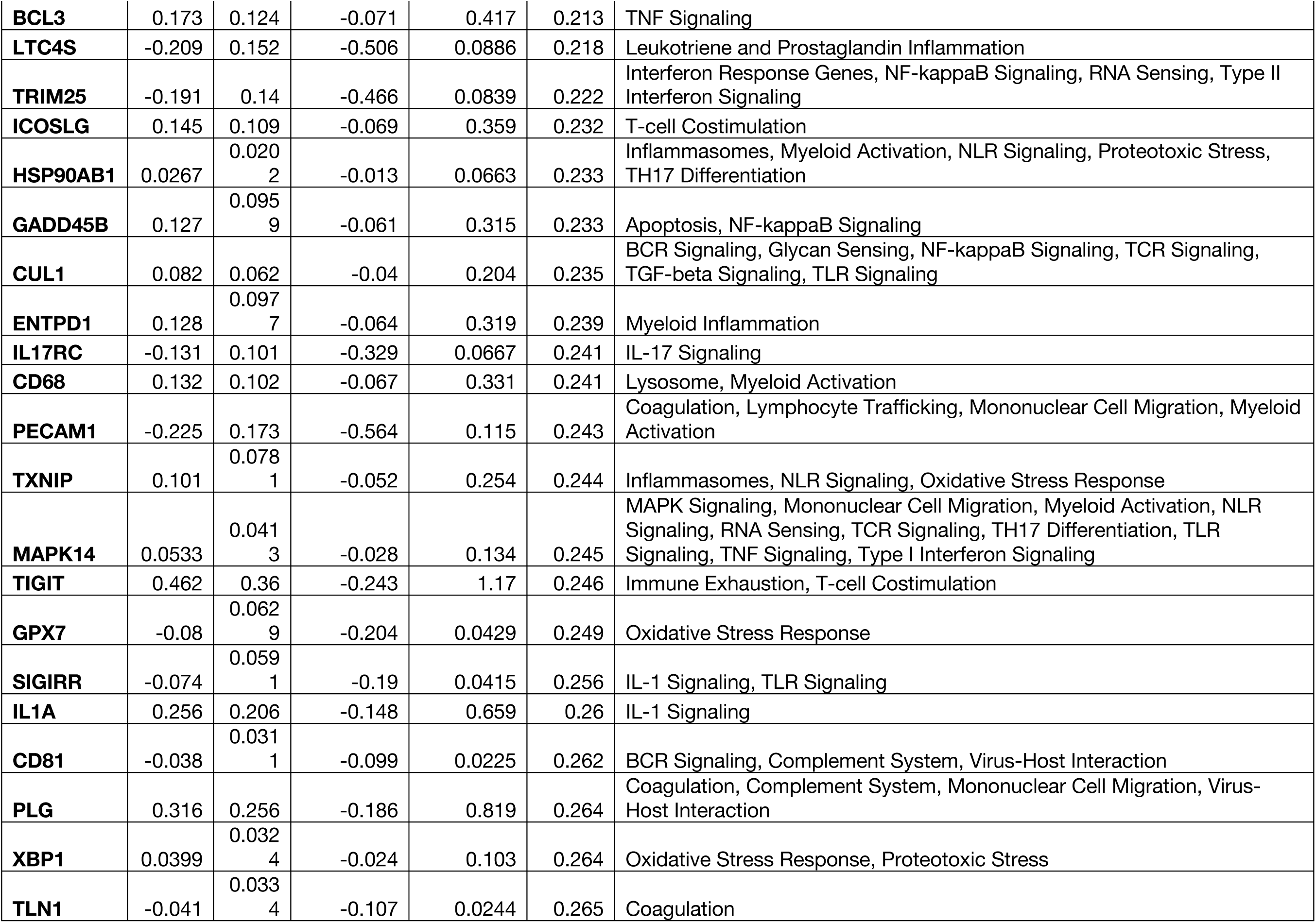
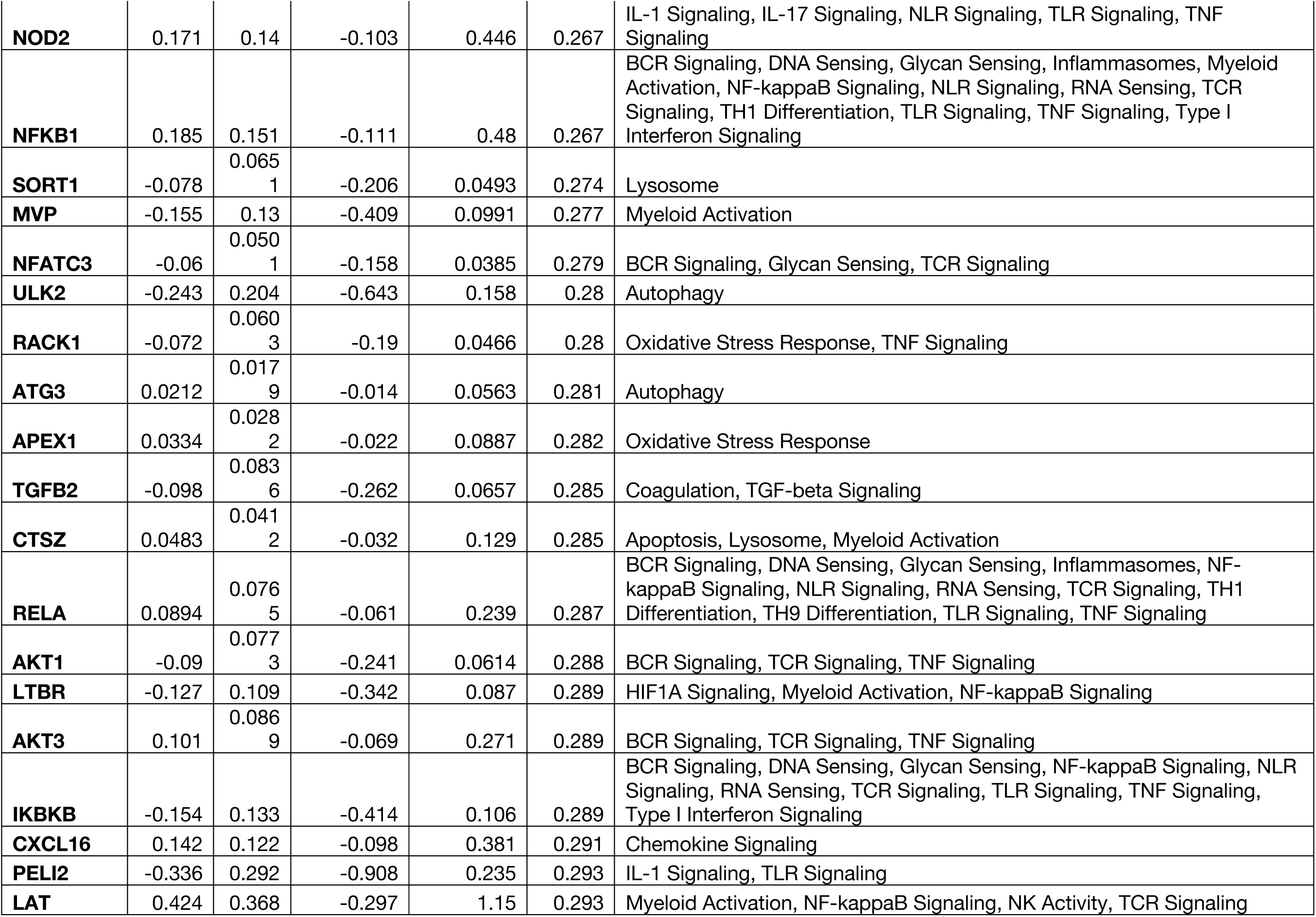
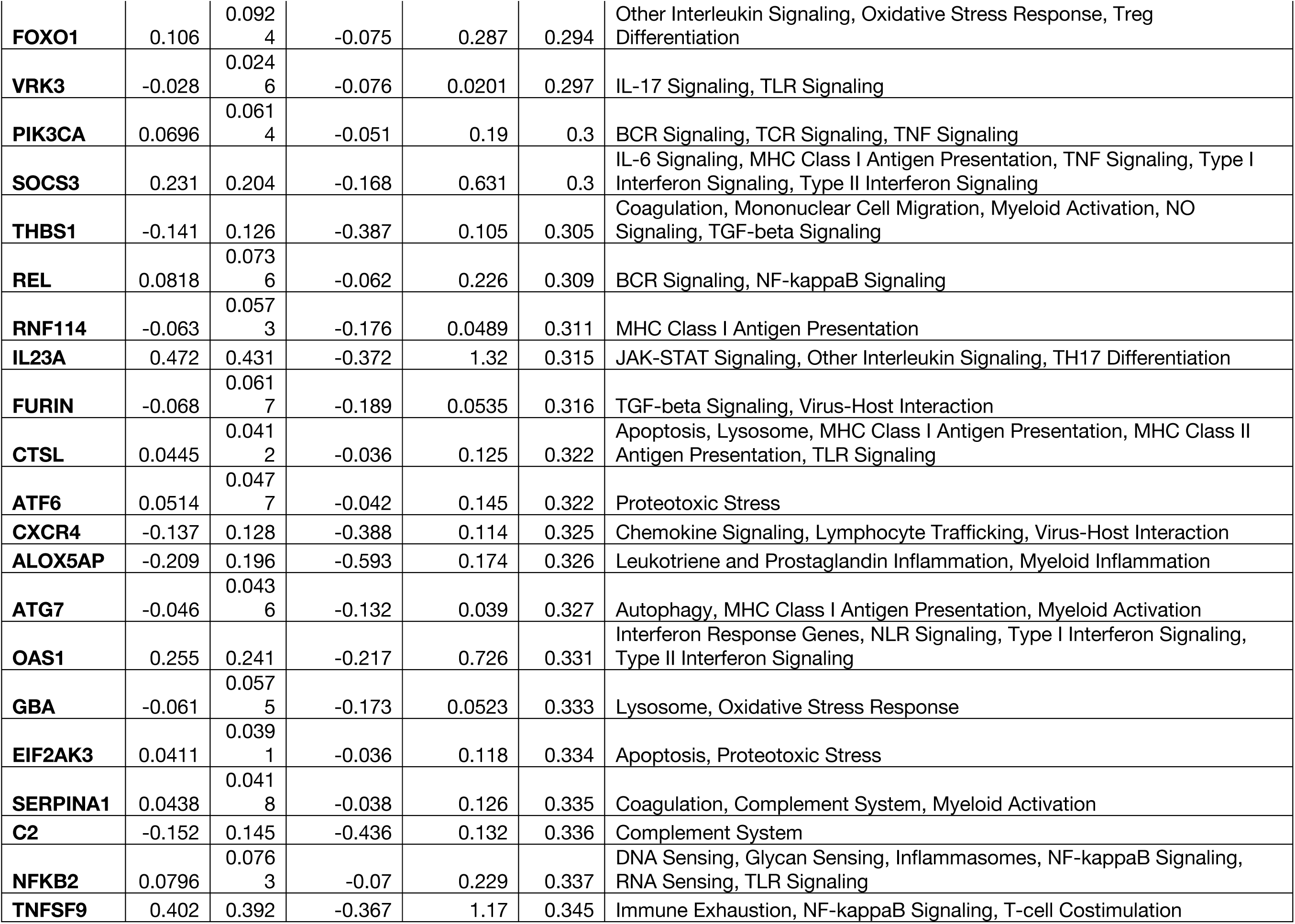
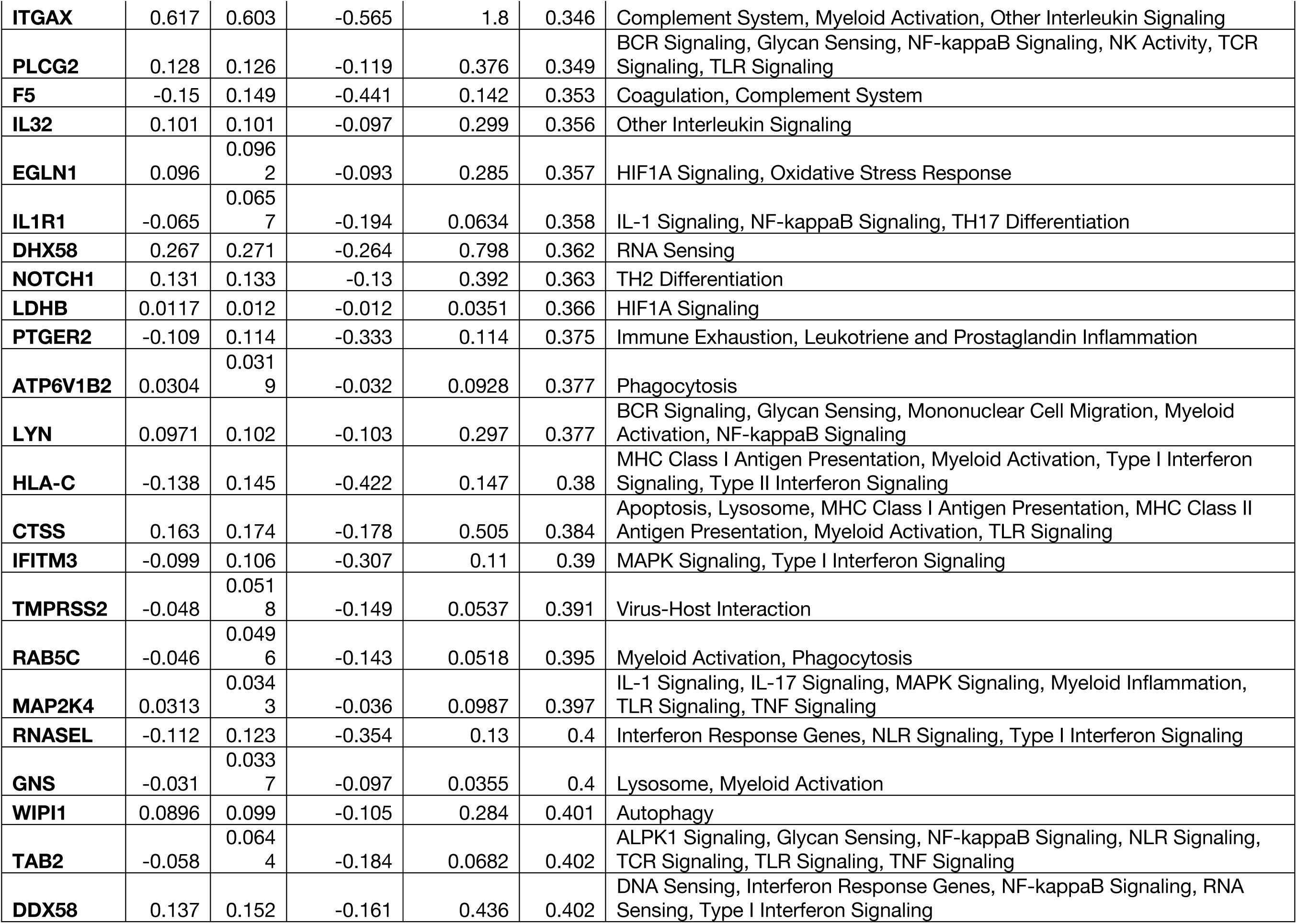
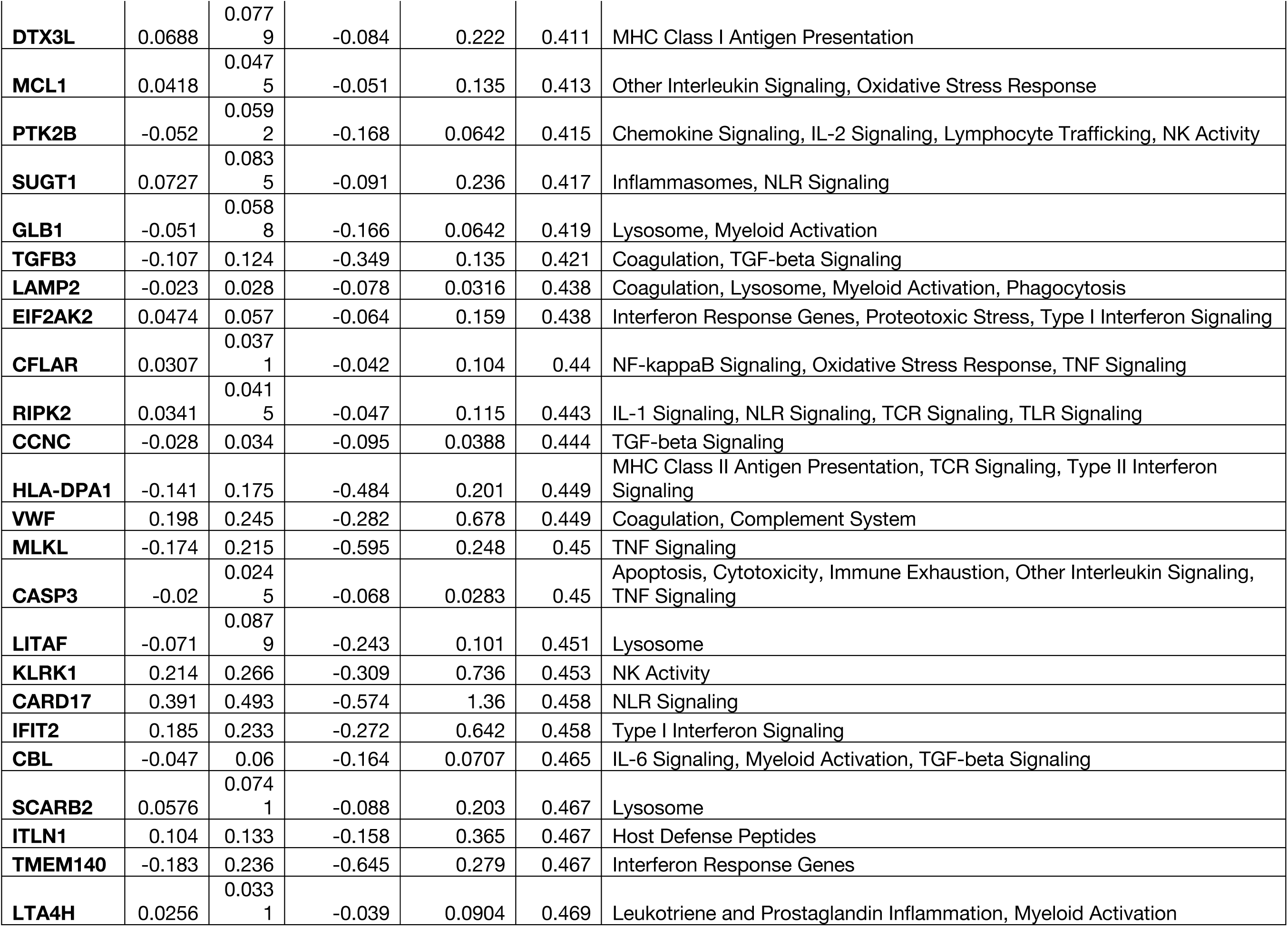
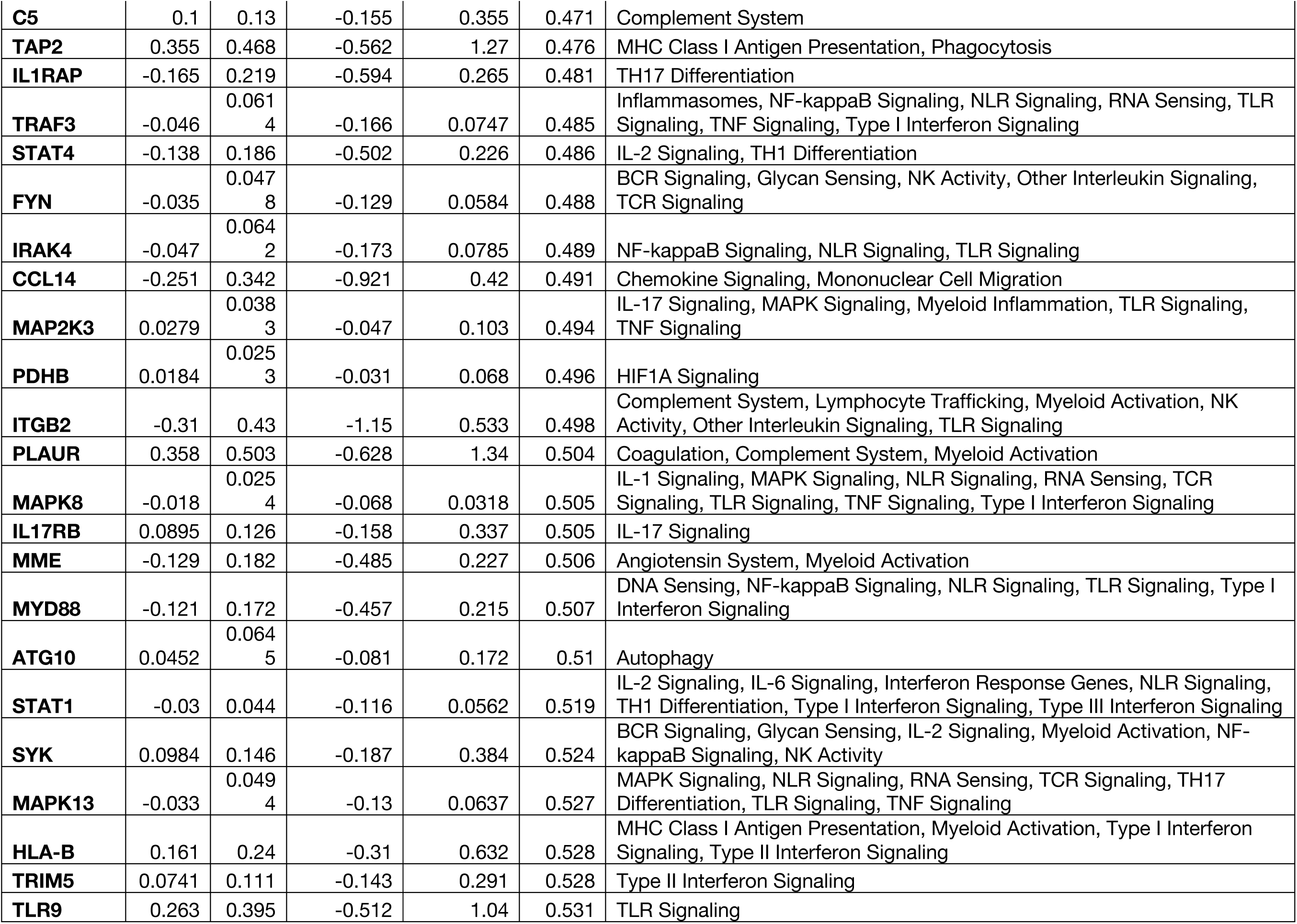
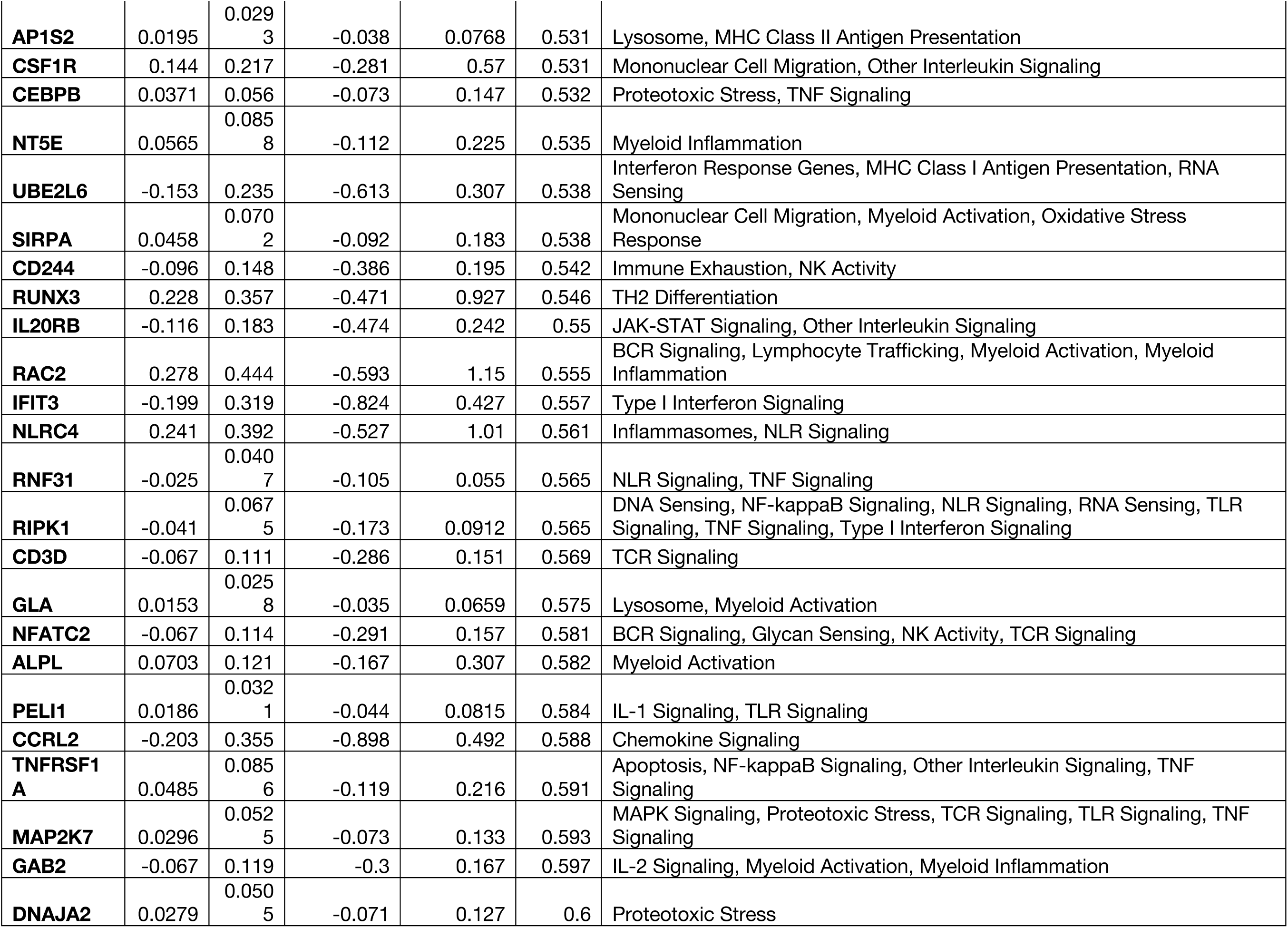
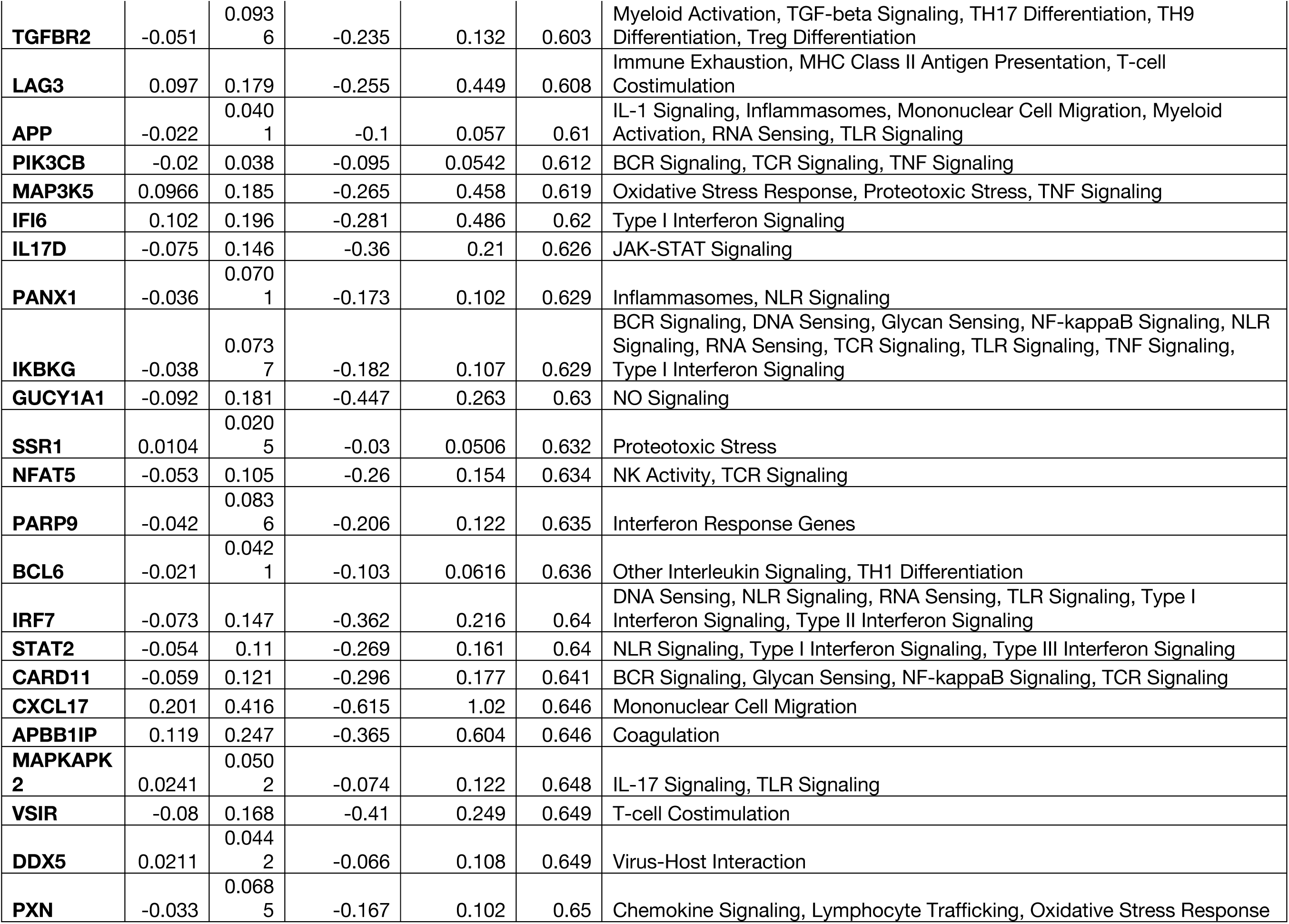
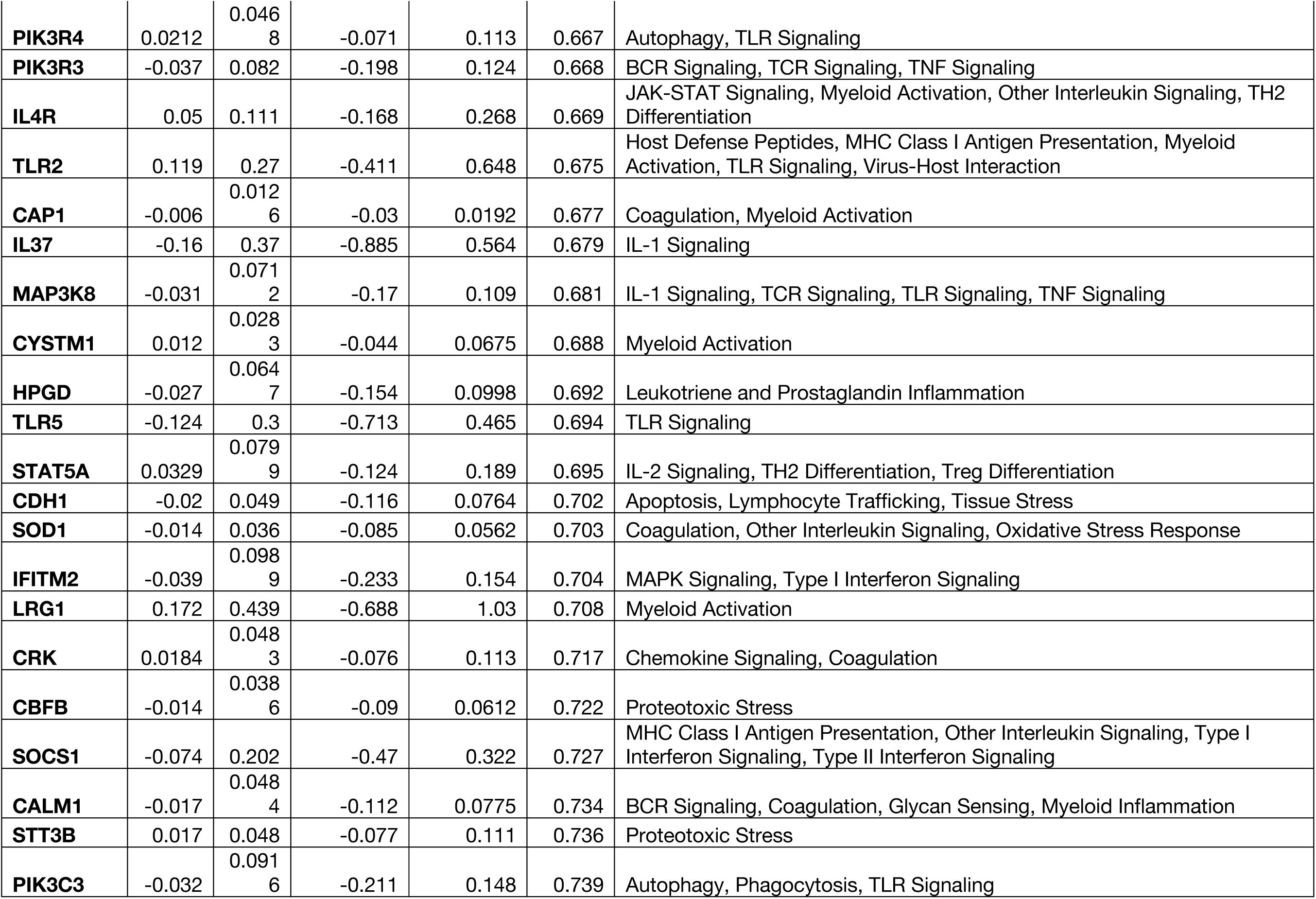
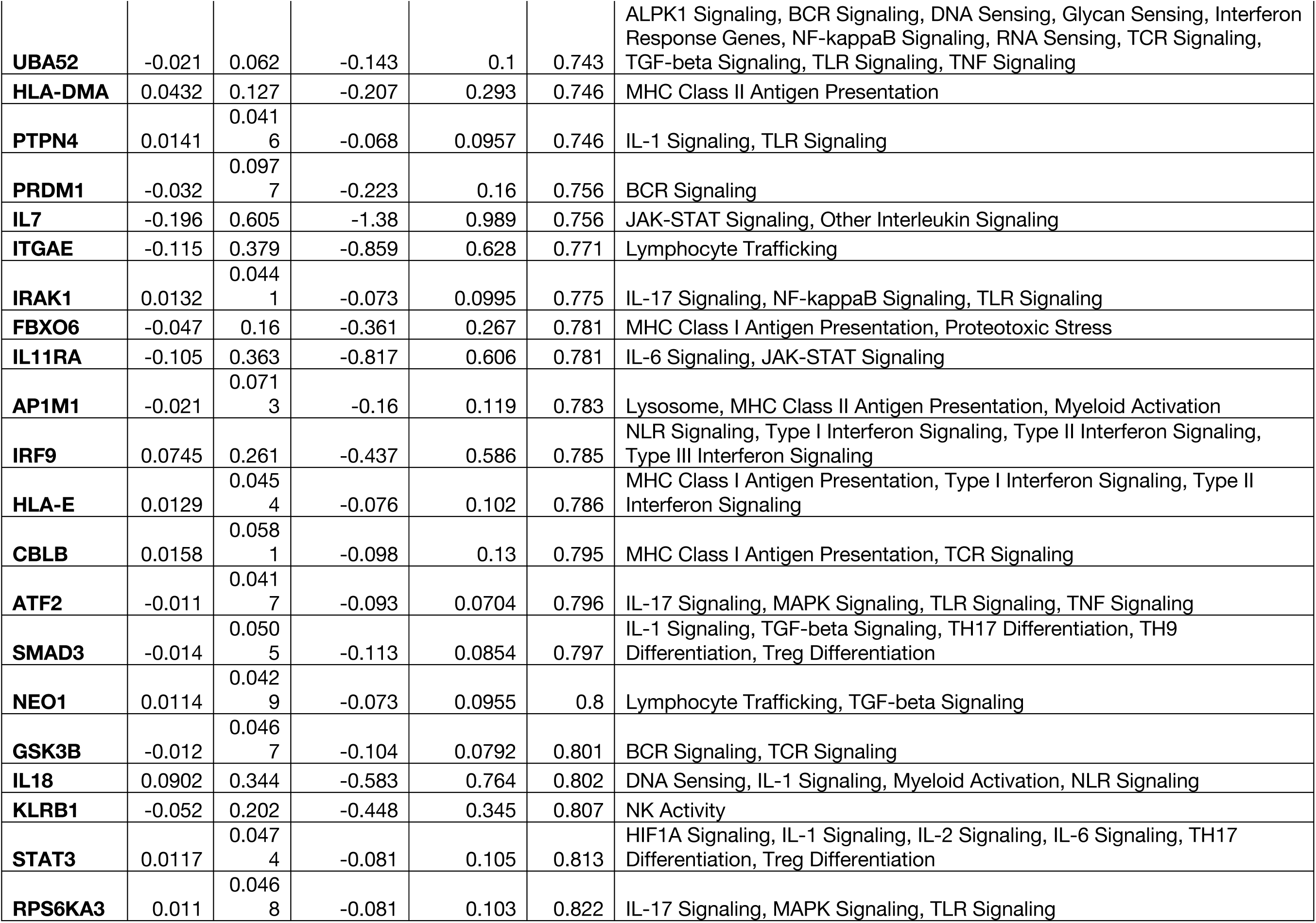
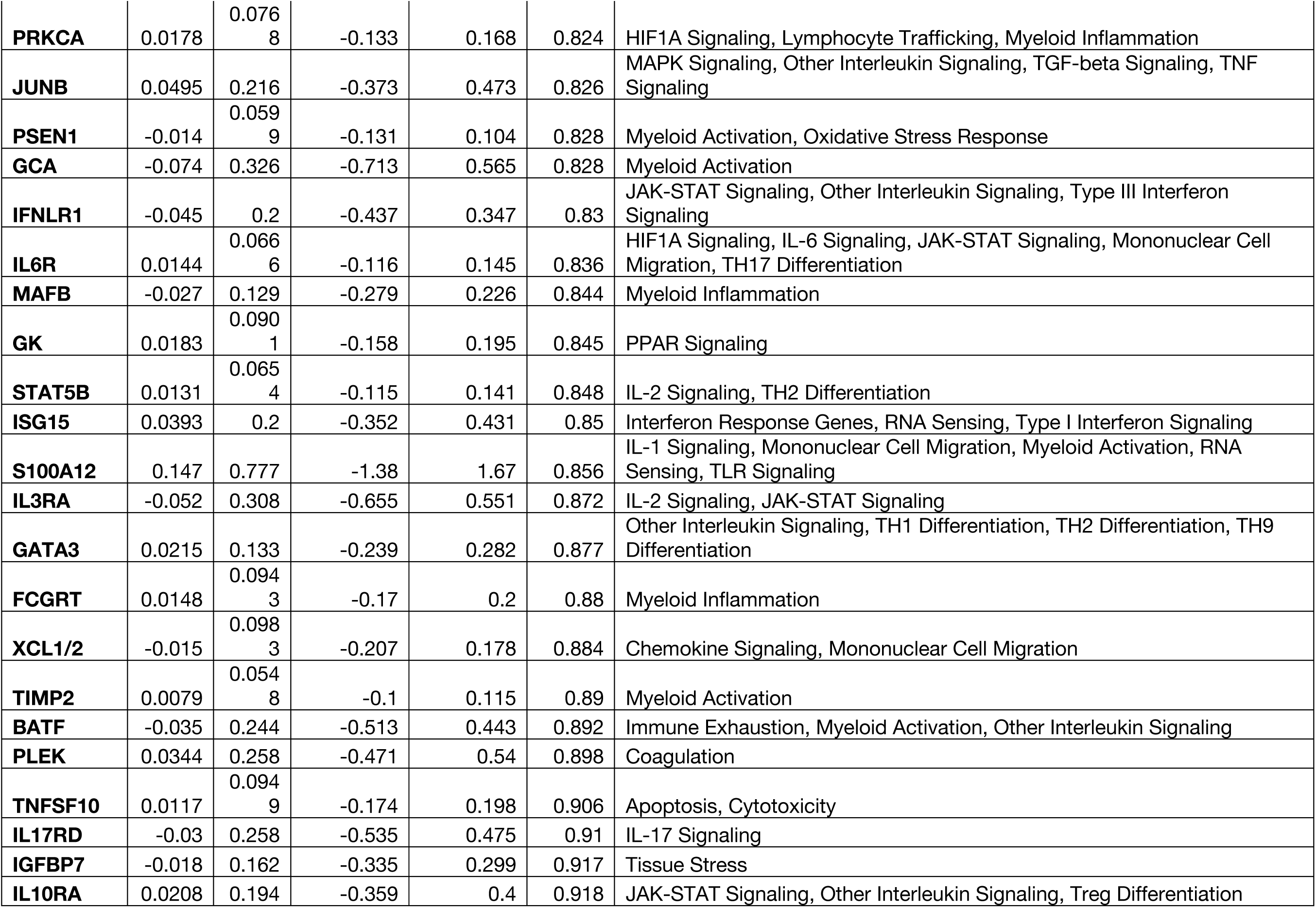
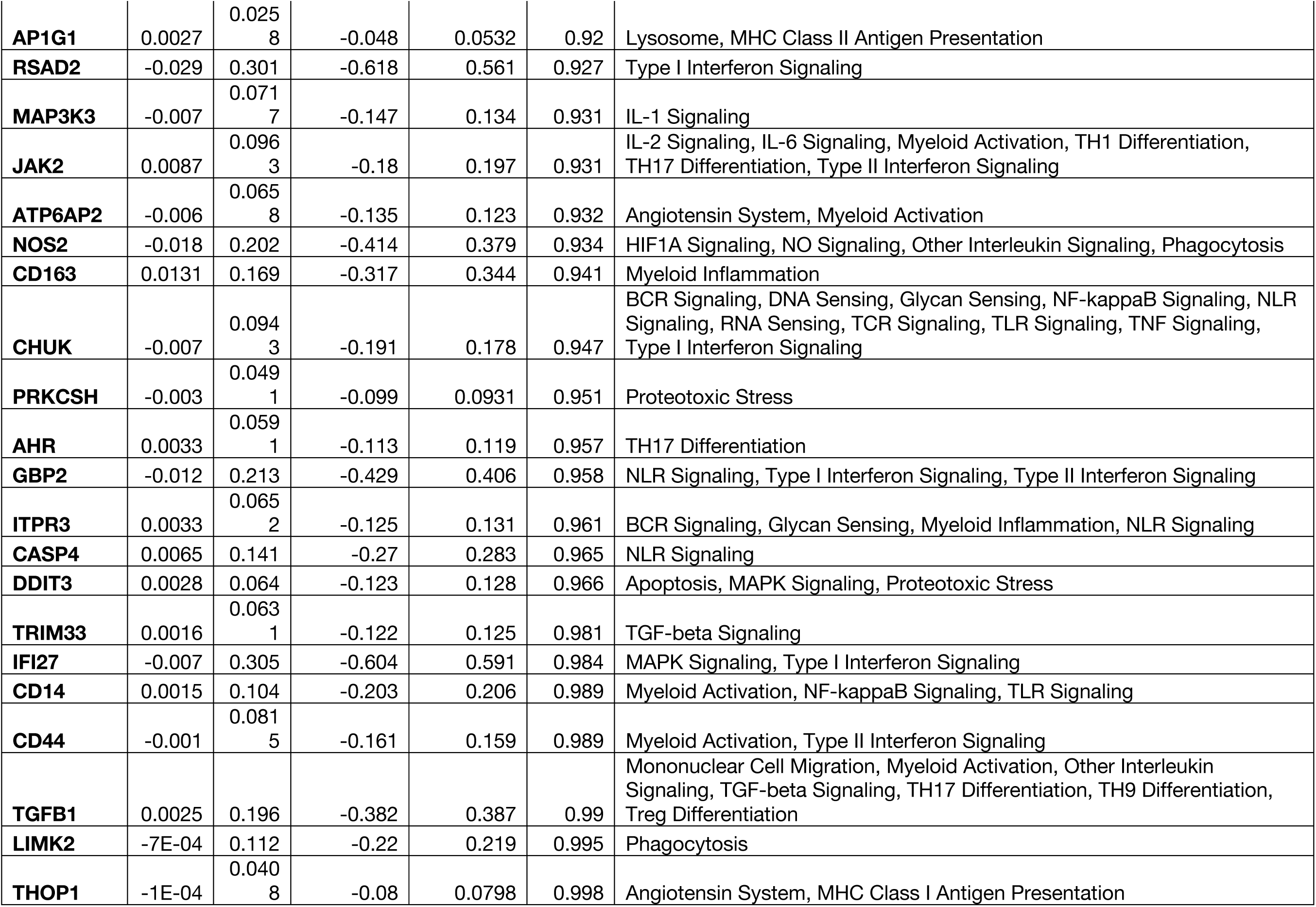
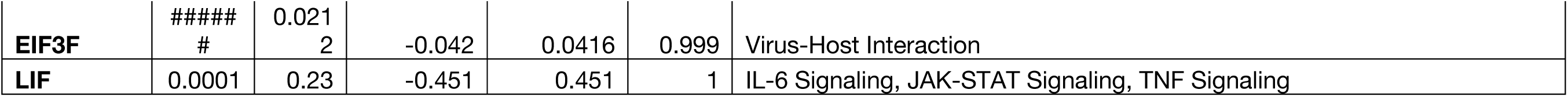
Differentially expressed genes of Caco-2 cells treated with TUDCA relative to toxins

**Supplemental Table 6.** Genes in biological process GO terms from Caco-2 cells treated with TUDCA relative to toxins

| Enrichment FDR | nGenes | Pathway Genes | Fold Enrichment | Pathway | Genes |
| --- | --- | --- | --- | --- | --- |
| 2.58 E-08 | 23 | 3847 | 4.130 | Immune system process | CTSA MAVS HMOX1 SAMHD1 TCIRG1 IL1R2 NPC2 IRF3 TRAF2 KDM6B PARP1 ACKR3 CD3G CCR1 ANPEP IL17RA NEU1 IFITM1 LAMP1 PSAP HLA-A CCL16 CCL15 |
| 3.79 E-06 | 22 | 4750 | 3.199 | Response to stress | MAVS HMOX1 SAMHD1 PLAT TCIRG1 IL1R2 IRF3 TRAF2 KDM6B AGT PARP1 ACKR3 ATP6V0D1 CD3G CCR1 TPP1 ULK1 IL17RA IFITM1 LAMP1 CCL16 CCL15 |
| 6.19 E-07 | 20 | 3363 | 4.108 | Cell surface receptor signaling pathway | MAVS HMOX1 SAMHD1 PLAT AKT2 TCIRG1 IL1R2 IRF3 TRAF2 AGT PARP1 ACKR3 ATP6V0D1 CD3G CCR1 IL17RA IFITM1 TNFRSF25 CCL16 CCL15 |
| 1.89 E-05 | 15 | 2221 | 4.665 | Defense response | MAVS HMOX1 SAMHD1 TCIRG1 IL1R2 IRF3 KDM6B AGT CD3G CCR1 IL17RA IFITM1 LAMP1 CCL16 CCL15 |
| 2.26 E-07 | 13 | 965 | 9.306 | Cytokine-mediated signaling pathway | MAVS HMOX1 SAMHD1 IL1R2 IRF3 TRAF2 ACKR3 CCR1 IL17RA IFITM1 TNFRSF25 CCL16 CCL15 |

